# Rapid functional classification of cardiac genetic variants directly informs precision cardiology

**DOI:** 10.64898/2026.04.15.718512

**Authors:** Xiao-Ting Wang, Po-Ta Chen, Joshua Mayourian, Leona Ripple, Yashasvi Tharani, Tiantian Shang, Nikoleta Pavlaki, Kevin Shani, Yongjun Jang, Christopher M. Janson, Douglas Mah, Kevin K. Parker, William T. Pu, Taekjip Ha, Vassilios J. Bezzerides

## Abstract

Large-scale clinical genome sequencing yields vast numbers of variants of unknown significance (VUSs). The high frequency of VUSs and the paucity of platforms to characterize their functional impact pose significant challenges for clinical decision making. Here, we present an integrated end-to-end platform, REVi-SCOPE (Rapid evaluation of variants in single cells by optogenetics and prime editing), for characterization of the impact of VUSs on cardiac physiology. Our strategy consists of (1) introduction of variants directly into wild-type (WT) human induced pluripotent stem cell-derived cardiomyocytes (hiPSC-CMs) via prime editing; (2) optogenetic assessment of calcium and membrane voltage dynamics in single hiPSC-CMs within the pool of edited and unedited cells; and (3) in situ single-cell genotyping of the phenotyped hiPSC-CMs with single-allele resolution. By optimizing and integrating each of these steps, we created a platform that enables VUS characterization in 10 days. We validated the REVi-SCOPE’s capabilities by analyzing the properties of established arrhythmogenic variants. We then used REVi-SCOPE to reveal the functional impact of a VUS, *TRPM4*^A320V^, identified in a child with a conduction block. Together, our results show that REVi-SCOPE enables functional characterization of VUSs linked to cardiac arrhythmias with unprecedented throughput.

## Introduction

The integration of genetic testing into cardiovascular medicine is transforming the diagnosis, risk stratification, and management of both rare and common diseases^1^. However, the growing use of clinical sequencing has uncovered numerous genetic variants, many with uncertain functional impact^2,3^. These variants of unknown significance (VUSs) dominate public databases, accounting for more than 50% of the interpreted variants in ClinVar^3^. VUSs pose diagnostic and patient management challenges, particularly when they fall within genes linked to potentially life-threatening arrhythmias such as long QT syndrome (LQTS) or catecholaminergic polymorphic ventricular tachycardia (CPVT)^2^. Standard strategies for variant interpretation rely on phenotypic concordance, familial segregation, and in silico prediction models, but the impact of many variants cannot be confidently predicted by these methods^4–7^. Experimental interrogation can provide direct insight into variant function, but established methodologies—such as variant assessment through manual electrophysiological recordings—are difficult to deploy broadly at scale, particularly for functional measurements within cardiomyocytes^2,8^. Although higher-throughput alternatives have emerged, many rely on simplified experimental contexts or sequencing-based assays that incompletely capture the physiological and functional complexity of human cardiomyocytes^9–11^.

Human induced pluripotent stem cell-derived cardiomyocytes (hiPSC-CMs) have emerged as a powerful platform for modeling inherited arrhythmias and cardiomyopathies in a human cellular context, enabling functional assessment of disease-associated variants^12–14^. Most existing approaches rely on reprogramming patient cells into patient-derived iPSCs lines or genome-editing reference iPSC lines, both of which require laborious isolation of clonal lines followed by expansion and differentiation into iPSC-CMs^14–16^. Variability between lines and between differentiations requires multiple replicates^17,18^, greatly expanding the time and resources required for rigorous results. These workflows often require months to complete and do not scale well, preventing systematic interrogation of large variant sets^14–16^. A scalable platform that surmounts these challenges and allows variants to be rapidly introduced and characterized in hiPSC-CMs would improve VUS annotation and inform the management of patients with VUSs in cardiac disease genes.

Here we present REVi-SCOPE (Rapid evaluation of variants in single cells by optogenetics and prime editing), an integrated experimental and analytical platform for scalable characterization of the impact of VUSs on hiPSC-CM calcium-handling and membrane voltage. By combining efficient population-level prime editing^19–21^, single-cell optical electrophysiology^22–24^, and coupled in situ single-cell genotyping^25,26^ with automated computational analysis, REVi-SCOPE enables rapid and systematic functional evaluation of cardiac gene variants. Together, these advances establish a foundation for more scalable and physiologically relevant interpretation of genetic variants, moving toward experimentally informed and clinically actionable variant classification.

## Results

### Develop and validate optogenetic tools for all-optical electrophysiology

To enable rapid functional assessment of VUSs in cardiac disease genes, we established an all-optical approach to quantify membrane voltage and intracellular Ca²⁺ dynamics, two processes frequently perturbed in inherited arrhythmia syndromes^27,28^. Our optimized approach combines single-cell optical pacing with red-shifted fluorescent biosensors, allowing non-invasive and scalable electrophysiological characterization of hiPSC-CMs. To improve the sensitivity and fidelity of Ca²⁺ transient measurements, we engineered a sarcoplasmic reticulum targeted Ca²⁺ sensor by fusing the red-shifted genetically encoded indicator jRCaMP1b^22^ to the sarcoplasmic reticulum (SR)-located protein junctin^29^ (Fig. 1a). This localizes the Ca^2+^ sensor near the pore of ryanodine receptor 2 (RYR2), a SR-localized intracellular Ca^2+^ channel which is often dysfunctional in arrhythmogenic disorders^30^. Adenoviral vectors encoding jRCaMP1b-Junctin and the optimized channelrhodopsin variant CheRiff^31^ were generated for delivery into hiPSC-CMs (Fig. 1b). Confocal micrographs of hiPSC-CMs transduced with adenoviruses for jRCaMP1b-Junctin and stained for cardiac Troponin T (cTnT) demonstrated effective expression of jRCaMP1b-Junctin in hiPSC-CMs (Fig. 1c).

**Fig. 1.**
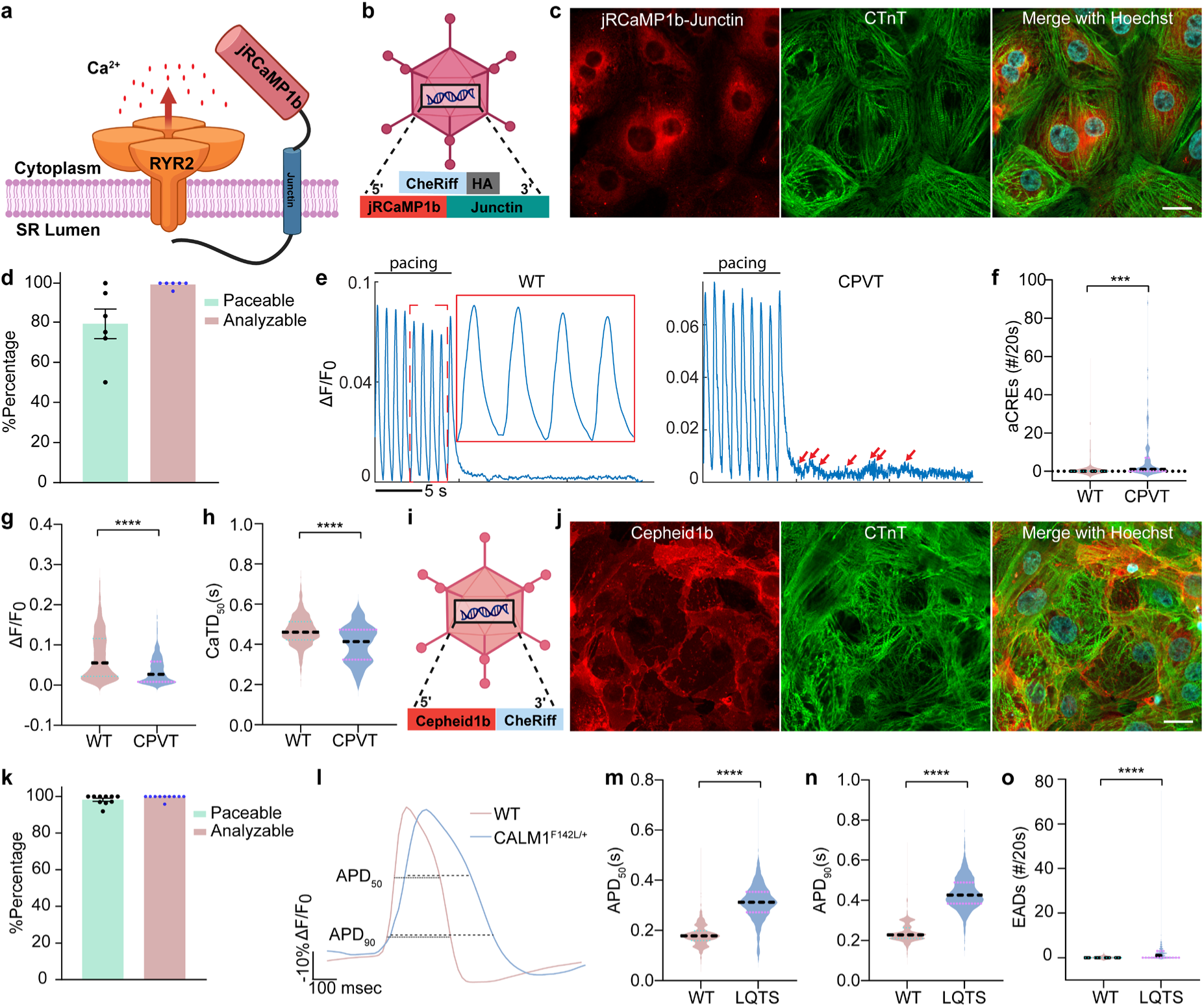
All-optical electrophysiology enables scalable functional phenotyping of hiPSC-CMs. **a**, Schematic illustrating the localization and function of the jRCaMP1b-Junctin construct. Created with BioRender.com. **b**, Schematics of AdV-jRCaMP1b-Junctin and AdV-CheRiff (AOE-MyoC) constructs. Created with BioRender.com. **c**, Representative confocal immunofluorescence images showing jRCaMP1b-Junctin expression in WT hiPSC-CMs infected with AdV-jRCaMP1b-Junctin. Cells were stained with the cardiac marker cTNT and the nuclear dye Hoechst. Scale bar = 20 μm. **d**, Percentage of paceable and analyzable WT hiPSC-CMs infected with AOE-MyoC. Mean ± SEM. **e**, Representative calcium transient traces from WT and CPVT hiPSC-CMs (*RYR2*^G3946S/+^) infected with AOE-MyoC. Red rectangle highlights an enlarged region of the waveform to illustrate trace morphology. Red arrows indicate aCREs. **f**–**h**, Quantification of aCREs (**f**), amplitude of calcium transient (**g**) and calcium transient duration at 50% recovery (CaTD_50_, **h**), from the groups in (**e**). The dotted horizontal line indicates the median. WT, *n* = 666 cells; CPVT, *n* = 240 cells. Mean ± SEM. Welch’s t-test: \*\*\**P* < 0.001, \*\*\*\**P* < 0.0001. **i**, Schematic of AdV-Cepheid1b-CheRiff (AOE-MyoV) construct. Created with BioRender.com. **j**, Representative confocal immunofluorescence images showing Cepheid1b expression in WT hiPSC-CMs infected with AOE-MyoV. Cells were stained with cTNT and Hoechst. Scale bar = 20 μm. **k**, Percentage of paceable and analyzable WT hiPSC-CMs infected with AOE-MyoV. Mean ± SEM. **l**, Optical recordings of action potentials from WT and LQTS (*CALM1*^F142L/+^) hiPSC-CMs infected with AOE-MyoV. **m**–**o**, Quantification of APD at 50% recovery (APD_50_, **m**), APD at 90% recovery (APD_90_, **n**) and EADs (**o**) from the groups shown in (**l**). The dotted horizontal line indicates the median. WT, *n* = 1165 cells; LQTS, *n* = 2160 cells. Mean ± SEM. Statistical significance was determined using Welch’s t-test. \*\*\*\**P* < 0.0001.

hiPSC-CMs harboring variants in *RYR2* that cause CPVT exhibit aberrant Ca^2+^ release after cessation of pacing^32^. As a proof-of-concept, we co-transduced jRCaMP1b-Junctin and CheRiff into WT or *RYR2*^G3946S/+^ hiPSC-CMs (Supplementary Fig. 1a), a variant known to cause CPVT^32^. A high proportion of cells in the assay could be paced and analyzed (Fig. 1d). Upon termination of 1-Hz optical pacing, we captured frequent spontaneous abnormal Ca^2+^ release events (aCREs) in *RYR2*^G3946S/+^ cells using an environmentally controlled multi-well imaging system equipped for optical pacing and high-speed fluorescent imaging, referred to as a kinetic imaging cytometer (KIC) (Fig. 1e and f). In addition, Ca^2+^ transients in CPVT hiPSC-CMs had lower amplitude (Fig. 1g) and shorter duration (Fig. 1h). Together, these results confirmed the robustness and sensitivity of the optogenetic Ca²⁺ imaging workflow.

In parallel, we developed an optogenetic strategy to measure membrane depolarization and repolarization^33^. We utilized the recently developed red-shifted genetically encoded voltage indicator (GEVI) Cepheid1b^24^, which improves upon the slow kinetics, suboptimal signal-to-noise ratios, or complex imaging requirements of earlier GEVIs^24,34,35^. To transduce hiPSC-CMs, we packaged the Cepheid1b along with CheRiff into a single adenoviral construct (Fig. 1i).

Transduced hiPSC-CMs robustly expressed Cepheid1b (Fig. 1j). Nearly all hiPSC-CMs captured could be paced and analyzed (Fig. 1k), supporting the robust expression and signal-to-noise ratio of this system.

To validate detection of disease-relevant voltage phenotypes, we examined hiPSC-CMs carrying a pathogenic variant in calmodulin (*CALM1*^F142L/+^) (Supplementary Fig. 1b), which causes severe action potential prolongation and LQTS^36^. Under steady-state optical pacing, Cepheid1b recordings revealed significantly increased action potential duration (APD) in *CALM1*^F142L/+^ cardiomyocytes compared with WT controls (Fig. 1l–n). *CALM1*^F142L/+^ cells also had more frequent early afterdepolarizations (EADs), indicating repolarization instability (Fig. 1o).

Together, these results demonstrate that this all-optical electrophysiology platform enables optical stimulation and quantitative measurement of intracellular Ca²⁺ dynamics and membrane voltage in hiPSC-CMs. The throughput and single-cell resolution of this system provide a scalable framework for functional characterization of cardiac gene variants. We refer to this system as **A**ll **O**ptical **E**lectrophysiology in **Myo**cytes for either Ca^2+^ dynamics (**AOE-MyoC**) or membrane voltage (**AOE-MyoV**).

### An end-to-end automated analysis program for high-speed functional imaging of isolated cardiomyocytes

High-speed Ca^2+^ or membrane voltage imaging remains one of the most commonly employed methods to ascertain the anti-arrhythmic or pro-arrhythmic effects of small molecules or genetic modifications in isolated cardiomyocytes, including hiPSC-CMs^37,38^. However, traditional methods for high-throughput Ca^2+^ or membrane voltage imaging with single-cell resolution pose analysis challenges, including accurate cell segmentation and low-complexity waveform processing^39–41^. To overcome these challenges, we developed CaT-Scan (Cardiomyocyte automated Tracking for Segmentation and Calcium Analysis) as an end-to-end pipeline to rapidly and accurately analyze high-speed Ca^2+^ or membrane voltage imaging data of hiPSC-CMs. We used KIC imaging to acquire high-speed (≥90 frames/sec) movies of hiPSC-CMs expressing either AOE-MyoC or AOE-MyoV. After down-sampling to reduce computational and storage demands, cell segmentation was performed using an algorithm based on Cellpose^42^ (Fig. 2a). For each cell, the pipeline then analyzed the Ca^2+^ or membrane voltage signal. It first denoised the data, corrected for photo-bleaching, and identified paced cells (Fig. 2b). For each paced cell, it analyzed the filtered Ca^2+^ transient or action potential waveform and extracted physiological parameters including upstroke and downstroke velocity, amplitude, and duration (Fig. 2b). These data were then auto-collated for further statistical analysis.

**Fig. 2.**
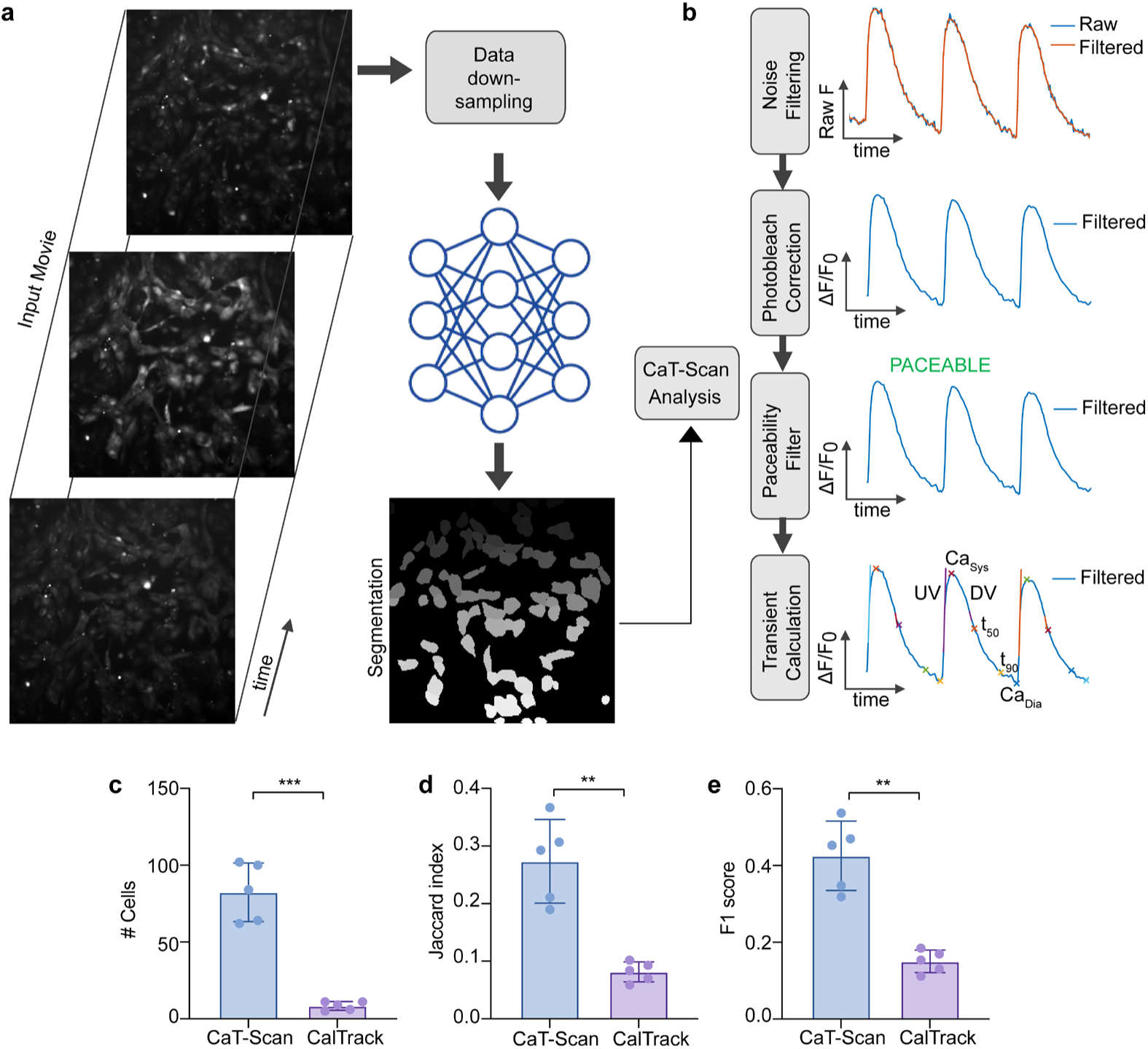
Quantitative extraction of single-cell phenotypes enables high-throughput variant assessment. CaT-Scan is an end-to-end pipeline for automated high-speed imaging analysis and function parameter extraction from fluorescent dyes and biosensors. **a**, High-speed imaging data as a movie is down-sampled, followed by automated cell segmentation through a machine-learning network. Mean fluorescence levels are measured from individual cellular masks for downstream analysis and functional parameter extraction. **b**, Functional imaging data during steady-rate pacing is subjected to multiple processing steps including noise filtering, photo-bleach correction, paceable cell identification and calcium transient or action potential parameter extraction. **c**–**e**, CaT-Scan was benchmarked against the previously published calcium imaging analysis algorithm, CalTrack. Performance relative to a human expert (gold standard) was evaluated by the number of cells identified (**c**), the Jaccard index (**d**), and the F1 score (**e**). Data are presented as mean ± SEM. Statistical comparisons were performed using Welch’s t-test. \*\**P*<0.01, \*\*\**P*<0.001.

We tested CaT-Scan against a previously published algorithm for automated Ca^2+^ imaging analysis, CalTrack^41^. When compared to gold-standard human manual segmentation, CaT-Scan successfully identified and segmented more cells, with higher Jaccard Index and F1 scores than the CalTrack benchmark (Fig. 2c–e). This resulted in a significantly higher number of analyzed hiPSC-CMs. Overall, these data demonstrate that our automated pipeline accurately captures single-cell Ca^2+^ transient and action potential parameters for phenotypic characterization at the single-cell level.

### Efficient prime editing in hiPSC-CMs

A significant barrier to the functional testing of cardiac gene variants in hiPSC-CMs is the prerequisite of clonal line generation. Additional barriers include the low yield of standard monolayer differentiation methods, batch-to-batch variability, and relative immaturity of hiPSC-CMs^17,18,43,44^. To overcome these limitations we incorporated recently described methods for efficient and large-scale hiPSC-CM differentiation by suspension culture in stirred bioreactors^43^, which produces hiPSC-CMs with high yields (>150 x 10^6^ hiPSC-CMs), purity (>90% cTnT positivity), and more mature functional properties compared to monolayer hiPSC-CMs^43^, even after cryo-recovery. These properties make these bioreactor hiPSC-CMs well suited as the starting point for our variant characterization pipeline^43^. We validated the high purity of the bioreactor-derived hiPSC-CMs used in this study by flow cytometry (Supplementary Fig. 1c–g).

Prime editing enables precise insertions, deletions, and single-base substitutions in human cells without double-strand breaks or donor DNA templates, providing broad coverage of genetic variant classes with reduced off-target activity compared to Cas9-based editing^19–21^. To enable efficient delivery of the prime editing system into hiPSC-CMs, we used combined adenoviral and modified RNA delivery to introduce the prime editor PE5max^20^ and engineered prime editing guide RNA (epegRNA)^19^, respectively. Because PE5max exceeds the adenoviral packaging limit, it was divided into two vectors: AdV-PEmax, encoding PEmax, and AdV-hMLH1dn, encoding dominant-negative human MLH1 (hMLH1dn)^20^ (Fig. 3a and b). Fluorescence imaging confirmed that both adenoviruses were robustly expressed in hiPSC-CMs (Fig. 3c and d).

**Fig. 3.**
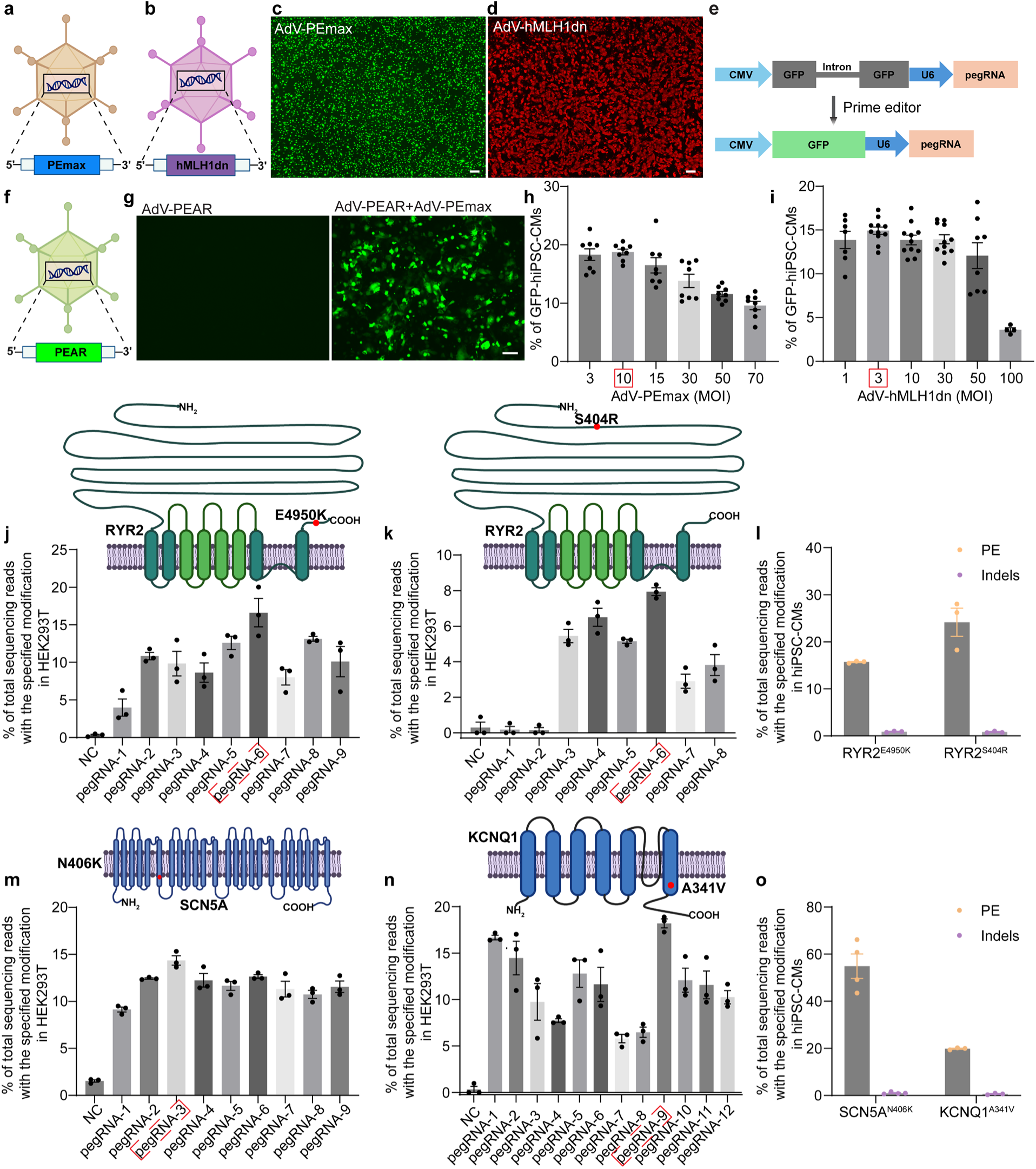
Efficient prime editing of pathogenic cardiac variants in hiPSC-CMs. **a**,**b**, Schematics of AdV-PEmax (**a**) and AdV-hMLH1dn-HA (**b**) constructs. Created with BioRender.com. **c**,**d**, Representative images showing expression of PEmax (**c**) and hMLH1dn (**d**) in WT hiPSC-CMs infected with AdV-PEmax or AdV-hMLH1dn. PEmax expression was detected by Cas9 staining (**c**), and hMLH1dn expression by HA staining (**d**). Scale bar = 200 µm. **e**, Schematic depicting the components and working principle of the PEAR (Prime Editor Activity Reporter) system. Created with BioRender.com. **f**, Schematic of AdV-PEAR construct. Created with BioRender.com. **g**, Representative images showing GFP expression in WT hiPSC-CMs infected with AdV-PEAR, with or without co-infection of AdV-PEmax. Scale bar = 100 µm. **h**,**i**, Percentage of GFP^+^ cells following infection with AdV-PEAR and varying MOIs of AdV-PEmax (**h**) or AdV-hMLH1dn-HA (**i**). Cells were sorted by FACS, and percentages were quantified using FlowJo. Red rectangles indicate the optimal working concentration. **j**,**k**,**m** and **n**, Editing efficiencies of epegRNAs targeting *RYR2*^E4950K^ (**j**), *RYR2*^S404R^ (**k**), *SCN5A*^N406K^ (**m**), and *KCNQ1*^A341V^ (**n**) in HEK293T cells, as assessed by Sanger sequencing. Red rectangles highlight epegRNAs with the highest editing efficiency. Mean ± SEM from *n* = 3 independent biological replicates. The diagram locations of variants in genes were created with BioRender.com. **l**,**o**, Editing efficiencies of CPVT variants (**l**) and LQTS variants (**o**) in WT hiPSC-CMs, as assessed by amplicon sequencing. Mean ± SEM from *n* = 3–4 independent biological replicates.

To optimize adenoviral dosing for maximal editing activity, we incorporated a Prime Editing Activity Reporter (PEAR)^45^ into an adenoviral vector (Fig. 3e and f). PEAR contains a GFP coding sequence disrupted by an inserted sequence. Successful prime editing precisely removes the inserted sequence and restores GFP fluorescence. GFP fluorescence from AdV-PEAR was only observed when hiPSC-CMs were transduced with AdV-PEmax, demonstrating that AdV-PEAR activity was contingent upon successful prime editing (Fig. 3g). Using AdV-PEAR as a functional readout, we titrated AdV-PEmax and quantified GFP^+^ cells by Fluorescence-Activated Cell Sorting (FACS), identifying the dose that maximized reporter activation (Fig. 3h and Supplementary Fig. 2a and b). We used the same approach to optimize the dose of AdV-hMLH1dn under this optimized PEmax condition (Fig. 3i and Supplementary Fig. 2a and c).

To evaluate the platform’s ability to introduce disease-associated variants into cardiac genes, we examined *RYR2*^S404R^ and *RYR2*^E4950K^, two variants linked to CPVT (Fig. 3j–l)^32,46^, and *SCN5A*^N406K^ and *KCNQ1*^A341V^, two variants linked to LQTS (Fig. 3m–o)^47,48^. Editing performance of candidate epegRNAs was first assessed in HEK293T (Fig. 3j,k,m and n). The top-performing epegRNAs were then delivered into hiPSC-CMs together with AdV-PEmax and AdV-hMLH1dn at optimized doses. Amplicon sequencing revealed precise installation of all four variants, with editing efficiencies ranging from 15.6% to 54.9%. Off-target indels within the amplicon were rare, remaining below 1.02% across all conditions (Fig. 3l and o).

### PEAR enrichment enables direct phenotypic readout of prime-edited variants

Our prime editing system introduced the *SCN5A*^N406K^ variant with more than 50% editing efficiency. We tested whether action potential prolongation caused by this variant could be detected within a population of prime-edited hiPSC-CMs using our optimized all-electrophysiology system. We treated 3D differentiated WT hiPSC-CMs with AdV-PEmax, AdV-hMLH1dn, and the *SCN5A*^N406K^ epegRNA. After prime editing, we transduced hiPSC-CMs with AOE-MyoV adenoviruses. Under 1-Hz optical pacing, the population of treated hiPSC-CMs, containing a mixture of cells with and without the variant, exhibited significantly increased APD at 50% repolarization (APD_50_) relative to control cells, consistent with its classification as a LQTS variant (Fig. 4a). Although statistically significant, overall effect size of APD prolongation was small, likely due to the mixed population of edited and unedited cells.

**Fig. 4.**
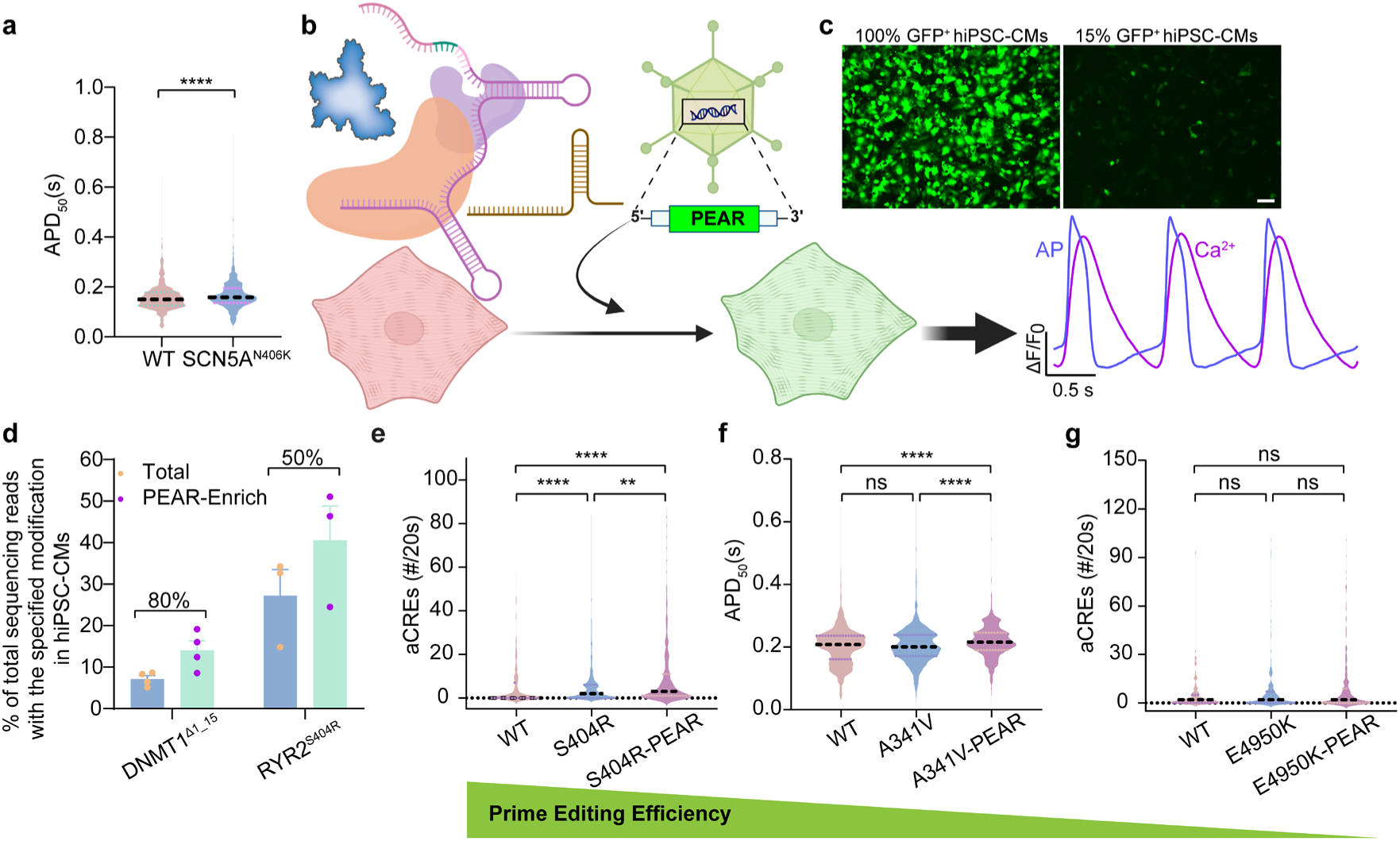
Functional classification of pathogenic variants without prior genotyping. **a**, Optical recordings of APD_50_ from WT and *SCN5A*^N406K^-edited hiPSC-CMs infected with AOE-MyoV. The dotted horizontal line indicates the median. WT, *n* = 1055 cells; *SCN5A*^N406K^, *n* = 2541 cells. Mean ± SEM. Kolmogorov-Smirnov test: \*\*\*\**P* < 0.0001. **b**, Schematic depicting overall experimental design of PEAR enrichment. Created with BioRender.com. **c**, Representative images for ∼100% and ∼15% GFP^+^ hiPSC-CMs infected with AdV-PEmax and varying MOIs of AdV-PEAR. Scale bar = 100 µm. **d**, Editing efficiencies without (total) and with PEAR enrichment, as assessed by amplicon sequencing. Mean ± SEM from *n* = 3–4 independent biological replicates. **e**, Optical recordings of aCREs from WT and *RYR2*^S404R^-edited hiPSC-CMs without and with PEAR enrichment. The dotted horizontal line indicates the median. WT, *n* = 310 cells; S404R, *n* = 708 cells; S404R-PEAR, *n* = 365 cells. Mean ± SEM. Kruskal–Wallis test: \*\**P*<0.01; \*\*\*\**P* < 0.0001; ns, *P*≥0.05. **f**, Optical recordings of APD_50_ from WT and *KCNQ1*^A341V^-edited hiPSC-CMs without and with PEAR enrichment. The dotted horizontal line indicates the median. WT, *n* = 2793 cells; A341V, *n* = 2529 cells; A341V-PEAR, *n* = 1715 cells. Mean ± SEM. Kruskal–Wallis test: \*\*\*\**P* < 0.0001; ns, *P*≥0.05. **g**, Optical recordings of aCREs from WT and *RYR2*^E4950K^-edited hiPSC-CMs without and with PEAR enrichment. The dotted horizontal line indicates the median. WT, *n* = 138 cells; E4950K, *n* = 298 cells; E4950K-PEAR, *n* = 290 cells. Mean ± SEM. Kruskal–Wallis test: ns, *P*≥0.05.

For variants with low editing efficiencies (<50%), we hypothesized that enriching cells with high prime-editing activity would increase the fraction of variant-containing cells and thereby improve phenotypic detection. We tested this hypothesis by prime editing hiPSC-CMs via co-delivery of AdV-PEAR, AdV-PEmax, AdV-hMLH1dn, and variant epegRNAs. hiPSC-CMs were subsequently treated with adenoviruses for AOE-MyoC or AOE-MyoV. Phenotyping was performed exclusively on GFP⁺ cells using optical selection (Fig. 4b). To increase the stringency of enrichment, PEAR was delivered at a low multiplicity of infection (MOI = 1), yielding approximately 15% GFP⁺ cells (Fig. 4c). We validated this enrichment strategy by performing prime editing to delete *DNMT1*^+1^ ^to^ ^15^ (*DNMT1*^Δ1_15^)^19^ or install *RYR2*^S404R^. GFP^+^ cells were isolated by FACS (Supplementary Fig. 3a–d) and editing efficiency was measured by amplicon sequencing. Inclusion of PEAR GFP selection increased the percentage of editing cells by ∼80% for *DNMT1*^Δ1_15^ and ∼50% for *RYR2*^S404R^ (Fig. 4d).

We next applied PEAR enrichment to the functional assessment of pathogenic variants spanning a range of baseline editing efficiencies. For *RYR2*^S404R^, which exhibited ∼30% editing efficiency, PEAR enrichment increased both the magnitude and statistical significance of aCREs (Fig. 4e). For *KCNQ1*^A341V^ (∼20% editing efficiency), PEAR enabled detection of a modest but significant prolongation of APD₅₀ that was not detectable in unenriched populations (Fig. 4f). In contrast, for *RYR2*^E4950K^ (∼15% editing efficiency), PEAR enrichment did not produce a detectable increase in aCREs relative to controls, suggesting that editing efficiencies below ∼20% are insufficient for functional assessment even with PEAR-mediated enrichment (Fig. 4g).

### Single-cell genotype–phenotype mapping using varGOLDFISH

Given that the mixture of edited and non-edited hiPSC-CMs impairs our ability to characterize the functional effect of cardiac gene variants, we pursued methods that would enable single-cell genotyping and thereby connect each cell’s electrophysiological properties with its genotype. To enable in situ single-cell genotyping, we integrated single guide Genome Oligopaint via Local Denaturation Fluorescence in Situ Hybridization (sgGOLDFISH)^25^. This high-sensitivity imaging technique enables single-allele resolution in genomic DNA in situ. In sgGOLDFISH, a guide RNA targeting the variant directs Cas9 to cleave the genomic locus, enabling loading of a superhelicase, which creates local DNA denaturation that enables probe binding. We implemented a dual-probe design in which the gene of interest was targeted with two fluorescent probes (labeled with Cy5 or Cy3B) one directed to the variant site (sgGOLDFISH), and a second targeting a neighboring reference region (refGOLDFISH) collectively referred to as varGOLDFISH. To minimize false-positive calls, only colocalized signals from both probes were scored as true mutant alleles (Fig. 5a).

**Fig. 5.**
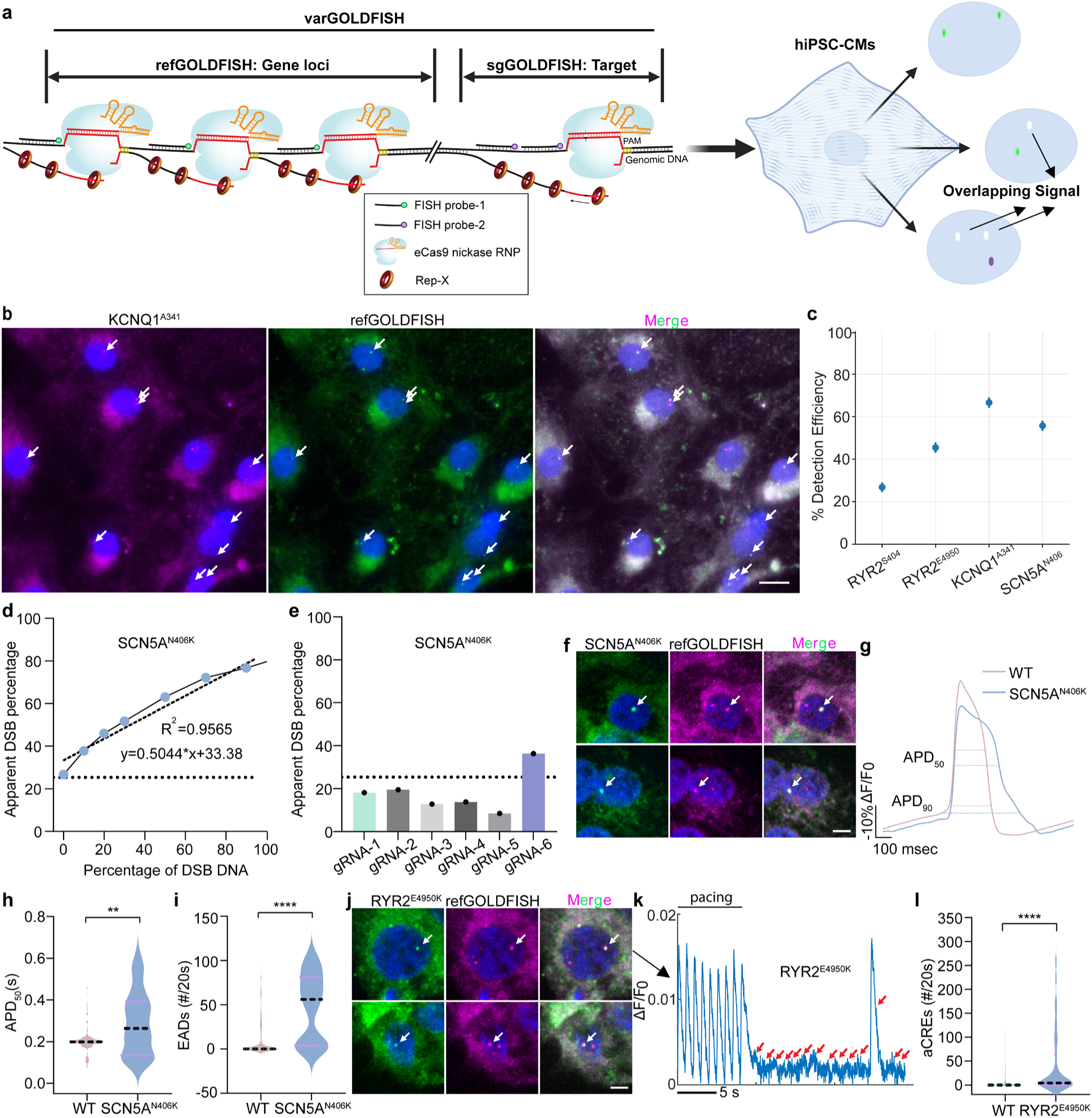
Single-cell genotyping links edited alleles to functional phenotypes. **a**, Schematic of dual-probe strategy. Modified with BioRender.com. **b**, Representative images showing refGOLDFISH and sgGOLDFISH signals for *KCNQ1*^A341^ in WT hiPSC-CMs using WT gRNA targeting the WT sequence. Scale bar = 10 µm. **c**, Detection efficiencies of sgGOLDFISH in WT hiPSC-CMs using WT gRNAs. Mean ± SD. *RYR2*^S404^, *n* = 357 cells, *RYR2*^E4950^, *n* = 421 cells, *SCN5A*^N406^, *n* = 408 cells, and *KCNQ1*^A341^, *n* = 368 cells. **d**, Standard curve of SSB-ddPCR for the *SCN5A*^N406^ locus. The dotted line indicates the true 0% DSB level (zero-nicking threshold). **e**, SSB-ddPCR results for multiple var-gRNAs in WT RPF1 cells, gRNAs with minimal off-target editing were selected for *SCN5A*^N406K^. The dotted line indicates the zero-nicking threshold from the standard curve. **f**, Representative images for detection of *SCN5A*^N406K^-edited cells. sgGOLDFISH signals indicate edited cells, and refGOLDFISH signals indicate *SCN5A* loci. Only colocalized signals are counted as true mutant calls (arrows). Scale bar = 5 μm. **g**, Optical recordings of action potentials from WT and *SCN5A*^N406K^-edited hiPSC-CMs infected with AOE-MyoV. **h**,**i**, Quantification of APD_50_ (**h**) and EADs (**i**) from the groups shown in (**g**). The dotted horizontal line indicates the median. WT, *n* = 282 cells; *SCN5A*^N406K^, *n* = 33 cells. Mean ± SEM. Statistical significance was determined using Welch’s t-test. \*\**P* < 0.01; \*\*\*\**P* < 0.0001. **j**, Representative images for detection of *RYR2*^E4950K^-edited cells. sgGOLDFISH signals indicate edited cells, and refGOLDFISH signals indicate *RYR2* loci. Only colocalized signals are counted as true mutant calls (arrows). Scale bar = 5 μm. **k**, Representative calcium transient traces from *RYR2*^E4950K^-edited hiPSC-CMs infected with AOE-MyoC. Red arrows indicate aCREs. **l**, Quantification of aCREs. The dotted horizontal line indicates the median. WT, *n* = 351 cells; *RYR2*^E4950K^, *n* = 77 cells. Mean ± SEM. Welch’s t-test: \*\*\*\**P* < 0.0001.

As varGOLDFISH has not previously been applied in hiPSC-CMs, we first evaluated its performance in this cell type using gRNAs targeting the reference sequence in unedited cells. Detection efficiencies ranged from approximately 20% to 60%, indicating reliable signal recovery in hiPSC-CMs (Fig. 5b and c). To rigorously evaluate specificity of gRNAs targeting variants (var-gRNAs), candidate var-gRNAs were screened using single-strand break digital droplet PCR (SSB-ddPCR)^25^ to quantify off-target activity in WT cells. Because uncleaved genomic DNA can confound ddPCR readouts, we estimated the zero-nicking threshold of this assay by using a restriction enzyme to generate a calibration curve (Fig. 5d). Var-gRNAs with apparent break frequencies at or below the calibrated zero-nicking threshold were classified as exhibiting no detectable off-target activity in WT cells and were therefore selected for downstream sgGOLDFISH analyses (Fig. 5e). For variants lacking suitable restriction enzyme sites for calibration, gRNAs with apparent break frequencies below 20% were selected based on empirical thresholds (Supplementary Fig. 4a and b).

Next, we extended varGOLDFISH to detect pathogenic variants introduced into hiPSC-CMs by prime editing. For these experiments, we focused on *SCN5A*^N406K^, linked to LQTS, and *RYR2*^E4950K^, associated with CPVT, covering both action potential and calcium-handling phenotypes. For each variant, we developed optimized var-gRNAs (Fig. 5d,e and Supplementary Fig. 4a). varGOLDFISH imaging results demonstrated robust detection of edited cells in both *SCN5A*^N406K^ (Fig. 5f, arrows) and *RYR2*^E4950K^ (Fig. 5j, arrows).

To link single-cell genotypes with their single-cell phenotypes, we further developed our CaT-FISH analysis pipeline to merge high-speed functional imaging results with high-resolution confocal varGOLDFISH imaging (Supplementary Fig. 5). After prime editing to introduce *SCN5A*^N406K^ or *RYR2*^E4950K^, cells underwent phenotyping using AOE-MyoV (*SCN5A*^N406K^) or AOE-MyoC (*RYR2*^E4950K^) platforms, followed by varGOLDFISH imaging to identify edited cells. Data analysis with the revised platform identified cells containing variants. Cells identified as carrying the *SCN5A*^N406K^ variant exhibited significant prolongation of APD (Fig. 5g,h and Supplementary Fig. 6a), along with an increased incidence of EADs, indicative of compromised repolarization stability (Fig. 5i). Notably, APD_50_ prolongation by *SCN5A*^N406K^ was greater when assessed by combined single-cell phenotyping and genotyping compared to analysis of the prime editor-treated pool (Fig. 4a), which contained both edited and unedited cells. For *RYR2*^E4950K^, combined single-cell phenotyping and genotyping demonstrated that the variant causes significantly increased aCRE frequency (Fig. 5k and l), whereas PEAR-based population analysis did not reach statistical significance for this phenotype (Fig. 4g).

Together, these results demonstrate the power of single-cell phenotyping coupled with single-cell genotyping to rapidly characterize the functional impact of variants in cardiac disease genes.

### Predicting the pathogenicity of patient-derived variants

To assess the utility of the platform for variant interpretation in a clinical context, we analyzed a VUS identified in a 5-year-old patient presenting with intermittent heart block and conduction abnormalities (Fig. 6a). The variant, TRPM4 p.A320V, lies within a highly conserved but intrinsically disordered region of the Ca²⁺-activated, non-selective cation channel TRPM4 (Transient Receptor Potential Melastatin 4)^49^ (Fig. 6b), and traditional variant prediction tools such as combined annotation dependent depletion (CADD)^50^ yielded inconclusive results.

**Fig. 6.**
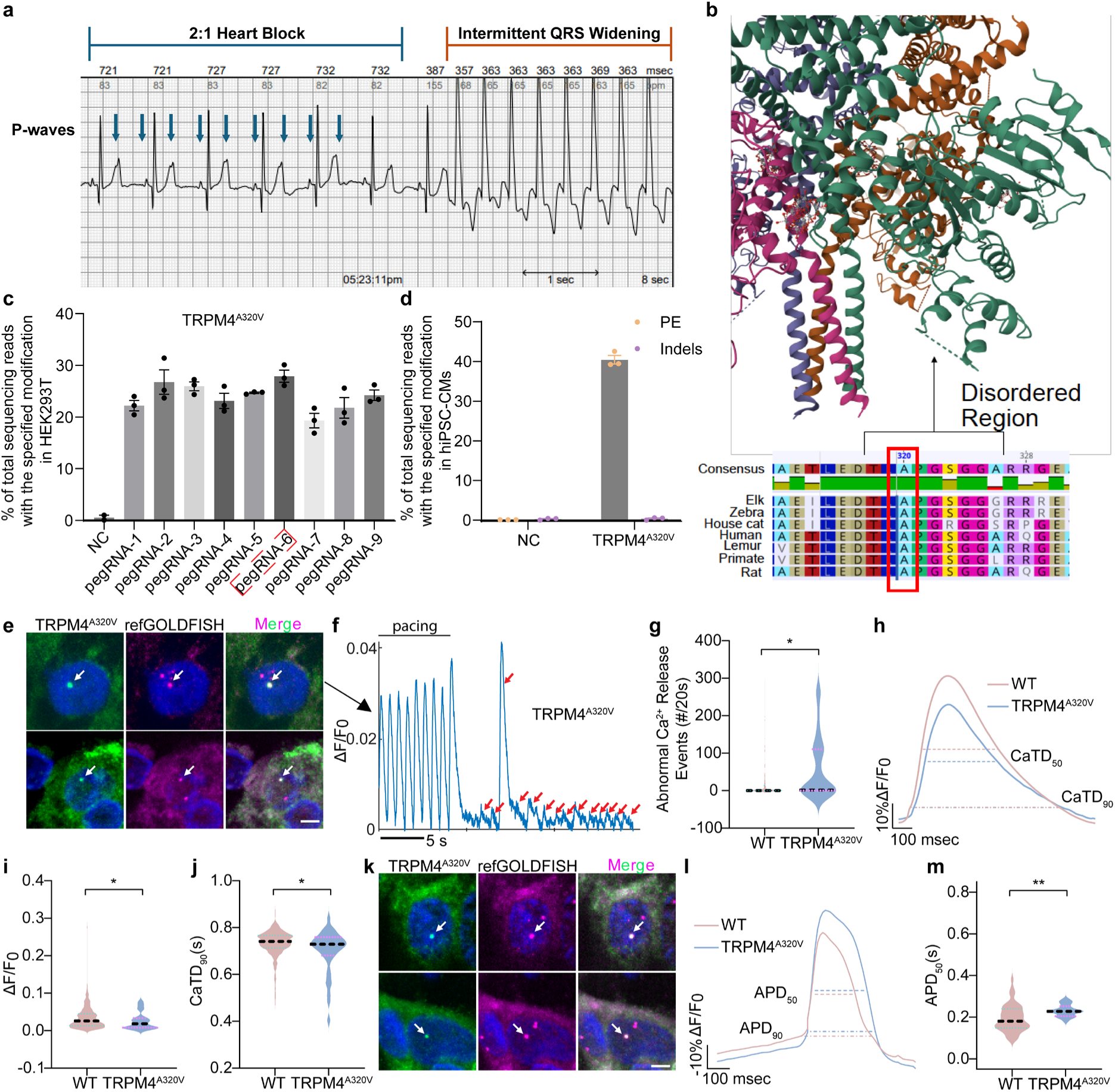
Functional reclassification of a patient-derived VUS. **a**, The ECG report from the patient carrying *TRPM4*^A320V^. **b**, The prediction of conservation for an inherently disordered region across species. **c**, Editing efficiencies of epegRNAs targeting *TRPM4*^A320V^ in HEK293T cells, as assessed by Sanger sequencing. Red rectangles highlight epegRNAs with the highest editing efficiency. Mean ± SEM from *n* = 3 independent biological replicates. **d**, Editing efficiencies of negative control (NC) and *TRPM4*^A320V^ in WT hiPSC-CMs, as assessed by amplicon sequencing. Mean ± SEM from *n* = 3 independent biological replicates. **e**, Representative images for detection of *TRPM4*^A320V^-edited cells infected with AOE-MyoC. sgGOLDFISH signals indicate edited cells, and refGOLDFISH signals indicate *TRPM4* loci. Only colocalized signals are counted as true mutant calls (arrows). Scale bar = 5 μm. **f**, Representative calcium transient traces from *TRPM4*^A320V^-edited hiPSC-CMs infected with AOE-MyoC. Red arrows indicate aCREs. **g**, Quantification of aCREs from WT and *TRPM4*^A320V^-edited hiPSC-CM. The dotted horizontal line indicates the median. WT, *n* = 335 cells; *TRPM4*^A320V^, *n* = 38 cells. Mean ± SEM. Welch’s t-test: \*\*\**P* < 0.001. **h**, Optical recordings of calcium transients from WT and *TRPM4*^A320V^-edited hiPSC-CMs infected with AOE-MyoC. **i**,**j**, Quantification of calcium transient amplitude (**i**) and CaTD_50_ (**j**) from the groups shown in (**h**). The dotted horizontal line indicates the median. WT, *n* = 335 cells; *TRPM4*^A320V^, *n* = 38 cells. Mean ± SEM. Statistical significance was determined using Welch’s t-test. \**P* < 0.05. **k**, Representative images for detection of *TRPM4*^A320V^-edited cells infected with AOE-MyoV. sgGOLDFISH signals indicate edited cells, and refGOLDFISH signals indicate *TRPM4* loci. Only colocalized signals are counted as true mutant calls (arrows). Scale bar = 5 μm. **l**, Optical recordings of action potentials from WT and *TRPM4*^A320V^-edited hiPSC-CMs infected with AOE-MyoV. **m**, Quantification of APD_50_ from the groups shown in (**l**). WT, *n* = 49 cells; *TRPM4*^A320V^, *n* = 18 cells. Mean ± SEM. Statistical significance was determined using Welch’s t-test. \*\**P* < 0.01.

Following epegRNA screening in HEK293T cells (Fig. 6c), the top-performing epegRNA was selected and achieved ∼40% prime-editing efficiency in WT hiPSC-CMs (Fig. 6d). To characterize the functional impact of this variant, we treated WT hiPSC-CMs with a prime editor to install *TRPM4*^A320V^. The population of edited and unedited cells was then functionally characterized using the AOE-MyoV and AOE-MyoC platforms. The same cells were then genotyped in situ by varGOLDFISH using an optimized var-gRNA (Supplementary Fig. 4b).

Calcium (Fig. 6e–j) and membrane voltage (Fig. 6k–m) parameters were compared between cells transfected with a control epegRNA and those transfected with the target epegRNA for *TRPM4*^A320V^. *TRPM4*^A320V^ cells exhibited pronounced abnormalities in calcium signaling, characterized by markedly more frequent aCREs (Fig. 6f and g). In addition, calcium transient amplitude was significantly reduced (Fig. 6h and i), while calcium transient duration was significantly shorter than controls (Fig. 6h and j). Optical action potential recordings revealed that *TRPM4*^A320V^ cells had significantly prolonged APD_50_, suggesting prolonged repolarization (Fig. 6l,m and Supplementary Fig. 6b).

Together, these results demonstrate that *TRPM4*^A320V^ induces mixed electrophysiologic dysregulation potentially consistent with its function as Ca²⁺-activated Na^+^ channel. This integrated genotype–phenotype analysis (Fig. 7) supports functional reclassification of *TRPM4*^A320V^ as a pathogenic variant and highlights the potential of the platform to inform clinical interpretation.

**Fig. 7.**
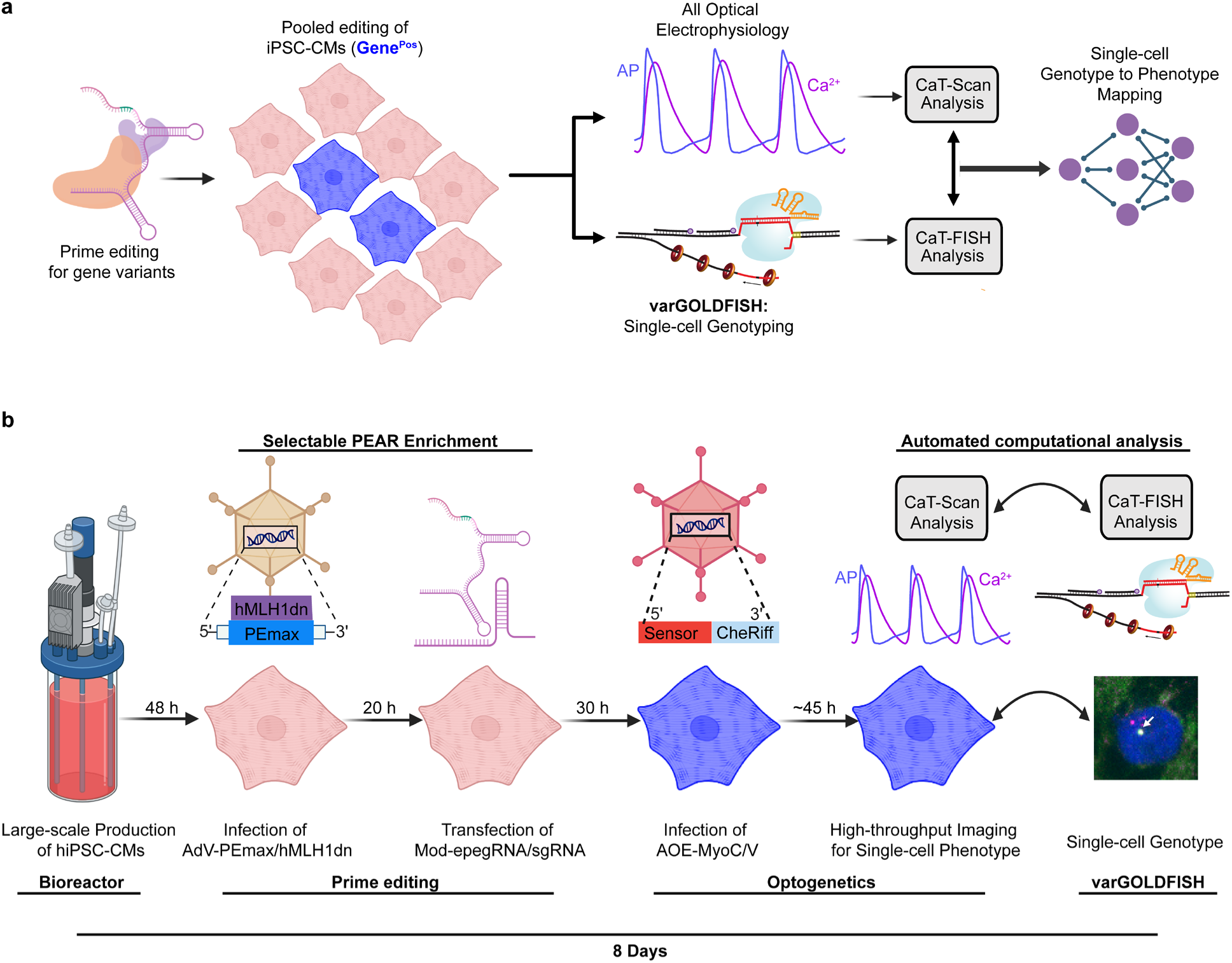
Integrated workflow for scalable genotype–phenotype mapping in hiPSC-CMs. **a**, The overview of single-cell genotype–phenotype mapping. **b**, The timeline details of single-cell genotype–phenotype mapping.

## Discussion

Existing strategies for assessing cardiac variants—such as clonal iPSC lines, surrogate cell systems, or bulk assays—are often low-throughput and insensitive to subtle, context-dependent effects^2,11,51,52^. To overcome these limitations, we developed REVi-SCOPE, an end-to-end integrated platform that enables direct functional interrogation of prime-edited hiPSC-CMs at single-cell resolution, without requiring clonal isolation or extended expansion. By combining single-cell optogenetic phenotyping with in situ single-cell genotyping, REVi-SCOPE links precise genomic edits to electrophysiological behavior within individual cells, providing quantitative and mechanistically interpretable readouts of variant function (Fig. 7). This strategy overcomes the low throughput of clone-based workflows and the reduced sensitivity of clone-free workflows that analyze populations containing a mixture of cells with and without the variant of interest.

We established several tiers of characterization with varying complexity and precision. For large effect variants with high editing efficiency (e.g., >50%), prime editing followed by population-level phenotyping may be sufficient to classify a variant as pathogenic. For intermediate cases, e.g. large effect variants with intermediate editing efficiency (e.g., 20–50%), PEAR enrichment may be sufficient for variant classification. However, both population level approaches may underestimate the variant’s effect size, due to inclusion of unedited cells. Combined single-cell phenotyping with single-cell genotyping yields the most rigorous assessment of variant effect, and may be needed for accurate classification of smaller effect variants or variants with low editing efficiencies (e.g., < 20%). By including a dual-probe approach with a strong signal-to-noise ratio, only cells containing colocalized sgGOLDFISH and refGOLDFISH signals were classified as edited. This enables accurate identification of edited hiPSC-CMs even at low editing efficiencies.

Understanding the potential pathogenicity of identified cardiac variants is a growing clinical problem, especially in cases of aborted cardiac arrest or victims of sudden death^53,54^.

Determining the pathogenicity of an identified variant in a victim of aborted cardiac arrest or sudden cardiac death (SCD) has critically important implications both for the patient or the surviving family members harboring the same genetic variant. If a variant is confirmed or suspected to be disease-causing, affected patients and at-risk family members can receive timely therapies and clinical follow-up, substantially reducing the risk of a fatal outcome. By significantly reducing the time and labor required for single-variant analysis, REVi-SCOPE enables pathogenicity determination on a clinically actionable timescale, directly improving outcomes for patients harboring potentially lethal genetic variants.

By enabling simultaneous characterization of Ca²⁺ handling and membrane voltage dynamics at single-cell resolution, REVi-SCOPE provides a powerful framework for uncovering unanticipated disease mechanisms and linking cellular phenotypes to clinical presentation. The demonstrated rapid analysis of the VUS *TRPM4*^A320V^ illustrates this capability. Functional interrogation of *TRPM4*^A320V^-edited cardiomyocytes revealed a complex electrophysiological phenotype characterized by increased arrhythmogenic calcium release events, reduced calcium transient amplitude and duration, and prolonged action potential repolarization. Aberrant TRPM4 activity has previously been associated with conduction slowing and atrioventricular block through membrane depolarization–mediated reduction of NaV1.5 availability^55,56^. The observed prolongation of repolarization and calcium-handling abnormalities in *TRPM4*^A320V^-edited cardiomyocytes are consistent with dysregulated depolarizing current and excitation–contraction coupling, providing a mechanistic framework linking the cellular phenotype to the patient’s intermittent heart block and conduction abnormalities. These findings support a pathogenic role for *TRPM4*^A320V^ and illustrate how integrated single-cell genotype–phenotype analysis can bridge molecular dysfunction to clinical electrophysiology.

While this study establishes REVi-SCOPE as the first end-to-end method for the rapid characterization of potentially lethal cardiac gene variants, several important considerations remain beyond the focus of this current work. Although we were able to determine the effects of gene variants, either known to be disease-causing or possibly disease-causing, the exact sensitivity and specificity of the platform is yet to be determined. Additionally, testing of both pathogenic and benign variants with comparison to clinical outcomes would firmly establish REVi-SCOPE as a robust clinical diagnostic tool. Increasing the number of tested variants may also reveal emergent single-cell phenotypes beyond those tested in this present work that would be an excellent application of machine learning techniques for extended phenotypic extraction. Another limitation of this study is the focus on Ca^2+^ and membrane dynamics, which restricts the scope of testable cardiac variants to those related to arrhythmia. Further iterations of the REVi-SCOPE platform could include any imaging-based phenotype indicated by biosensors or functional classification at the single-cell level. REVi-SCOPE’s functionality could also be extended to other cell types, such as iPSC-derived neurons, greatly expanding the range of clinically relevant gene variants that can be characterized. While REVi-SCOPE’s current throughput is lower than that of large-scale sequencing-based methods^57^, its unique ability to directly link variant genotype to changes in live-cell physiology positions it as the ideal tool for rapidly characterizing variants predicted to dynamically affect cardiac function. In the future, REVi-SCOPE will likely be used alongside complementary variant analysis methods to comprehensively evaluate genetic variants, improving clinical outcome prediction and advancing precision medicine.

Looking ahead, the single-cell sensitivity and precise genotypic resolution of our platform position it for high-throughput and pooled prime-editing applications. While epegRNA optimization was performed in HEK293T cells in this study, direct genotyping of edited cardiomyocytes is expected to minimize the need for extensive epegRNA pre-screening, which would substantially reduce experimental effort while broadening the scope of variant discovery and prioritization. Methods for the simultaneous introduction and identification of multiple variants could also be employed through the sequential application^58^ of multiple varGOLDFISH probes. This platform could also be leveraged for other cardiac biosensor readouts, including contractility, cell survival, or drug responses, or other cell types such as neurons. Together, this platform provides a generalizable framework for linking genotype to phenotype in cardiac diseases and offers a scalable route toward functional interpretation of human genetic variation and development of precision therapies.

## Methods

### Culture and infection of hiPSC-CMs

Stocks of hiPSC-CMs were generated using a stirred bioreactor differentiation protocol^43^. Cryopreserved hiPSC-CMs were thawed at 37°C, gently diluted dropwise with RPMI 1640 medium (Thermo Fisher Scientific, 11875093), and pelleted at 200 × g for 5 min. Cells were resuspended in RPMI 1640 containing B-27™ Supplement (Thermo Fisher Scientific, 17504044) and 10 μM Y-27632 (R&D Systems, 1254), and plated onto 1% Geltrex-coated (Thermo Fisher Scientific, A1413302) culture vessels. After 24 h, the medium was replaced with RPMI/B-27 without Y-27632, and cells were maintained for ≥ 24 h before adenoviral transduction at the indicated MOI. After infection for 16 h, the medium was replaced with fresh RPMI/B-27.

Both CPVT line (*RYR2*^G3946S/+^)^32^ and LQTS line (*CALM1*^F142L/+^)^36^ were established in the WTC-11 (male origin; Coriell) background via CRISPR-Cas9 genome editing. All cell cultures were routinely tested for mycoplasma contamination (monthly, negative throughout).

### Optical recording of action potentials and calcium transients

Membrane voltage and Ca²⁺ transients were recorded using a Kinetic Image Cytometer (KIC; Vala Biosciences). Prior to imaging, hiPSC-CMs were stained with 40 µM Hoechst 33342 (Santa Cruz Biotechnology, sc-394039) and 2 µM Calcein AM (Thermo Fisher Scientific, C34852) for 15 min at 37°C. Following incubation, cells were washed once with fresh medium, and the culture plate was transferred to the pre-warmed KIC system. Recordings were acquired using a Nikon 20× Plan Apo VC objective at a frame rate of 90 Hz for 30 s. Optical pacing was applied at 1 Hz with 10 pulses (8 ms pulse width) using 15% intensity in the cyan stimulation channel. Emission signals were collected on the TRITC detection channel. Data was analyzed using a custom-written CaT-Scan.

### Analysis pipeline of single-cell phenotype-genotype mapping

Cytoplasmic (Cyto-Mask) and nuclear (Nuc-Mask) masks of individual hiPSC-CMs from KIC imaging were generated separately using Cellpose3 segmentation software^42^. Calcium and membrane-voltage parameters were extracted from KIC recordings using CaT-Scan. Following KIC imaging, cells were labeled with sgGOLDFISH and refGOLDFISH probes and imaged as 3 × 3 tiled fields with z-stack acquisition under 405-, 561-, and 640-nm laser excitation to detect nuclear, sgGOLDFISH, and refGOLDFISH signals, respectively.

In the CaT-FISH pipeline (Supplementary Fig. 5), we first generated a coarse nuclear mask from the confocal nuclear channel and used it to register the confocal and KIC images. Registration was performed by identifying the pixel shift that maximized the 2D cross-correlation between the nuclear masks. After alignment, the confocal image was cropped to the matched field of view for downstream analysis.

Nuclei within the cropped image were segmented from the nuclear channel and assigned unique cell identifiers. refGOLDFISH puncta were detected using a difference-of-Gaussians filter followed by an empirically optimized intensity threshold. Cells positive for refGOLDFISH were then evaluated for sgGOLDFISH. sgGOLDFISH puncta exceeding an optimized threshold were classified as genotype-positive (edited) events.

Nuclear masks from genotype-positive (edited) cells identified were spatially aligned with Nuc-Masks derived from KIC imaging. Genotype-positive Nuc-Masks were retained only when one-to-one alignment was observed between confocal and KIC images. For each retained Nuc-Mask, the corresponding cytoplasmic mask was used to extract phenotypic calcium and voltage parameters. Only data from genotype-positive hiPSC-CMs were included in downstream statistical analyses.

### Prime editing in hiPSC-CMs

Cells were seeded at a density of 7×10^4^ cells per well on 96-well plates (Greiner Bio-One, 655892) pre-coated with 1% Geltrex. At 48 h after plating, cells were co-infected with AdV-PEmax (MOI = 10) and AdV-hMLH1dn (MOI = 3). 16 h following infection, the culture medium was exchanged for RPMI 1640 supplemented with B-27™ Supplement. 20 h post-infection, modified epegRNAs (5 pmol) and sgRNAs (3.25 pmol) were introduced using Lipofectamine™ RNAiMAX Transfection Reagent (Thermo Scientific, 13778100) according to the supplier’s guidelines. Synthetic epegRNAs and sgRNAs (Supplementary Table 1) were obtained from Integrated DNA Technologies and incorporated 2′-O-methyl modifications at the first and last three nucleotides together with phosphorothioate linkages at the terminal three nucleotides; oligonucleotides were used without further processing. Cells were harvested 72 h after RNA delivery for downstream evaluation of genome editing.

### varGOLDFISH in hiPSC-CMs

Recombinant eCas9 and Rep-X were produced and purified, and sgGOLDFISH/refGOLDFISH guide RNAs and probe sequences were designed following established strategies^25,26^. Custom oligonucleotide FISH probes (Supplementary Table 2) were obtained from Integrated DNA Technologies and fluorescently conjugated with Cy3B-NHS (Lumiprobe, 29320) or Cy5-NHS (Lumiprobe, 23320) esters using standard amine-reactive labeling chemistry. gRNAs (Supplementary Table 3) were generated by in vitro transcription with the EnGen® sgRNA Synthesis Kit (S. pyogenes; NEB, E3322V) according to the supplier’s recommendations.

For cellular labeling, hiPSC-CMs were fixed in pre-chilled methanol: acetic acid (1:1, v/v) at - 20°C for 20 min, followed by three washes in PBS. Fixed cells were equilibrated in binding-blocking buffer (20 mM HEPES, pH 7.5; 100 mM KCl; 7 mM MgCl₂; 5% glycerol; 0.1% Tween-20; 1% BSA; supplemented immediately before use with 1 mM DTT and 0.1 mg mL⁻¹ *E. coli* tRNA) for 10 min at 37°C. Ribonucleoprotein complexes were assembled by combining eCas9 nickase (100 nM) with var-gRNA (100 nM) in a small volume of (1/10 of the final volume) binding-blocking buffer and incubating for 15 min at room temperature. For refGOLDFISH experiments, additional eCas9 nickase (20 nM) and multiple guide RNAs targeting the downstream region of the mutation site (20 nM each) were included during complex formation.

Cells were subsequently incubated with assembled RNPs for 2 hours at 37°C. After incubation, RNP buffer was taken off and the cells were supplemented with ATP (2 mM) and Rep-X (400 µM) for 90 min at 37°C in binding-blocking buffer, followed by three PBS washes. Residual RNA was removed by treatment with RNase Cocktail™ Enzyme Mix (Invitrogen, AM2286) diluted 1:100 in PBS for 30 min at 37°C, after which cells were washed three additional times with PBS. Samples were pre-equilibrated in freshly prepared hybridization buffer (20% formamide, 2× SSC, 0.1 mg mL⁻¹ E. coli tRNA, 10% dextran sulfate, 2 mg mL⁻¹ BSA) for 10 min at room temperature. Fluorescent oligonucleotide probes (0.2 nM per oligo) were then applied in the hybridization buffer and incubated for 1.5 h at 37°C.

Post-hybridization washes were performed three times using a wash buffer (30% formamide, 2× SSC), including a 15-min wash at 37°C followed by a 5-min wash at room temperature with gentle agitation (40 rpm). Cells were rinsed three times with PBS, counterstained with Hoechst 33342 Ready Flow™ Reagent (Invitrogen, R37165) diluted 1:1000 for 2 min at room temperature, and transferred into imaging buffer (1 mM Trolox, 1 mM Trolox Quinone, and 2 mM PCA) supplemented with PCO (3U/mL OYC Americas #46852004, 50 mM Tris-HCL, pH 8.0, 50 mM KCl) immediately before imaging.

### Imaging for prime edited cells labeled by varGOLDFISH

To enable registration of identical fields of view across imaging modalities, membrane voltage and Ca²⁺ transients were first recorded using a KIC equipped with a Nikon 20× Plan Apo VC objective. High-resolution imaging was subsequently performed on a Nikon Ti2 inverted microscope coupled to a spinning-disk confocal module (Nikon Instruments) using a Plan Fluor 40×/1.30 NA oil-immersion objective. Images were acquired as 3 × 3 tiled fields with z-stack collection (14-µm total depth, 21 optical sections) under 405-, 561-, and 640-nm laser excitation. Large-field image acquisition was carried out using Nikon Digital Sight 50M and Digital Sight 10 CMOS cameras (Nikon Instruments).

### Development of an end-to-end pipeline for high-speed calcium imaging data analysis

CaT-Scan is an end-to-end pipeline that takes high-speed functional imaging videos of fluorescent biosensors as inputs, and outputs single-cell quantifications of calcium transient or membrane voltage parameters and aCREs or EADs. To do so, the following steps are performed: 1) down-sampling of imaging videos; 2) automating cardiomyocyte segmentation; 3) calcium transient or action potential quality control and pre-processing; 4) quantification of calcium transient or actional potential parameters and aCREs/EADs.

Unprocessed high-speed videos are 2048 x 2048 pixels for 30-35 seconds, amounting to >10 Gb per video. In our hands, we empirically chose to down-sample the videos to a size of 256 x 256 pixels (amounting to ∼1Gb per video) to balance data integrity, processing time, and data storage.

Next, active cardiomyocytes were segmented. To do so, a pixel-wise standard deviation operation was performed during the pacing period of a down-sampled video. Segmentation was then performed on this image using a Cellpose backbone^42^ that was fine-tuned on 30 manually segmented hiPSC-CM videos. Fine-tuning was performed using a 5:1 train: validation split, and a learning rate of 1 × 10^-4^ for 10 epochs.

Next, the mean fluorescent intensity of each biosensor detected within each segmented cardiomyocyte was retrieved as a function of time. For each cell, a custom MATLAB script was implemented (Fig. 2b). Pre-processing included denoising (via a median filter, wavelet signal denoising, and Savitzky-Golay filtering), removing photobleaching effect (via a moving minimum function), and data normalization (relative to the moving minimum). Cells without beat frequencies that match the predetermined pacing frequency were removed. Standard calcium transient or action potential parameters were then obtained per beat and averaged across all pacing beats. Finally, aCREs or EADs were identified with a minimum threshold of 5% of the mean pacing amplitude, and the metrics included number and duration.

### Construction of Adenovirus

Target genes (Supplementary Table 4) were cloned into the pENTR/TEV/D-TOPO vector (Invitrogen, K2525-20). Recombinant adenoviruses were generated using the Invitrogen adenoviral expression system (Invitrogen, V493-20) and propagated in HEK293A cells maintained in Dulbecco’s modified Eagle medium (Thermo Scientific, 11965118) supplemented with 10% fetal bovine serum (Thermo Scientific, A5670701). Viruses were purified with Vivapure AdenoPack columns (Sartorius, VS-AVPQ020), and titers were determined using the Adeno-X qPCR Titration Kit (Takara, 632252).

### Benchmarking to existing calcium transient software

We benchmarked CaT-Scan to previously published software—CalTrack^41^. To do so, randomly selected hiPSC-CM studies were manually segmented by a blinded and experienced hiPSC-CM researcher to serve as the human gold standard. None of these studies overlapped with the fine-tuning samples. The segmentation performance of CaT-Scan and CalTrack was evaluated relative to the human annotations using standard metrics, including the Jaccard index and F1 score, with higher values indicating better performance.

### Molecular cloning of epegRNAs and sgRNAs

Prime-editing guide constructs were assembled using a type IIS–based Golden Gate strategy^21^. Plasmid backbones supporting either epegRNA expression (Addgene plasmid 174038) or the associated nicking sgRNA (Addgene plasmid 65777) were first linearized by overnight digestion with the appropriate type IIS enzyme (BsaI-HFv2 or BsmBI-v2; New England BioLabs). Linear products were separated on 1% agarose gels, and the desired bands were purified using a column-based gel extraction kit (Takara, 740609).

Guide-specific DNA oligonucleotides incorporating the spacer, scaffold, and any 3′ extension sequences were synthesized by Integrated DNA Technologies. Complementary strands were combined at equal molar ratios, heated to 95°C for 3 min, and allowed to cool slowly to ambient temperature to facilitate duplex formation. Annealed oligos were phosphorylated with T4 polynucleotide kinase at 37°C for 1 h.

Assembly reactions (10 µL) contained the purified linear vector, phosphorylated duplex oligos, the corresponding type IIS restriction enzyme, and T4 DNA ligase. Reactions were incubated at room temperature for an initial ligation phase and then subjected to cycling between 16°C and 37°C (5 min at each temperature, eight cycles). A final 37°C ligation step and heat inactivation at 80°C were performed before holding reactions at 12°C. Reaction mixtures were transformed into chemically competent *E. coli*. Individual colonies were screened by Sanger sequencing to confirm correct guide assembly.

### Flow cytometry

Single-cell suspensions of hiPSC-CMs were generated by enzymatic dissociation using ACCUMAX™ (STEMCELL Technologies Inc, 07921). To determine differentiation efficiency, cells were chemically fixed in 4% paraformaldehyde for 15 min at 4°C and subsequently permeabilized for 1 h at 4°C in a buffered solution composed of PBS, 5% (v/v) FCS/FBS (R&D Systems, S11150), 0.5% (w/v) saponin (Sigma, 47036-50G-F), and 0.05% (w/v) sodium azide (Sigma, S2002). Cells were collected by centrifugation (200 × g, 5 min), resuspended in permeabilization buffer, and labeled with FITC-conjugated anti-cardiac troponin T antibody (1:100; Miltenyi Biotec, REA400) or the corresponding human IgG1 isotype control (1:100; Miltenyi Biotec, REA293) for 1 h at 4°C. After washing, samples were transferred to PBS and measured on an LSRFortessa flow cytometer (BD Biosciences).

For isolation of prime editing activity reporter (PEAR)-positive hiPSC-CMs, dissociated cells were resuspended in PBS and sorted using a SH800S benchtop cell sorter (Sony, Tokyo, Japan). Flow cytometry data were analyzed using FlowJo software (BD Biosciences).

### Immunofluorescence

hiPSC-CMs were chemically fixed in 4% paraformaldehyde for 15 min at room temperature, followed by three washes in PBS. Cell membranes were permeabilized using 0.3% Triton X-100 in PBS for 15 min and washed again three times with PBS. Non-specific binding was minimized by incubation in a blocking solution consisting of 5% bovine serum albumin and 10% donkey serum in PBS for 1 h at room temperature. Samples were then incubated with an anti-cardiac troponin T primary antibody (1:250; Abcam, ab45932), anti-Cas9 (1:200; Cell Signaling Technology, 14697) and anti-HA (1:200; Cell Signaling Technology, 3724) prepared in the blocking solution at 4°C overnight. After extensive PBS washing, cells were incubated with Alexa Fluor™ Plus 488/555-conjugated secondary antibody (Thermo Fisher Scientific, A32790TR/A32794/A32790TR) for 1 h at room temperature. Following final washes in PBS, samples were prepared for imaging.

### Prime editing in HEK293T cells

HEK293T cells were plated in 24-well tissue-culture plates at a density of 5 × 10⁴ cells per well in DMEM supplemented with 10% fetal bovine serum. Cells were allowed to adhere overnight, and transfections were performed the following day when cultures reached approximately 50–60% confluence. Plasmid delivery was carried out using Polyethylenimine “Max” (PEI MAX) (Histo-Line Laboratories, 24765) following the manufacturer’s instructions. For each well, a mixture containing 1 µg of the prime editor expression plasmid, 200 ng of the epegRNA construct, and 65 ng of the nicking sgRNA plasmid was used. Following a 72-h incubation period, cells were harvested for genomic DNA isolation. Editing efficiencies were quantified from Sanger sequencing traces using the EditR software package. All epegRNAs and nicking sgRNAs sequences used in HEK293T cells are listed in Supplementary Table 5.

### Genomic DNA extraction

Cells were rinsed once with PBS (Gibco, 10010023) before genomic DNA was released using the KAPA Mouse Genotyping Kit (KK7302). Lysates were incubated at 75°C for 10 min to facilitate digestion, followed by a 5 min inactivation step at 95°C.

### Preparation of amplicon sequencing samples

Primers (Supplementary Table 6) targeting the regions of interest were synthesized by Integrated DNA Technologies. Optimal annealing temperatures were identified by gradient PCR and verified by agarose gel electrophoresis to ensure product specificity. Genomic loci (Supplementary Table 7) were amplified using PrimeSTAR HS DNA Polymerase (Takara, R010B), and resulting amplicons were purified with the QIAquick Gel Extraction Kit (Qiagen, 28706) and eluted in 40 µL of nuclease-free water. DNA concentrations were quantified with the Qubit dsDNA Assay Kit (Thermo Scientific, Q32851). For deep sequencing, 500 ng of purified PCR products were submitted to GENEWIZ (Azenta Life Sciences) for library preparation, amplicon sequencing, and data processing. The data analysis was performed using CRISPResso2 (https://crispresso2.pinellolab.org/submission).

### SSB-ddPCR

SSB-ddPCR was conducted using a previously established workflow^25^, with no changes to the overall procedure. In brief, fixed RPF1 cells were subjected to Cas9 nickase-mediated strand break generation (gRNAs used are provided in Supplementary Table 8), followed by genomic DNA isolation and droplet digital PCR analysis as described. For this study, locus-specific primers and hydrolysis probes (Supplementary Table 9) were newly designed for each target locus and synthesized by Integrated DNA Technologies. All reaction components, enzyme concentrations, droplet generation steps, PCR cycling conditions, and data acquisition settings were identical to the published protocol, except that an annealing temperature of 58 °C was used, and restriction enzymes included in the ddPCR reaction were selected based on the target locus (HindIII-HF for *SCN5A*^N406K^ and *TRPM4*^A320V^; BamHI-HF for *RYR2*^E4950K^). For generation of the standard curve for *SCN5A*^N406K^, double-strand breaks were introduced using SexAI (New England Biolabs, R0605S) as specified by the manufacturer, and genomic DNA containing double- and single-strand breaks was mixed at defined ratios (100%, 90%, 70%, 50%, 30%, 20%, 10%, and 0%).

### Statistics

Statistical analyses were performed using Prism 10 (GraphPad, San Diego, CA, USA). Data are presented as mean ± SEM. Welch’s t-test was used for comparisons between two groups with normally distributed data, whereas the Kolmogorov–Smirnov test was used for non-normally distributed samples. For comparisons among three groups with non-normally distributed data, the Kruskal–Wallis test (nonparametric one-way ANOVA) was used. The statistical test applied to each dataset is indicated in the corresponding figure legend. Differences were considered statistically significant at P < 0.05.

## Data availability

All data supporting the findings of this study are available from the corresponding authors upon reasonable request.

## Code availability

The source code for CaT-Scan and CaT-FISH is available on GitHub: https://github.com/lonehusky76/CaTScan

## Acknowledgements

X.-T.W. was supported by an American Heart Association Postdoctoral Fellowship (AHA25POST1379370). V.J.B. was supported by the American Heart Association (AHA953610). T.H. was supported by National Institutes of Health grants R35-GM122569 and U01 DK127432, and by the NSF Science and Technology Center Quantitative Cell Biology (2243257). K.S. and V.J.B. were supported by NCATS Tissue Chips Consortium (UH3 TR003279). Y.J. was supported by R01HL167450. We acknowledge the Microscopy Resources on the North Quad Core at Harvard Medical School, in particular Paula Montero Llopis and Praju Vikas Anekal, for technical support and access to instrumentation. We thank Alexander Sousa (David R. Liu laboratory) for assistance with prime editing, Yanbo Wang (Taekjip Ha laboratory) for assistance with sgGOLDFISH, and Nicholas Bezzerides for assistance with the code.

## Author contributions

X.-T.W. and V.J.B. conceived and designed the study. T.H. directed the varGOLDFISH experiments. X.-T.W. performed experiments, analyzed the data, and drafted the manuscript. P.-T.C., X.-T.W. and T.S. conducted varGOLDFISH experiments. J.M., L.R., P.-T.C., N.P., K.S., Y.J., W.P. and K.K.P. contributed to analysis code development. Y.T. generated bioreactor-derived hiPSC-CMs. C.M.J. and D.M. provided clinical information related to the patient variant. W.P., V.J.B. and T.H. revised the manuscript. X.-T.W. and V.J.B. finalized the manuscript with input from all authors. All authors read and approved of the final manuscript.

