## Supplemental data for "Rapid functional classification of cardiac genetic variants directly informs precision cardiology"

### Supplementary information

1. Supplementary Figures 1 to 6
2. Supplementary Tables 1 to 9

#### Supplementary Figures

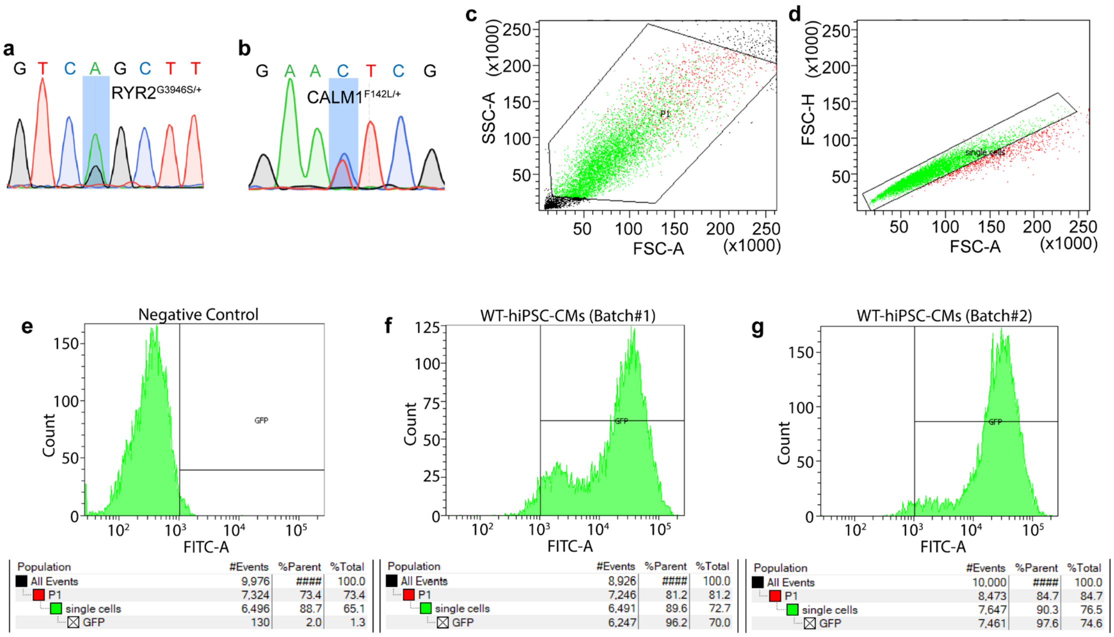


**Supplementary Figure. 1 | Genotyping of disease-specific iPSC lines and quality assessment of WT hiPSC-CMs. a**, Sanger sequencing confirming the *RYR2*$^{G3946S/+}$ variant in CPVT iPSCs. **b**, Sanger sequencing confirming the *CALM1*$^{F142L/+}$ variant in LQTS iPSCs. **c**–**e**, Flow cytometry gating strategy established using WT hiPSC-CMs stained with an IgG1 isotype control antibody. **f**,**g**, Percentage of cTNT-positive cells in two independent batches of WT hiPSC-CMs, quantified by flow cytometry using the gates defined in (**c**–**e**).

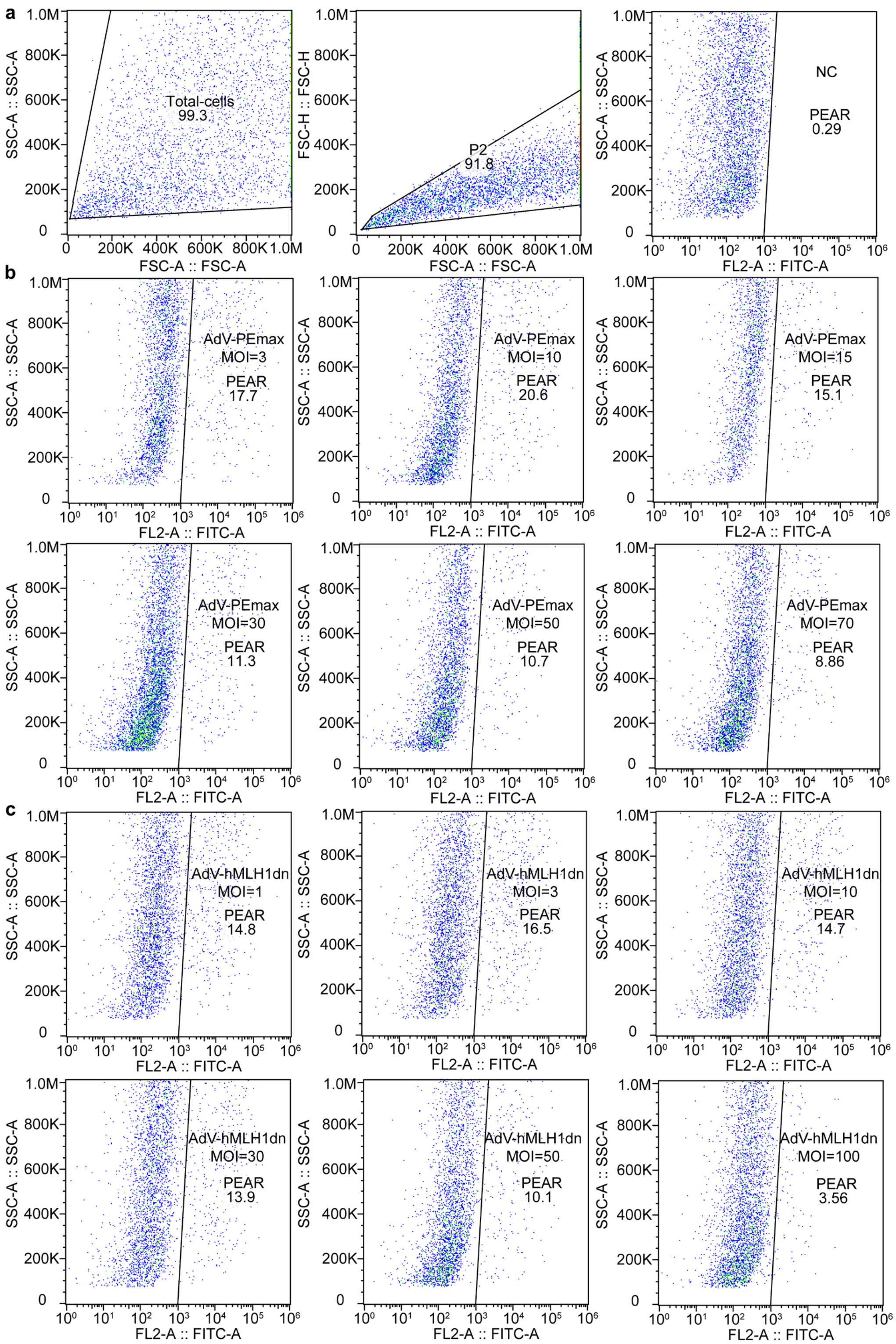


**Supplementary Figure. 2 | Dose optimization of AdV-PEmax and AdV-hMLH1dn. a**, FACS gating strategy established using WT hiPSC-CMs infected with AdV-PEAR. **b**, Percentage of $GFP^+$ cells in WT hiPSC-CMs infected with AdV-PEAR and gradient doses of AdV-PEmax, quantified by FACS using the gates defined in (**a**) and analyzed with FlowJo. **c**, Percentage of GFP$^{+}$ cells in WT hiPSC-CMs infected with AdV-PEAR and AdV-PEmax together with gradient doses of AdV-hMLH1dn, quantified by FACS using the gates defined in (**a**) and analyzed with FlowJo.

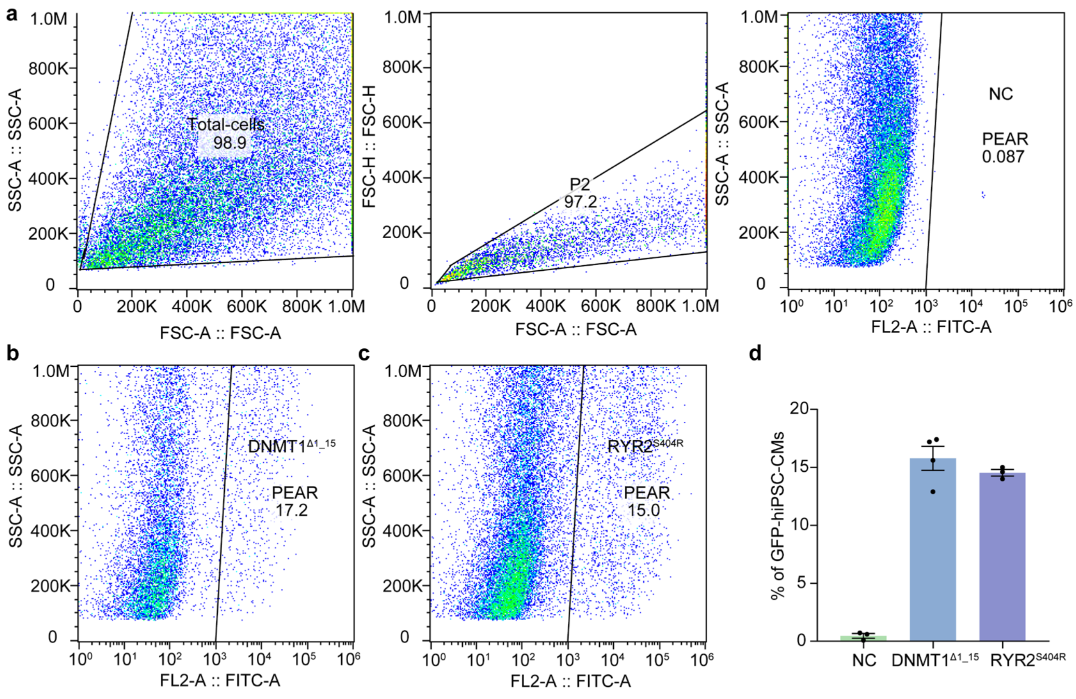


**Supplementary Figure. 3 | FACS of PEAR-enriched hiPSC-CMs. a**, Gating strategy in WT hiPSC-CMs infected with AdV-PEAR and edited with scrambled epegRNA. **b**,**c**, Fraction of GFP$^{+}$ cells after prime editing targeting *DNMT1*$^{\Delta 1\_15}$ (**b**) and *RYR2*$^{S404R}$ (**c**), quantified by FACS using gates from (**a**) and analyzed with FlowJo. **d**, Summary of sorting percentages across conditions analyzed with FlowJo. Mean ± SEM.

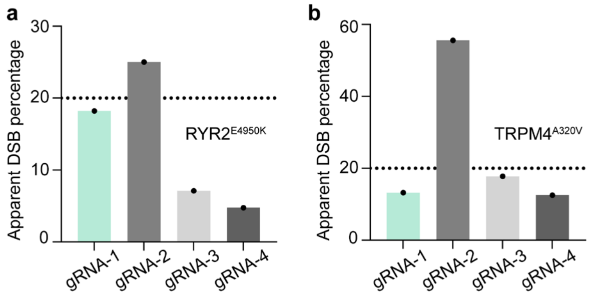


**Supplementary Figure. 4 | SSB-ddPCR screening of var-gRNAs. a**,**b**, SSB-ddPCR results for multiple var-gRNAs in WT RPF1 cells. var-gRNAs with minimal off-target nicking on WT cells were selected for the sgGOLDFISH of *RYR2*$^{E4950K}$ (**a**) and *TRPM4*$^{A320V}$(**b**). The dotted line indicates the zero-nicking threshold.

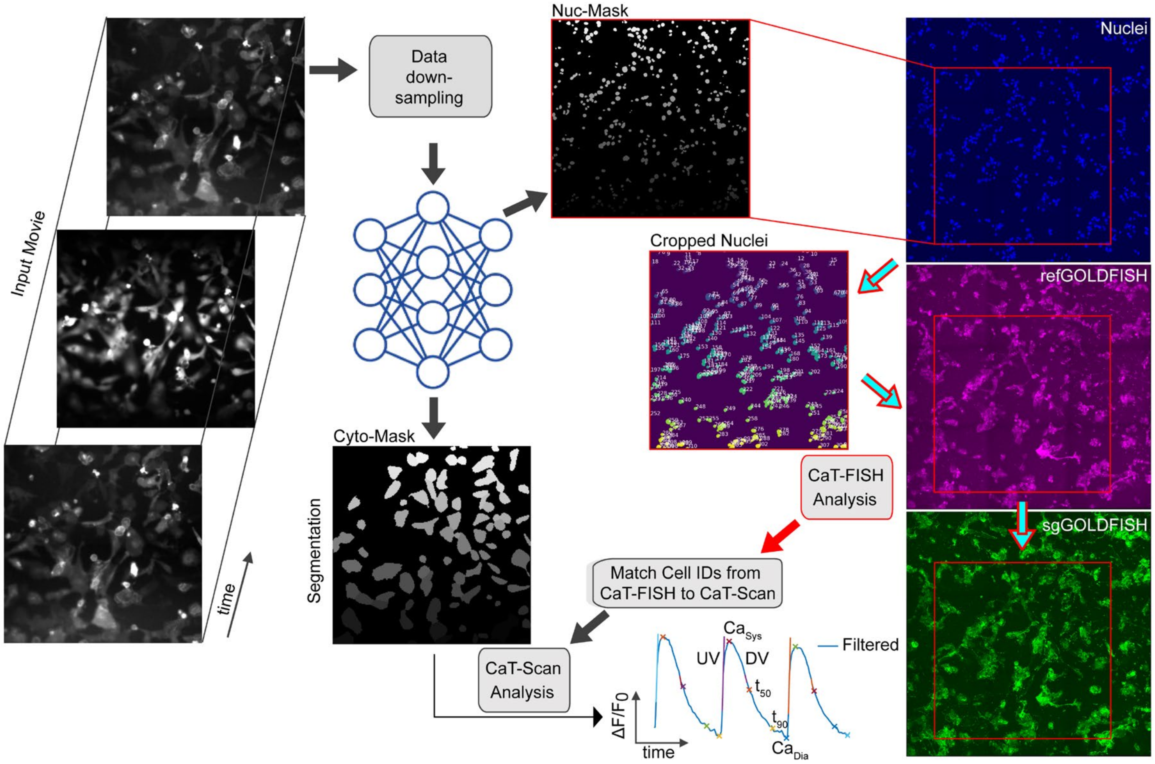


**Supplementary Figure. 5 | Analysis pipeline of single-cell genotype–phenotype mapping.**

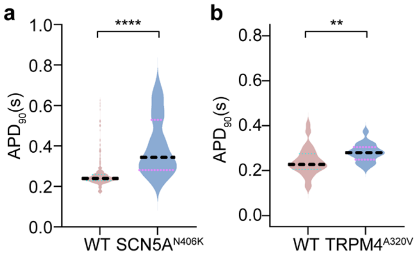


**Supplementary Figure. 6 | APD prolongation in the LQTS variant and VUS.** Optical recordings of action potentials from hiPSC-CMs infected with AOE-MyoV. The dotted horizontal line indicates the median. **a**, $APD_{90}$ in WT and *SCN5A*$^{N406K}$-edited hiPSC-CMs (WT, $n = 282$ cells; *SCN5A*$^{N406K}$, $n = 33$ cells). **b**, $APD_{90}$ in WT and *TRPM4*$^{A320V}$-edited hiPSC-CMs (WT, $n = 49$ cells; *TRPM4*$^{A320V}$, $n = 18$ cells). Mean ± SEM; Welch's t-test; $^{**}P < 0.01$, $^{****}P < 0.0001$.

#### Supplementary Tables

**Supplementary Table 1.** Sequences of epegRNAs and sgRNAs used in hiPSC-CMs.

| | |
|---|---|
| **Fig. 3l** | |
| RYR2$^{S404R}$-epegRNA | GCAUGAAGGCCACAUGGAUGAGUUUUAGAGCUAGAAA UAGCAAGUUAAAAUAAGGCUAGUCCGUUAUCAACUUGA AAAAGUGGCACCGAGUCGGUGCGACAAACGUAUGCCAU CAUCCAUGUGGCCUUCGCGGUUCUAUCUAGUUACGCGU UAAACCAACUAGAAUUU |
| RYR2$^{S404R}$-sgRNA | GAACGUAUGCCAUCAUCCAUGGUUUUAGAGCUAGAAAU AGCAAGUUAAAAUAAGGCUAGUCCGUUAUCAACUUGAA AAAGUGGCACCGAGUCGGUGCUUU |
| RYR2$^{E4950K}$-epegRNA | GAUGUAUCAAGAAAGGUGUUGUUUUAGAGCUAGAAAU AGCAAGUUAAAAUAAGGCUAGUCCGUUAUCAACUUGAA AAAGUGGCACCGAGUCGGUGCGCUGGGAAAAUUUCCA ACACCUUUCUUGAUACGCGGUUCUAUCUAGUUACGCGU UAAACCAACUAGAAUUU |
| RYR2$^{E4950K}$-sgRNA | GCUGGUCUUCAUACUGUUUCGUUUUAGAGCUAGAAAUA GCAAGUUAAAAUAAGGCUAGUCCGUUAUCAACUUGAAA AAGUGGCACCGAGUCGGUGCUUU |
| **Fig. 3o** | |
| SCN5A$^{N406K}$-epegRNA | GUUGCGACCACGGCCAGGAUCGUUUUAGAGCUAGAAAU AGCAAGUUAAAAUAAGGCUAGUCCGUUAUCAACUUGAA AAAGUGGCACCGAGUCGGUGCUCUACCUGGUGAAACUG AUCCUGGCCGUGCGCGGUUCUAUCUAGUUACGCGUUAA ACCAACUAGAAUUU |
| | GCUACCUGGUGAAACUGAUCCGUUUUAGAGCUAGAAAU |

| | |
|---|---|
| SCN5A$^{N406K}$-sgRNA | AGCAAGUUAAAAUAAGGCUAGUCCGUUAUCAACUUGAAAAAGUGGCACCGAGUCGGUGCUUU |
| KCNQ1$^{A341V}$-epegRNA | GCCCACGGGGCACCUACCGCUGUUUUAGAGCUAGAAAUAGCAAGUUAAAAUAAGGCUAGUCCGUUAUCAACUUGAAAAAGUGGCACCGAGUCGGUGCUUCUUUGUGCUGCCAGCGGUAGGUGCCCCGUGCGCGGUUCUAUCUAGUUACGCGUUAAACCAACUAGAAUUU |
| KCNQ1$^{A341V}$-sgRNA | GCUCCUUCUUUGUGCUGCCAGGUUUUAGAGCUAGAAAUAGCAAGUUAAAAUAAGGCUAGUCCGUUAUCAACUUGAAAAAGUGGCACCGAGUCGGUGCUUU |
| **Fig. 4d** | |
| DNMT1$^{Δ1_15}$-epegRNA | GAUUCCUGGUGCCAGAAACAGUUUUAGAGCUAGAAAUAGCAAGUUAAAAUAAGGCUAGUCCGUUAUCAACUUGAAAAAGUGGCACCGAGUCGGUGCUGCUAAGGACUAGUUCUGCCCUUCUGGCACCAGGACCUCUUCUCGCGGUUCUAUCUAGUUACGCGUUAAACCAACUAGAAUUU |
| DNMT1$^{Δ1_15}$-sgRNA | GCCCUUCAGCUAAAAUAAAGGGUUUUAGAGCUAGAAAUAGCAAGUUAAAAUAAGGCUAGUCCGUUAUCAACUUGAAAAAGUGGCACCGAGUCGGUGCUUUUUU |
| RYR2$^{S404R}$-epegRNA | GCAUGAAGGCCACAUGGAUGAGUUUUAGAGCUAGAAAUAGCAAGUUAAAAUAAGGCUAGUCCGUUAUCAACUUGAAAAAGUGGCACCGAGUCGGUGCGACAAACGUAUGCCAUCAUCCAUGUGGCCUUCGCGGUUCUAUCUAGUUACGCGUUAAACCAACUAGAAUUU |
| RYR2$^{S404R}$-sgRNA | GAACGUAUGCCAUCAUCCAUGGUUUUAGAGCUAGAAAUAGCAAGUUAAAAUAAGGCUAGUCCGUUAUCAACUUGAA |

| | |
|---|---|
| | AAAGUGGCACCGAGUCGGUGCUUU |
| **Fig. 6d** | |
| NC-epegRNA | GACAUCGUUAGGGACAGCCUUGUUUUAGAGCUAGAAAUAGCAAGUUAAAAUAAGGCUAGUCCGUUAUCAACUUGAAAAAGUGGCACCGAGUCGGUGCGUCCAAAAAUUCUGCGUGCGGUCCACACAGUUGACAUGGUCGCGGUUCUAUCUAGUUACGCGUUAAACCAACUAGAAUUU |
| NC-sgRNA | GUCCCUGUACCGAUAGUGAAGUUUUAGAGCUAGAAAUAGCAAGUUAAAAUAAGGCUAGUCCGUUAUCAACUUGAAAAAGUGGCACCGAGUCGGUGCUUU |
| TRPM4$^{A320V}$-epegRNA | GCCUGGCUCCCCCACUCCCUGGUUUUAGAGCUAGAAAUAGCAAGUUAAAAUAAGGCUAGUCCGUUAUCAACUUGAAAAAGUGGCACCGAGUCGGUGCGAAGACACUCUGGUCCCAGGGAGUGGGGGAGCCGCGGUUCUAUCUAGUUACGCGUUAAACCAACUAGAAUUU |
| TRPM4$^{A320V}$-sgRNA | GCUGGAAGACACUCUGGUCCCGUUUUAGAGCUAGAAAUAGCAAGUUAAAAUAAGGCUAGUCCGUUAUCAACUUGAAAAAGUGGCACCGAGUCGGUGCUUU |

**Supplementary Table 2.** Oligonucleotide probe sequences used for sgGOLDFISH and refGOLDFISH.

| | |
|---|---|
| SCN5A$^{N406K}$ | |
| sgGOLDFISH | GACCACGGCCAGGATCAGGT |
| | GTTTTGCTCCTCATAGGCCA |
| | CTCGGTCTCAGCGATGGTGG |
| | CTCCTGGAAGCGCTTTTCCT |

| | |
|---|---|
| | TTCTTTCTTGAGCATTTCCA |
| | GTGGCACTGCTCACCCACCT |
| | AGGTGAGGGCTTAGAGGCTC |
| | CCCCACCTATAGGCACCTAC |
| | GGCACCAGCTTTAACTGGCA |
| | TTACCCAGCATGTACCACGC |
| | GCACCATGCCCACAGGCACC |
| | ACAGGCACTGGCTTTACCCA |
| | CCCAGCAGGCACTGCACCAT |
| | CGATACCACATTCACAGAAA |
| | AACCCAGCAGGTTCCAAATT |
| | GCTCCTCACTAACAGGCACG |
| | GCAGATGTCACTTTTACCTA |
| | CACTCATAGGACCCACCTTA |
| | GGAACAAGTTTACTCAGCAC |
| | ACCTGGAGGACATGACGACC |
| | CATTTTTACCCAGCTGGCGC |
| | CAGGAGCTACCACCCAGCAG |
| | ACTATGCTCACAGGGACCAT |
| | ACATGCATCTCTTATCCAGC |

| | |
|---|---|
| | TCCCTTGACCAGAATAAACT |
| | TCTTCACTACCCCACCAGAA |
| | GCTCTGTTCTCTGACATTGG |
| | TGTCAGTGCTTATTTTCTGA |
| | GGGGGACTAGGGATGGCTAG |
| | TATAGCCAGCCTCCAGGCTC |
| | GCTCTGGCTTCTAATAGCTC |
| | AAGAACAAGTGGATGCTTCT |
| | GAATATGTGGCATGGGGTAG |
| | AAGGTTTAGCTGGCCAAAAG |
| | GGCAGAACCAGAAGGCAGAG |
| | GGTGGGTGTAGATGAGCAGA |
| | ACTGGAGTGTGTGGAACAGG |
| | GTGGCTAAAGTGGGCAGCCT |
| | CAAGATCCAGGAAGGAAGAT |
| | GACCCAGAATGTTTTGGGAA |
| | TGAGGACATCATGGAGATAA |
| | TGGACTTGCACACCTACCCC |
| | TGACTTGGTTTTCCCTATCA |
| | CCTCAGGCTCAGGCAGGCCA |

| | |
|---|---|
| | TTCCATCATTGAGACTTTGT |
| | GGAAAAGAGTGGAGGAACGG |
| | CCACACCCCTGATGGTGAGG |
| | CCAAGGAGCTACGGGACACG |
| | GTTTACTGGGGCCAAAGGGG |
| | CCTCTTGCTTCTTCTCTCAT |
| | AGTTCCTGAAGACATCCGTT |
| | GGAGCCTGTCCTCCCCACAC |
| | GGGACCATCTTCTGAGTCAG |
| | CACAGCTGGGATTACCATTG |
| | GACCCACCCTGGAAAAGCTA |
| | TAGCATGATGGGATTTTCAT |
| | ATGGGGGCTGTGACTGTACA |
| | ACAGGAAGCGCAGAGATGAG |
| | CTACTCTAAGGAAGGCCATG |
| | TCCCTCTGCCCAGAGCTCCC |
| | ATCACAACACCCAAAACTAC |
| | GGATACTGGCCTCAGGTTGG |
| | ATGGGGTCTCGTCCCCCAGC |
| | CTTGAACTCCTACCAACCCA |

| | |
|---|---|
| | TGGACCCTCACTCTTATGGG |
| | TCCTGGTGTTGCAGACAGGC |
| | CAAGGGAGAAGGAGGCAGGA |
| | GCTTGGAGTTAGAGCTCAGG |
| | GCTGGGAGCCTGAAACTTAA |
| | TCCCCTCAAATCTATGTCTC |
| | ATCCTGTATCTTCTGGGATG |
| | AAGAGGGGATGTGGGGTTAG |
| | CCACACAGGAACCAGGTGGG |
| | CAGAGCATTGAGGGCTTTTT |
| | GCCACTGGCCTCAGCTAAAC |
| | TCATGTGCTCCCAGCCTGTC |
| | GGACACGACCATGACATGTG |
| | TGATTCTAAGGGCATGAGGA |
| | AGAAGTCCTGCTGAGGCCAC |
| | AAAATGCTCCCGCGGCTGGA |
| | TCAGAACCCAGGTCTCGCCT |
| | GCTGTTTTCATCATCTGCAA |
| | CAGTGATGTGTGGTGGCTCT |
| | CCACCCCATTGCAGTCCACA |

| | |
|---|---|
| | TTCCTGGGGATGTGGCCTCT |
| | ATCACAGGGCGGAGGAGGTG |
| | TGGGAGACCAGACCTGCTGT |
| | TTGGACTTGGCACTGGTGAT |
| | CAGTAGGTGCTCAACAAATA |
| | GCACCTAGAGCCCAGCACAC |
| | TAGCATGGATGCTCTATGAG |
| | TGAGTTTTGTCTGTCTGTCC |
| | TGAGCCATCGCACCTGGCTT |
| | CTGACCTCAGGTGATCTCCC |
| | ACCATGTTGGCCAGGCTGGA |
| | GGGATTACAGGCACGCGCCT |
| | ATTCTCCTGTCTCAGCCTCC |
| | AACCTCTGCCTCCTGAGTTC |
| | AATGGCATGATCTCAGCTCA |
| | ACTCTGTTGCCCAGGCTGAA |
| | CATCCCTCTTTATCACCATT |
| | GGCTTTACTAGTTGTGACAT |
| | AGGCTTCCCCAGATGATGGT |
| refGOLDFISH | GCGAGAGTCTGAGTGGCACA |

| | |
|---|---|
| | CCTGTACCAAGTCCCACAGA |
| | TTGAGGCACAAAGAGATTAA |
| | GCTGGCGTATAATTTTCACT |
| | CTCTTTCAGTCCTCACCACA |
| | GTCTCAACACTGGACCTAGT |
| | GGCTCTAAGACCAGGCAAGT |
| | CCTTCCTGCTTACTAATACC |
| | GCAGTGTACCCCAGGATCCT |
| | GGAGTGTGGGGCCCTGGGCA |
| | CCTCCCTTCACCCTGTGCTC |
| | GGTCTGCCCCCAGGGGTGCC |
| | CTCCCCAGCTCTACCTAAAG |
| | GACACCCCACCCTTCAGATG |
| | CTGGTTCAGTGTTCCTGGAA |
| | GAGGCTGAGGCTCCAGGGCT |
| | TCCCAGCTGCTGACCCGATT |
| | TGCAGTACAATGAGATGATT |
| | TGAACTTGGATTCCTCCCGA |
| | AGTCAGAATTCCTGGTAATG |
| | CCCCACAGAAGTGAGAATGG |

| | |
|---|---|
| | CTGCTAATCCCTTTAAGGTT |
| | CTTTGCAAATGACCCTGTCT |
| | TCTTGGTTTGCTAATTCCTT |
| | CCTGCTAATCTCTTTGACTC |
| | ACAGAAGGCATTCCCAGGGG |
| | TCTCACTGGGGACCCAGTGG |
| | TCAGACTAGAACTGGGCAGA |
| | TGTGCCCAACCCTGTGCTTT |
| | CCCATCCTTGTTGATTAGGC |
| | CTGAACGTGAGCTGTCTGCT |
| | CAGCCCCTACCTCCCAATCT |
| | CAAGCAAGGGGCAAAGTAAG |
| | CTTTTGGAACCAGGCAGTCT |
| | GTGCTTCTCCCACCTCCATC |
| | GGGGAGAAAAGGAATGGATA |
| | CAGGAAATGAGGATGAAGAT |
| | TGTTCCTAACTTGCTAGGTG |
| | AAGGCTTGGGTTAGAGTTCC |
| | GGATGGATAAATTCTTGGGA |
| | GGCTGGCACTAGATAGATAT |

| | |
|---|---|
| | CTCAACACTGCCTGGCACGC |
| | TTGTTCACTGCTATAACCCT |
| | TCCAAGAGACAGTGATCTTG |
| | TGTCTTCCTAACTACGGTAT |
| | CATGACTTATTGCTACCTGA |
| | CCCTCCCCTTAGCAGCTTTC |
| | GGCCTTTCCTGAGTATTCTA |
| | CCCTAAAATCATTTGCCTGT |
| | TCCTCTGCATCTTTAGGGCT |
| | CTATTCCCTCTGCCTGAACG |
| | TCCTCCTCAGGGGCTTTGCA |
| | TAGTCCCTCTCATCCCTTGG |
| | ATGTTTCACCACACCTGGCC |
| | AGTCCAAATCCTAACTCATG |
| | CCCATCTTACTCACACAGTA |
| | ACAACTCCTCCCACCCTACT |
| | CAGATAATGTCACTCCTCTG |
| | GCATAGTAGAAAGGGATTCC |
| | TCTTCTTCTCATCCCTGTAT |
| | AACAGCCTCCTAACTGGCTT |

| | |
|---|---|
| | TCTTCTCTTTCCCAAAACCA |
| | ACACCATCCTAGTCCAGGCA |
| | GGTAGTTTGTTACACAGCAT |
| | CTTGTTGCTAGAGATTGACT |
| | CAGATAAGTGAGCATGAGAA |
| | CTACATGGGCCATGTTGTAG |
| | GCAGGGCTTCCCAGCTAAGC |
| | AACCCACAGCATGGAGCCCA |
| | AGAACAAAGATGCAGAGCAG |
| | TGTCACCTTTTCAGCCTAAG |
| | ATTCTCCATGAAAAGAGCTT |
| | TTATCTTGCTCTTGTGCTCC |
| | AAACCTCAACTGGGCTAGTG |
| | TGTTAGTGGACATGCTGTGA |
| | AATGACTTGCTTTGGACAAC |
| | GAATTAACTTCCCTGCCCTG |
| | TCTCTTTCCCCTCCCACCGA |
| | GGTTATCACTTGGCACATAT |
| | CCCTCCTTCTTTGCCATCTA |
| | GCTTAACTCACAAGCAAGTT |

| | |
|---|---|
| | CCAACGCAACCATGAATTTG |
| | TGGTGCAATGGTACTACCCT |
| | AGTGGAGGCACCTGAGAGGG |
| | AGCTCCCAGTCATGGAAGTA |
| | CTGAAATGTTGCCTAGGGAG |
| | ATCCCGGCCATTGGTAGGAC |
| | TAGATCTGGGCTCAAGTTAA |
| | AAGCAGGGCATAGGGTGGAT |
| | GGCCTAGATTAGCCAGGGTA |
| | TCTTCATAGGGCAGAGGGAA |
| | CATGCAATAGCAAGACTCTA |
| | TTCACTTCAGCAGGGCCCTG |
| | CCACAGAACCAAGGAGGGCC |
| | CTCTAGGCTTACCTGGAGCC |
| | GGAAACAGCCCTGGTGAAGG |
| | GCATGCACCATTTTTATCAG |
| | CACTGTGGGGTTTGTTTTCA |
| | CCCTTTTCACTTCATGTACT |
| | GGAGGGGAAAAAGAGGGAGA |
| | ATATCTGGGGAATGGGATTG |

| | |
|---|---|
| | TCTGGAAAATGCATACCAAA |
| | CCAAGACAAACAGTTTGATG |
| | CTTTGGATGCTCCCATGCGG |
| | GACAACAAAGAGAACGAGGC |
| | ACAGTCCAACTCTGGGCTCT |
| | ACCAATCAATAGCAACCATA |
| | AGCAGTGTTCCTAATACCAT |
| | AAGTCCTCGTCTAGTAATCT |
| | GGCTGGATTTACCAAGAAAA |
| | CTGTGTCTTTTGTGGAGGCA |
| | TTACATATGGCTGGTGGGGG |
| $RYR2^{E4950K}$ | |
| sgGOLDFISH | TGTTGGGAATTTTTCCCAGC |
| | GCAGGAATCTTATGTCTGGA |
| | CACACTTTGGGGAAAATGTT |
| | CGTGCGATATAGGTAACAGA |
| | GAGACGTTAAGGTCTGGTAT |
| | GCCCAGCCAACATATTTTTA |
| | GCCTGAACTCAAGCGATCCT |
| | CTCTTATGTTGCTCAGGCTG |

| | |
|---|---|
| | CCTGAAACTACAGGCATGCA |
| | CCTGTTTCAGCCCCACAAGT |
| | TTCCTGGGCTCAATAAATCC |
| | AGGACCTACTACAGCCTTGA |
| | CCAAGTTCAAGTGGCATGAT |
| | ACCCTCAGAAGCATAAATGT |
| | CCACTGTGCAGATTGTTAGT |
| | TGCCTAAGGTGTTTAAGCTA |
| | TGCTGGATTTGAGTTTTGAA |
| | TGGAAAAGATTGCCTTTGTT |
| | GCACGGAATTACTTTCAGAT |
| | ATGTGTGCATTAAACAGTGA |
| | ACAGGACAGGTAGGTAAATT |
| | CCGTTCATTTCTGATCAGTT |
| | GCCATAATTGATTGCAACCA |
| | TGTGGCGTTTTCTCTGTGCT |
| | TTTCCTCTTTCTGGTACACT |
| | AGAAAAGTCAGGTCCACAGC |
| | AGGGTCAAATAGCAACAGAT |
| | TGACCAGTGTGTATATTCCT |

| | |
|---|---|
| | TCAGTGGAACTTGATCAGTA |
| | CCGAGTAGCATTATAGGGAA |
| | TTTTCCGACCCAATTTTAGT |
| | ACTCGAGATTAGCCAAAAGT |
| | ACCAACTTAAAAGCTGCATT |
| | TCTCGTCAGGCACTGTTGGA |
| | GAGTCTCCGTTTTTCTTCTT |
| | GGCTGCTTCCTAGGGAGATG |
| | GTCTTGGTCATGTTTCTCAT |
| | AGGAGCAGGGCCTCTGTGGG |
| | GAAATGAAGAGGTGGACAGG |
| | GCTCTTCATGTAAGTGATGC |
| | CTCTGGAGTAGGTTGACTTA |
| | ACAGCTAATGGCTCTCTTTT |
| | CCCATCTCAGAATAGAGAGG |
| | AGTAGACGCCACTGGTCCCT |
| | TTCAGAATCTTCCACCTCTC |
| | ATTACTTGTGAGTGTGCCCG |
| | TACAGGAGCACAACTTGGCT |
| | ATGGCTTTGAAACCCACACT |

| | |
|---|---|
| | ATTACTTCGACACAGTGCCA |
| | TCATCTGTGGGATAGGCAAT |
| | CCCTTTGTTTCTAGACCAA |
| | GCTTTCTGAACTCTGACGTT |
| | CGAGTTGTGTTTTCCTTTTG |
| | ATAATAGCTGCCCCTACATG |
| | GACGTGTATTGAGTGAACCA |
| | AGCCTCCATGTGATGATCTT |
| | TTATATTCGTGGGCTTGGCA |
| | TAGTTACAGCGACAGTTCCA |
| | ATTTAAGCCAACACCAAGAT |
| | CTTGCTGCTGGTACCTGTGT |
| | CCCTGTCTGCGTCTGCTACT |
| | GCCATCATCATCATCATCTT |
| | AGGAAGAGACGTTGCTTTTC |
| | ACATTTTCACCTAGGCTAGA |
| | AGAGTTGATCACTAAGCCTG |
| | AGACACAGTTAATCCTGCTT |
| | GGGTGGGTACTAAGTTTGTA |
| | CTTGGTCGTCCGACTTTGTT |

| | |
|---|---|
| | GGCCTTTTCTAAAGGAAAGT |
| | GGATGTCTCATAGCTTGGTA |
| | ACCCTTGCTAAACTGTAGAT |
| | TACCTAGCACAGCACCTGGA |
| | CACATCCACTGGCTAATTCT |
| | TGCCGAAAGTTGTTCTCCAT |
| | GACCAGGAATCTGGTCCCTA |
| | CAGAGAAGGTGGAACCAAGA |
| | GTGCTTAGGAGAGGCGTGGT |
| | AAGCAGGTGCCCCTCTACAG |
| | CTGTCTGCAAGGGGCAGACC |
| | CTTGGGTTTTGTGGACCAGG |
| | AGAGTCACATGGGACAATTA |
| | TGGCAGGGAAGGATGAGCAC |
| | TGGCATAAAACAGCTTACAA |
| | GGTCTCATAAACATGCTTCA |
| | CAGTAAGGCAGGAAGAATTT |
| | CAGCTGTCAGCAGTTGTTGA |
| | CAGTTGCTCAATGTGACCTA |
| | GACCTGATCGATTGTTCTCC |

| | |
|---|---|
| | ACATCATTGGAGTGGTATGA |
| | TGATTTGAGCCACTTCAATT |
| | CATCGGTAAAGCTGATGAGT |
| | AACCCAGAGGAAGAACAGGT |
| | GCATAGTTAGCTGACCTAGG |
| | TTAGCCACCAGATGGATGGT |
| | GGGATATCCAGGCTTCAGGC |
| | GGAGAAGATGGAGCAAGCAA |
| | GGACTTGGTGACTGACCAAA |
| | AGTTTCTGCATCCTCAGCGA |
| | AAGTACTGCCTGAATCCATC |
| | GGTGGCACATGGGAGTTGGG |
| | GAGATTCTGAAGCTTTTGCT |
| | CTGTCAGCATTTTCCAGGTG |
| | GAACAGGGACAGGGAAGTGG |
| | GCATGAAGTAGAAACTGGCA |
| | CCAGATGATCAGATTTGCAT |
| refGOLDFISH | AGTTTCTGCATCCTCAGCGA |
| | AAGTACTGCCTGAATCCATC |
| | GGTGGCACATGGGAGTTGGG |

| | |
|---|---|
| | GAGATTCTGAAGCTTTTGCT |
| | CTGTCAGCATTTTCCAGGTG |
| | GAACAGGGACAGGGAAGTGG |
| | GCATGAAGTAGAAACTGGCA |
| | CCAGATGATCAGATTTGCAT |
| | AAAGGGCTTTAAGCAGAGCA |
| | TGTGGGCTGTGGGAAGCAAG |
| | AGAGGAATACCAGTGAATGA |
| | AAATTCTGATGATGAGGCTG |
| | CAGTGCTACCAGATGCAGAT |
| | AAGTTGATGTGCAGGTTTTT |
| | CCCATCACCACCTGAAATTA |
| | ATTCACAGTGGGACCTTTGA |
| | ATGGACATCTCCAGGGGTAC |
| | CCCAACCCCTTCTCCCACAC |
| | GGAGGACAGTTTGGGCAACC |
| | TGTGGGTTTGATAAGGAGAG |
| | GGCCTGAAGACTGTGGAAGT |
| | AAGAAGGGATGTCTGAGCTG |
| | GATGGTCAGGGAATTTCCTA |

| | |
|---|---|
| | CTGGTATCTGATTCACTCCT |
| | GGAGTGGTAGGGTCTCTAGG |
| | CCTTTATGGGAGTAAAGCCC |
| | TTACAGAGCAGTGTGGTGAG |
| | CCAGAAGTGGAGAGACTATT |
| | ATAGGACACCATCCTTTCCC |
| | GCCACATGCATGCAAACAGC |
| | TCGGCTGAAATTGTGGTGGA |
| | ACATATAGTTGAGCTGCGTG |
| | CGTGTTTTCCATTCAAGCAA |
| | CACAACTTGTGGTTTCACTG |
| | GGGAAGGAGGAGGGGGAAGG |
| | TCGTTTTCCTCACTACCTTT |
| | CTTCTTTGTTGTCCTGGGCC |
| | GCTTCTCCATTCATGACTCA |
| | TCAAAGAAGACATGGAGGTA |
| | TAAGAGACCAACAGGAACAA |
| | GGATTAGCCAAGGAACTGAA |
| | ACATCTCCCCTTTCCCTGGG |
| | AGCATTTTGCAAGGTAGTTG |

| | |
|---|---|
| | AGTCACGTTTACCATGTTCT |
| | TCTCCTAGGAGTATCGGATA |
| | AAGACAGAACACCGCCTCCT |
| | CTGCACATACTTCTGTCCCC |
| | AGTGAGTGTATGTCCCAGTA |
| | CCTGCTGTCACTTCATTTTA |
| | TGCCAAGTTAACAAAAGGTT |
| | TTTTTCGGCATGAAGTATGT |
| | TCGTTTAAGTGGTGATGTTT |
| | AAAGACTTGAGGCAGATTTC |
| | ACTTGACTGGGCATGGTGGC |
| | TCAGGGTGTGGTGGTGTGCG |
| | CCTGTCTGTACAAACAGAAA |
| | GCCTGGTCAACAAGACGAAA |
| | GGGGCCTGGCACCATGGCTC |
| | CTGATTGTCACAGTCAATGA |
| | AAACAGCTCAACACAGTGTT |
| | TCAGGAGGGAAAAGGAGCTT |
| | AAAGTGTGAAGCCAGCAGGT |
| | GGGCAAAGTTATAACTTCGA |

| | |
|---|---|
| | ACAAGCTTAAGGGACTCATA |
| | CACAGACATCAAATGACACT |
| | CATGTATCCTGGCTTGAATT |
| | GTCATGTTATCTTTTCCGCT |
| | TGGTGTTTTTCTCTCTCAGA |
| | ACTTCCGGGCAACTTGGAGT |
| | GACCCACTAGCTCTTCCTTT |
| | CTTCAACCACGGTGCTGTAT |
| | AGGCAATCTGCATCCAGACC |
| | GGCTGGAATCCATCTATACC |
| | CCAAAGTGCACAGCTAAACA |
| | AGCCACAATAAGTCTAAGGA |
| | CCTTATGAGAATGGTGCTCT |
| | GCCCAGCACATGTATTAACT |
| | GATGCATAGTCAGTGCTTTA |
| | ACATTCAAGTACCCAGTCTA |
| | AAAACCACCATAGCACGTGT |
| | GATGACAAGTTGATGGGTAC |
| | GGTTGGGATAGCATTAGGAA |
| | GGCATCACACACCAGAGCCT |

| | |
|---|---|
| | AACAATGGTAACACATTGGG |
| TRPM4$^{A320V}$ | |
| sgGOLDFISH | TGCCTGGCTCCCCCACTCCC |
| | ATTCGATCTCGGGCTTCGCC |
| | CCTTTGGGAAAGAAACGCCT |
| | GCCTGCAGGACCTCAAGGTC |
| | GGCCCCCAGTGTCATACCTG |
| | CCATCTCAGGATCCAGAGTT |
| | GCCCAAGTCCCCAGTTCCCT |
| | TGGAAGTCCAGGCCCCCAGC |
| | GCACCCTCTTCTCCCGGAAC |
| | CCTCCAGGACCTAATAGTCT |
| | AGGGATGGAGGAGGCCATTC |
| | CTCCTGTTGGGCCCTGTCCA |
| | CTAGCCCGCCTCCATCCCCA |
| | GGAATAGCCAGTTCTGCATC |
| | GTCATAATCCTCTCCACCTG |
| | ACTGTCAGGAGCTCCTTCCG |
| | GACCCATCCTCAGAAGAATA |
| | ACTATGGTCTCGAATTCCTC |

| | |
|---|---|
| | CCCTTCACAAGGGCCTTCAA |
| | GACTGGAGGGTACAACTTTT |
| | GTTGAACTGAGAGAGGGGGA |
| | AGGCCAGGAAAGGTGTCTAA |
| | GAGAGGGTGGATGAAGTGCC |
| | GGGGGCATGGGCCTTGGATT |
| | CCCCAAAGGGGAAAAATATT |
| | CCCAAATAGCACCATGCAAA |
| | TGAAAGCATCTGGGAAAACT |
| | AAAGCAGATGCTGGAACATA |
| | GATGGATGGATCCAACAGAG |
| | TGTGGATGGATGGAAAGATG |
| | AGATGGAAAGATCGATCTGT |
| | CTGGAAGGATGGTAGGTAAA |
| | GGATGAAAGAATGAGGGATG |
| | TGGACAGGTGAAAGGATGGA |
| | CTGAATGGAATTGTCAGTGG |
| | GGAAGAATGAGCAGATGGCA |
| | TAGATGGACAGGTGAGAAGA |
| | CAGGAATTTTCGGTGGATGT |

| | |
|---|---|
| | GGAATGAGTGGATGGAATTC |
| | GGATGGCATTCTCAGGAATT |
| | GATGGAATTGTCAGTGGATG |
| | AGATGGATGGATGGAACAAT |
| | TTCAGTGGATGTGTGGATGG |
| | AGAAGAATGAGCAGATGGCA |
| | CAGATGGACAGGTGAGAAGA |
| | CAGGAATTTTCAGTGGATGT |
| | GGAGGAATGAGTAGATGGAA |
| | TGGATGGAGGAATGAGCGGA |
| | GGAGGAATGAGTGAATGGAA |
| | AATGGATGGAAGATGGGTGG |
| | AAGATGGATGGGTAGATGGG |
| | ACAGTTGGATGGATGAATGA |
| | AGGAAGAATGGGTGGGTGGA |
| | GCCAGAGTGCTTCATGTATA |
| | CAGTGCTTTGGTTCATGTCT |
| | CTGGACTTCCTCACTGGGAG |
| | GCTGCCATTTCTAGTGAAAT |
| | TTTCATCATTCCGTCCTCCA |

| | |
|---|---|
| | ATTATGCTCCCCCGCCTTGG |
| | CCCAGCCTCTCCCAGGTGTG |
| | GACCCTGTCTCAATAAGTGA |
| | TACTCCAGCTTGGCCTACAG |
| | GTATGTTCCCAGCTACTTGG |
| | ACCAGGTGTAGTGGTGGGCG |
| | GAGCAAGACCTCGTCTGTAC |
| | AGGAGGCTGGCATGGTGGCT |
| | GGAGCCTAGGATTGTATTTG |
| | AATGGGTTGGCTTTAGTCTC |
| | CTGGGGTCTTTGAGAGCAGG |
| | TCCAAACAATCCTAGTGGGA |
| | CCTGTGCTTCTGTCTTAAAT |
| | CCTTCTAACGACAGTGATTC |
| | ACACAACTTTTCCTTGAAGT |
| | CAGCCGGGCATGATGGCACG |
| | ACTTCTCTGGCCAGGTGCAG |
| | CAAGGTCCGGTTTAAATGCC |
| | TGCTGAAATTACACGTGTGA |
| | TATAGAGGTGGCATTTCACC |

| | |
|---|---|
| | ACCCTCCCCAACTTATTATT |
| refGOLDFISH | CTTTCCACAGAGAGAAACTG |
| | GAGGCTTATAACACTCCTGT |
| | CACAACTCATACCTTGCCAT |
| | TGGGCCCCCACCGGGACCGG |
| | CCCAGAGGAAATGGAATCAT |
| | CCCACTGAGAGTTACCTTAT |
| | CCGCAGAGTGAATCAGATTT |
| | AGGAGGCTGCTGAGTTAACT |
| | AGAGCTGCCGTATTTTCTCC |
| | CCCCCTCAAACCACATCTTG |
| | CCTGCTGGGCTTACAACACC |
| | TGGACTGTGATCACTTCCAC |
| | AGCCATCTCCAGTTCCCCCA |
| | TCTGCTATCCCCAAGCCTAT |
| | GCTGAAATTCCATCCATTCA |
| | ATGACCTTGCCCTTCTTTCT |
| | CCCGGGGGTTGCAGTGAGCT |
| | AGATCACGAGGTCAGAAGGT |
| | CATGGGCCAGGGGTGGTGGC |

| | |
|---|---|
| | TGCTAGACTCTTAGGAAGGA |
| | GCCCAGCACTGGGCCTGCCT |
| | GGGCAAAAGAGTACACCAAC |
| | CAGGCAGGGAAGGGAGATGA |
| | GCAAAATGGGGTGATAATCC |
| | CTCTCTCTGAGCCTCAGTCT |
| | ACCCTGGGCAAGCCATTTTT |
| | ACCCCTGCTACTAGCTGGAT |
| | CAGACTCACATGAAAAGCCT |
| | GCAATGGGCTTGAGAATTAT |
| | CTCTCTCCAAACATGCCCCT |
| | CCAACCCTGCGAGCCTCTGA |
| | CCTTCATGCCAATGATACAT |
| | CTGGCCTTGTCTCCAACACT |
| | GGTTTGGCCCTACCTGCCTC |
| | CTGGCATTTGAGGTCCTGGG |
| | GGACAGCCTTAGCCCCTCTG |
| | TGGTCCTCTGCTGCCTCTCA |
| | CTTGGCCAGAGAGTCTTTTT |
| | TAATAGGCATGAGCCTTCAC |

| | |
|---|---|
| | TAGTCTTCCAAGGTGCTGTG |
| | GCTCAAGCGAACCTCCTTCC |
| | AGTACCATACCCAGCTAATT |
| | GAGACCTCCCAGGCTCAAGC |
| | CATCACAGCTCACTTCAGCT |
| | CGTCTGTCACCCAGTCTGGA |
| | CCAGAGAGTCTGTTTTTGTT |
| | CCACTCTGCCCCCACCCACT |
| | CTGCAGTCTCTTCTCTCCTC |
| | CGGCACCCCCAAACACCATC |
| | CCCCAGCTATAGCTCAAGCT |
| | CCTCCCCGACTGTCCTCCTG |
| | GCCTCCTGACTCTCTGCTGA |
| | AAGCTTCATGTCCCACGTCC |
| | TCTCACAGGCCATCAGCCCC |
| | TGTCCCTTCCTGCTCCTCCC |
| | CAGACCCGCAGGTACTGTCC |
| | CTGTCCACCCAATCAATAAG |
| | CTCATCTCAGGAATGGCCCC |
| | CAAACCCATCTTCCCTACGT |

| | |
|---|---|
| | CTCAACTCAGCATCTAGCCC |
| | TCAGACTCACCGTGTTCCAT |
| | CCCTGGCCCCACCTTGTCCC |
| | ATTTGTCACCTGGGCACTCG |
| | ATTCCGCCTTAATCTGGCAC |
| | CGAAACTCTGTTGCAAAAAT |
| | ATAGGCTGGATGCTGTGGCT |
| | GAAAGGGAGACTCCGTCTCA |
| | ATGGGCGGATCGCCTGAGCT |
| | TACAGGCCGGGCGCCGTGGC |
| | CCACAGTTTCAAAAGAAGGA |
| | CGATTCTATGAAGCTCTCCT |
| | GGACTTTTCCACGACTTTCT |
| | ACCCAGACGTCCATTCAGAG |
| | GGCTTTCTGTGCTCTTTCAA |
| | GTAACATGTCTAGGACCTCA |
| | AAACAGAGGCCCAGAAAGGC |
| | CCTCCAAGGCTGTGCTTGAG |
| | ACTACAGCTTGGCCAGTGTA |
| | GAAGTTCCACGCTGTGGCGA |

| | |
|---|---|
| | CCCTTCATAGCACTGTAATC |
| | TAAATGGGAAATAGTGGCAC |
| | TTGGGAGCCTCAGTTTCCTC |
| | CTTCAGGCAAGCCATTGAAC |
| | CCACCAGTACCAACTGAGTG |
| | ACGTGGGTTCCAGTCCCCGG |
| | AATAGGCTCTGGGGCAGGCA |
| | TCTCATGTAGTGGAGGTGAG |
| | CTGGTACCACAGAAAGAGAT |
| | TCATTTTGGTTGCTCCAAAA |
| | CTCCACTTGCTCCATATCTA |
| | TGGAGTTAGGTTTCCTTCTG |
| | TGAAGGGCATTTGTCGTGAC |
| | CAACAGGGGCATGAAAAGTC |
| | GTGGATCCAAACATGTGCAG |
| | AGAGAACCAGAAACCCTTAA |
| | GGAGGTCACTGAAACAGCAA |
| | AGAGGGACTCCGAGAGACCC |
| | ACAGCAGAAAATGAGAACAG |
| | TGTGAAGGGAAAGAGGTGAC |

| | |
|---|---|
| | TGATTTCTGAAACTTGGGGT |
| | GATGACAAACGTGAGTCACT |
| | CCGGGTGCAACACACACACA |
| | AGGCACAGCAGAAGCAAAGG |
| | GAAAAGGGCGTTCAAAGCAG |
| | GGAGGTATGCCCCTAAGGTG |
| | TTTGAAAGACCACCAGGACT |
| | GAATGGGTGTTGAGGTAGCA |
| | TGGCCCAAGAGAGCTCCCAG |
| | GGATATGGTCACAGAGGTAT |
| | ATATGATTAGGGCCCAGAGG |
| | GGAGGCATCTATAGGCATAT |
| | GCAGGGTGGAGGGGTGTCTA |
| | CTGGACAGGCCATAATGGAA |
| | GATGGGCGATTAAAGGGAAG |
| | GCAGACGGACAATCAGAGAT |
| | GGAGAGAGGAAACAAGATGG |
| | GATAGATGGACAGACGGATG |
| | GATGTACAGGGTGGATAGAT |
| | TGAACGGATGGATGGATAGA |

| | |
|---|---|
| | TGGGATGGATAAATGGATGG |
| | ACAGATAGAAGGATGGATGG |
| | CCTCAGAGGTAGGCATATAA |
| | TACCCTCAGGGTGCTTGGTG |
| | TTGGGGGAAATATGTCCCCC |
| | ACCCTGAAGAAGCTGGTTCT |
| | GGATGGAGAGAGAAATCAGA |
| | TTATGGACCAAGCAGTGGGA |
| | CCCAAGAAGAAACAAAAGGA |
| | ACAGACCACTGGTTTACAGC |
| | CATTTGGCTCAGGGCCAGAG |
| | AGGCTATCATAGGCATAAGG |
| | TGTGGGGGTCTTAAAGCAGT |
| | AGGGGTTGCTACTGGGCAGA |
| | GCCTCTATTTTGCATGTCTG |
| | GGACATCTGCTGAATGTCTA |
| | GACAAGAAGTCAGGGGAGGA |
| | GGAAGGATTAGAGGGAGAGG |
| | GCCTCCGAGCTCCCACAGGC |
| | AGCTCATCCAGGTAGGCTGA |

| | |
|---|---|
| | TTCCAAGCCACAGCCAAACG |
| | CTCTGGGCAATGTCCACGCG |
| | CCCGCCATTGGATGTCCCCC |
| | CCCCCAGGCCCTGACCCCTC |
| | TCCCTTTGTCAGTATGCCCA |
| | CAGAGAGGGAGGACAGCACA |
| | GGATCACACCTCTGCACTCC |
| | CAGAGAGGCCGGGTGCGGTG |
| | GTTGAGGGGCGTCTAAGAAG |
| | CAGGCCATAATGGAAGGCAG |
| | GACAATTAAGGGAAGGCTGG |
| | GCAGATGGACAGAGATGGAT |

**Supplementary Table 3.** Templates used to synthesize sgRNAs for sgGOLDFISH and refGOLDFISH.

| SCN5A$^{N406K}$ | | |
|---|---|---|
| sgGOLDFISH | WT | TTCTAATACGACTCACTATAGGATCAGGTTCACCAGGT AGAGTTTTAGAGCTAGA |
| | Mut1 | TTCTAATACGACTCACTATAGGATCAGTTCCACCAGGT AGAGTTTTAGAGCTAGA |
| | Mut2 | TTCTAATACGACTCACTATAGGATCAGTTTAACCAGGT AGAGTTTTAGAGCTAGA |

| | | |
|---|---|---|
| | Mut3 | TTCTAATACGACTCACTATAGGATCAATTTCACCAGGT AGAGTTTTAGAGCTAGA |
| | Mut4 | TTCTAATACGACTCACTATAGGATCCGTTTCACCAGGT AGAGTTTTAGAGCTAGA |
| | Mut5 | TTCTAATACGACTCACTATAGGATAAGTTTCACCAGGT AGAGTTTTAGAGCTAGA |
| refGOLDFISH | ref1 | TTCTAATACGACTCACTATAGGTGGCACAAAATAAGC CACGGTTTTAGAGCTAGA |
| | ref2 | TTCTAATACGACTCACTATAGCTGTGCTCCAGAGGAGT GTGGTTTTAGAGCTAGA |
| | ref3 | TTCTAATACGACTCACTATAGGCTGCTGACCCGATTCC ATGGTTTTAGAGCTAGA |
| | ref4 | TTCTAATACGACTCACTATAGAGCACAGAAGGCATTC CCAGGTTTTAGAGCTAGA |
| | ref5 | TTCTAATACGACTCACTATAGATGAAGATTTAACATGT AGGGTTTTAGAGCTAGA |
| | ref6 | TTCTAATACGACTCACTATAGTTAATGTTACCTTCTTG GGAGTTTTAGAGCTAGA |
| | ref7 | TTCTAATACGACTCACTATAGAGTATATCCAAACTTAT CTAGTTTTAGAGCTAGA |
| | ref8 | TTCTAATACGACTCACTATAGATAAGTGAGCATGAGA ATAAGTTTTAGAGCTAGA |
| | ref9 | TTCTAATACGACTCACTATAGTGTGAACAGAAACCTCA ACTGTTTTAGAGCTAGA |

| | | |
|---|---|---|
| | ref10 | TTCTAATACGACTCACTATAGGGAGGCACCTGAGAGG GCTGGTTTTAGAGCTAGA |
| | ref11 | TTCTAATACGACTCACTATAGTGGAGCCTCCCACAGAA CCAGTTTTAGAGCTAGA |
| | ref12 | TTCTAATACGACTCACTATAGGATAGACAACAAAGAG AACGGTTTTAGAGCTAGA |
| RYR2$^{E4950K}$ | | |
| sgGOLDFISH | WT | TTCTAATACGACTCACTATAGTGTTGGGAATTTTTCCC AGCGTTTTAGAGCTAGA |
| | Mut1 | TTCTAATACGACTCACTATAGTGTTGGACATTTTTCCC AGCGTTTTAGAGCTAGA |
| | Mut2 | TTCTAATACGACTCACTATAGTGTTGGAACTTTTTCCC AGCGTTTTAGAGCTAGA |
| | Mut3 | TTCTAATACGACTCACTATAGTGTTCGAAATTTTTCCC AGCGTTTTAGAGCTAGA |
| | Mut4 | TTCTAATACGACTCACTATAGTGTCGGAAATTTTTCCC AGCGTTTTAGAGCTAGA |
| refGOLDFISH | ref1 | TTCTAATACGACTCACTATAGAGTTTCTGCATCCTCAG CGAGTTTTAGAGCTAGA |
| | ref2 | TTCTAATACGACTCACTATAGCAGTTTAATATGTGGGC TGTGTTTTAGAGCTAGA |
| | ref3 | TTCTAATACGACTCACTATAGCACACCATGGACATCTC CAGGTTTTAGAGCTAGA |

| | | |
|---|---|---|
| | ref4 | TTCTAATACGACTCACTATAGTAGATGATTACAGAGCAGTGGTTTTAGAGCTAGA |
| | ref5 | TTCTAATACGACTCACTATAGACAAGTCAAAGAAGACATGGGTTTTAGAGCTAGA |
| | ref6 | TTCTAATACGACTCACTATAGCTCCTTTCTCCTAGGAGTATGTTTTAGAGCTAGA |
| | ref7 | TTCTAATACGACTCACTATAGGGCAGATTTCTTCGTTTAAGGTTTTAGAGCTAGA |
| | ref8 | TTCTAATACGACTCACTATAGAATGACTTGACTGGGCATGGGTTTTAGAGCTAGA |
| | ref9 | TTCTAATACGACTCACTATAGTGTACAAACAGAAATTATCAGTTTTAGAGCTAGA |
| | ref10 | TTCTAATACGACTCACTATAGGAAGCCAGCAGGTCTCAGGAGTTTTAGAGCTAGA |
| | ref11 | TTCTAATACGACTCACTATAGTCCGCTTAACACATGTATCCGTTTTAGAGCTAGA |
| | ref12 | TTCTAATACGACTCACTATAGTTTCACCAACCTTATGAGAAGTTTTAGAGCTAGA |
| | ref13 | TTCTAATACGACTCACTATAGTGAGGGGTTGGGATAGCATTGTTTTAGAGCTAGA |
| $TRPM4^{A320V}$ | | |
| sgGOLDFISH | WT | TTCTAATACGACTCACTATAGTGGGGCCAGAGTGTCTTCCAGTTTTAGAGCTAGA |

| | | |
|---|---|---|
| | Mut1 | TTCTAATACGACTCACTATAGTGGGAACAGAGTGTCTTCCAGTTTTAGAGCTAGA |
| | Mut2 | TTCTAATACGACTCACTATAGTGGGACAAGAGTGTCTTCCAGTTTTAGAGCTAGA |
| | Mut3 | TTCTAATACGACTCACTATAGTGGGACCCGAGTGTCTTCCAGTTTTAGAGCTAGA |
| | Mut4 | TTCTAATACGACTCACTATAGTGAGACCAGAGTGTCTTCCAGTTTTAGAGCTAGA |
| refGOLDFISH | ref1 | TTCTAATACGACTCACTATAGCTTTCCACAGAGAGAAACTGGTTTTAGAGCTAGA |
| | ref2 | TTCTAATACGACTCACTATAGGCCATCTCCAGTTCCCCCAGGTTTTAGAGCTAGA |
| | ref3 | TTCTAATACGACTCACTATAGGATTCATGGGCCAGGGGTGGGTTTTAGAGCTAGA |
| | ref4 | TTCTAATACGACTCACTATAGGCTACTAGCTGGATGACCCTGTTTTAGAGCTAGA |
| | ref5 | TTCTAATACGACTCACTATAGTAGGCATGAGCCTTCACGCTGTTTTAGAGCTAGA |
| | ref6 | TTCTAATACGACTCACTATAGTCTTCGTCTGTCACCCAGTCGTTTTAGAGCTAGA |
| | ref7 | TTCTAATACGACTCACTATAGCCCAGCTATAGCTCAAGCTTGTTTTAGAGCTAGA |
| | ref8 | TTCTAATACGACTCACTATAGCTACGTCCTCATCTCAGGAAGTTTTAGAGCTAGA |

| | | |
|---|---|---|
| | ref9 | TTCTAATACGACTCACTATAGGCTTTACAGGCCGGGCG CCGGTTTTAGAGCTAGA |
| | ref10 | TTCTAATACGACTCACTATAGACCCAGACGTCCATTCA GAGGTTTTAGAGCTAGA |
| | ref11 | TTCTAATACGACTCACTATAGGAGGTGAGCAATAGGC TCTGGTTTTAGAGCTAGA |
| | ref12 | TTCTAATACGACTCACTATAGGGGACTCCGAGAGACC CTGGGTTTTAGAGCTAGA |
| | ref13 | TTCTAATACGACTCACTATAGGAAAGACCACCAGGAC TGGGGTTTTAGAGCTAGA |
| | ref14 | TTCTAATACGACTCACTATAGGGATAGATGGATGTACA GGGGTTTTAGAGCTAGA |
| | ref15 | TTCTAATACGACTCACTATAGGTATGTTATGGACCAAG CAGGTTTTAGAGCTAGA |
| | ref16 | TTCTAATACGACTCACTATAGCAAACGCAGCTCATCCA GGTGTTTTAGAGCTAGA |

**Supplementary Table 4.** Sequences of target genes cloned into the adenoviral vectors.

| >Fig. 1b CheRiff-HA |
|---|
| atgggcggagctcctgctccagacgctcacagcgccccacctggaaacgattctgccggaggcagtgagtaccatgcccc agctggatatcaagtgaatccaccctaccaccccgtgcatgggtatgaggaacagtgcagctccatctacatctactatgggg ccctgtgggagcaggaaacagctaggggcttccagtggtttgccgtgttcctgtctgccctgtttctggctttctacggctggc acgcctataaggccagcgtgggatgggaggaagtgtacgtgtgctccgtggagctgatcaaagtgattctggagatctatttc gagttcaccagtcctgctatgctgttcctgtacggagggaacattaccccatggctgagatatgccgaatggctgctgacatgt cccgtgatcctgattcatctgtctaacatcaccggcctgagtgaggcatacaataagcggacaatggctctgctggtgtccga cctgggaactatttgcatgggagtgacagccgctctggccactgggtgggtgaagtggctgttttactgtatcggcctggtgta |

tggaacccagacattctacaacgctggaatcatctacgtggagtcttactatatcatgcctgccggcggctgtaagaaactggt
gctggccatgactgccgtgtactattctagttggctgatgtttcccggcctgttcatctttgggcctgaaggcatgcacaccctg
agcgtggctgggtccactattggccataccatcgccgacctgctgtccaagaatatttggggactgctggggcacttcctgcg
gatcaaaattcacgagcatatcattatgtacggcgatatcaggagaccagtgagctcccagtttctgggacgcaaggtggac
gtgctggccttcgtgacagaggaagataaagtggcggccgccaagagctacccatacgatgttccagattacgcttaa

>Fig. 1b jRCaMP1b-Junctin

atgctgcagaacgagcttgctcttaagttggctggacttgatattaacaagactggaggaggttctcatcatcatcatcatcatg
gtatggctagcatgactggtggacagcaaatgggtcgggatctgtacgacgatgacgataaggatctcgcaacaatggtcg
actcatcgcgacgtaagtggaataagtggggtcacgcagtcagagctataggtcggctgagctcagcgaacaacaccgaa
atgatgtacccagcggatggtggtctgcgtggttacactcacatggcgctgaaagttgatggcggcggtcacctgtcctgttct
ttcgtgaccacctaccgctccaaaaagactgtcggcaacattaagatgcctgccattcattacgtcagccaccgtctggagcg
cctggaggagagcgataacgaaatgtttgtcgtacagcgtgaacacgcagttgccaagtttgtgggcctgggtggtggcgg
cggtaccggagggagcatgaactccctgatcaaggagaacatgcgtatgaaagtggttctggaaggctccgtaaacggcc
accagttcaaatgcactggtgaaggcgaaggcaacccgtatatgggcacccagactatgcgtatcaaagtgatcgagggtg
gtccgctgccgtttgcgttcgacatcctggcgacgtcctttatgtatggctcccgtaccttcatcaaatatccgaaaggcatccc
ggatttctttaagcagtccttcccggaaggttttacctgggaacgtgtgacccgttacgaagacggcggcgtaattaccgttat
gcaagacacgtctctggaggatggctgcctggtgtatcacgtgcaggttcgcggtgtgaacttcccgagcaatggtgctgta
atgcaaaagaaaaccaaaggttgggagcctacggactcccaactgactgaagagcagatcgcagaatttaaagaggctttct
ccctatttgacaaggacggggatgggacaataacaaccaaggagatggggacggtgatgcggtctctggggcagaaccc
cacagaagcagagctgcaggacatgatcaatgaagtagatgccgacggtgacggcacaatcgacttccctgagttcctgatt
atgatggcaggcaaaatgaaatacacagacagtgaagaagaaattagagaagcgttcggcgtgtttgataaggatggcaat
ggctacatcagtgcagcagagcttcgccacgtgatgacaaaccttggagagaagttaacagatgaagaggttgatgaaatg
atcagggaagcagacagcgatggggatggtcaggtaaactacgaagagtttgtacaaatgatgacagcgaagGGTGG
AGGCGGTTCAGGTGGCGGCGGATCAGGAGGTGGCGGATCGtccggactcagatctatg
gctgaagataaagaggcaaagcacggaggacacaagaatggcggagaggaggaatttccggagggtcctttttcacgtg
gttcatggtcattgcgttgctgggcgtctggacatcggtggccgtcgtgtggtttgacctcgttgattatgaggaagttctagga
aagctaggagtctatgacgcagatggtgatggggacttcgatgtggatgatgccaaggttttattagaaggacctggtgggtt
agccaagaggaaaactaaggctaaagttaaagaacccaccaaagaagagctcaagaaggagagagagaaagctgtgcct
agcaagactgaagagggaagaagggggaagaaggagcgggaggatggaaggaaggagaagaaaaagtctgactcgg

| |
|---|
| acatatcccagaaagcgtctcctggaggtaagagagacagagcgaaggagaaagccagctcagataaaagtgggaagtc<br>caaggaaaccgtgaaaaaggcagcagagacaaagtccgttcctagtaaagtggcttccacagacaaagacaggaaagga<br>agaggctcctccagccatgctccagtcacaaaggaaaatagccagaagagaaagaactga |
| >Fig. 1i Cepheid1b-CheRiff |
| ATGGCTGACGTGGAAACCGAGACCGGCATGATTGCACAGTGGATTGTCTTT<br>GCTATTATGGCTGCTGCTGCTATTGCTTTTGGAGTGGCTGTGCACTTTCGGCC<br>TTCAGAGCTGAAGAGCGCATACTATATCAACATTGCCATCTGCACTATCGCCG<br>CTACCGCTTACTATGCAATGGCCGTGAACTACCAGGACCTGACAATGatggtgag<br>caagggcgaggcagtgatcaaggagttcatgcggttcaaggtgcacatggagggctccgtgaacggccacgagttcgaga<br>tcgagggcgagggcgagggccgcccctacgagggcacccagaccgccaagctgaaggtgaccaagggtggccccctg<br>cccttctcctgggacatcctgtcccctcagttcatgtacggctccagggccttcaccaagcaccccgccgacatccccgacta<br>ctataagcagtccttccccgagggcttcacctgggagcgcgtgatgaacttcgaggacggcggctccgtgaccgtgaccca<br>ggacacctccctggaggacggcaccctgatctacaaggtgaagctccgcggcaccaacttccctcctgacggccccgtaat<br>gcagaagacaacaatgggctgggaagcgtccaccgagcggttgtaccccgaggacggcgtgctgaagggcgacattaag<br>atggccctgcgcctgaaggacggcggccgctacatcgcggacttcaagaccacctacaaggccaagaagcccgtgcaga<br>tgcccggcgcctacaacgtcgaccgcaagttggacatcgtctcccacaacgaggactacaccgtggtggaacagtacgaa<br>gcctccgtgggccgccactccaccggcggcatggacgagctgtacaagAATGGTGAAAGGCAGGTGGT<br>CTACGCAAGATATATTTGTTGGGTGCTGACCACACCACTGCTCCTGCTCGATC<br>TCATCGTCATGACCAAGATGGGCGGAGTGATGATTTCTTGGGTCATCGGCGC<br>AGACATTTTCATGATCGTGTTTGGTATTCTGGGCGCCTTCGAGGATGAACAC<br>AAGTTCAAATGGGTGTACTTTATCGCTGGATGTGTGATGCAGGCAGTCCTGA<br>CATACGGGATGTATAACGCCACTTGGAAAGACGATCTGAAGAAAAGCCCCG<br>AGTACCATAGCTCCTATGTCAGTCTGCTCGTCTTCCTGTCAATCCTCTGGGTG<br>TTTTATCCTGTCGTGTGGGCTTTCGGGTCTGGTAGTGGCGTGCTGTCCGTCG<br>ACAATGAGGCCATTCTCATGGGAATCCTGGATGTGCTCGCTAAGCCACTGTT<br>TGGAATGGGGTGCCTCATTGCCCATGAGACTATCTTCAAGaccggtgccgccgaccgc<br>ccggtagtagcagtgagcaaggcggccgccGCTAGCaaatcaaggatcacctctgagggcgagtatatccctctgga<br>tcagatcgacattaacgtcggtggaaaatcccgcataacctctgagggcgagtacatccctctggatcaaatcgatataaacg<br>tggggggaaaaagtcgaatcacttctgagggcgagtatataccccctcgatcagattgacatcaatgtgggggggcAACAT |

GGCCATCATCAAGGAGTTCATGCGCTTCAAGGTGCGCATGGAGGGCTCCGT
GAACGGCCACGAGTTCGAGATCGAGGGCGAGGGCGAGGGCCGCCCCTACG
AGGGCTTTCAGACCGCTAAGCTGAAGGTGACCAAGGGTGGCCCCCTGCCCT
TCGCCTGGGACATCCTGTCCCCTCATTTCACCTTTGGCTCCAAGGCCTACGT
GAAGCACCCCGCCGACATCCCCGACTACTTCAAGCTGTCCTTCCCCGAGGG
CTTCAAGTGGGAGCGCGTGATGAACTACGAGGACGGCGGCGTGGTGACCGT
GACCCAGGACTCCTCCCTGCAGGACGGCGAGTTCATCTACAAGGTGAAGCT
GCGCGGCACCAACTTCCCCTCCGACGGCCCCGTGATGCAGAAGAAGACCAT
GGGCTGGGAGGCCTCCTCCGAGCGGATGTACCCCGAGGACGGTGCCCTGAA
GGGCAAGATCAAGATGAGGCTGAAGCTGAAGGACGGCGGCCACTACACCT
CCGAGGTCAAGACCACCTACAAGGCCAAGAAGCCCGTGCAGCTGCCCGGC
GCCTACATCGTCGACATCAAGTTGGACATCACCTCCCACAACGAGGACTAC
ACCATCGTGGAACAGTACGAACGCGCCGAGGGCCGCCACTCCACCGGCGG
CATGGACGAGCTGTACAAGcagagccagcctatcctgaacaccaaggagatggcccctcagagcaagcc
tcctgaggagctggagatgagcagcatgcctagccctgtggcccctctgcctgccaggaccgagggcgtgatcgacatga
ggagcatgagcagcatcgacagcttcatcagctgcgccaccgacttccctgaggccaccaggttcTTCTGCTATG
AAAATGAAGTGggctccggagccacgaacttctctctgttaaagcaagcaggagacgtggaagaaaaccccggt
cccggcggagctcctgctccagacgctcacagcgccccacctggaaacgattctgccggaggcagtgagtaccatgcccc
agctggatatcaagtgaatccaccctaccaccccgtgcatgggtatgaggaacagtgcagctccatctacatctactatgggg
ccctgtgggagcaggaaacagctaggggcttccagtggtttgccgtgttcctgtctgccctgtttctggctttctacggctggc
acgcctataaggccagcgtgggatgggaggaagtgtacgtgtgctccgtggagctgatcaaagtgattctggagatctatttc
gagttcaccagtcctgctatgctgttcctgtacggagggaacattaccccatggctgagatatgccgaatggctgctgacatgt
cccgtgatcctgattcatctgtctaacatcaccggcctgagtgaggcatacaataagcggacaatggctctgctggtgtccga
cctgggaactatttgcatgggagtgacagccgctctggccactgggtgggtgaagtggctgttttactgtatcggcctggtgta
tggaacccagacattctacaacgctggaatcatctacgtggagtcttactatatcatgcctgccggcggctgtaagaaactggt
gctggccatgactgccgtgtactattctagttggctgatgtttcccggcctgttcatctttgggcctgaaggcatgcacaccctg
agcgtggctgggtccactattggccataccatcgccgacctgctgtccaagaatatttggggactgctggggcacttcctgcg
gatcaaaattcacgagcatatcattatgtacggcgatatcaggagaccagtgagctcccagtttctgggacgcaaggtggac
gtgctggccttcgtgacagaggaagataaagtggcggccgccaagagctacccatacgatgttccagattacgcttaa

>Fig. 3a PEmax

atgaaacggacagccgacggaagcgagttcgagtcaccaaagaagaagcggaaagtcgacaagaagtacagcatcggc
ctggacatcggcaccaactctgtgggctgggccgtgatcaccgacgagtacaaggtgcccagcaagaaattcaaggtgctg
ggcaacaccgaccggcacagcatcaagaagaacctgatcggagccctgctgttcgacagcggcgaaacagccgaggcc
acccggctgaagagaaccgccagaagaagatacaccagacggaagaaccggatctgctatctgcaagagatcttcagcaa
cgagatggccaaggtggacgacagcttcttccacagactggaagagtccttcctggtggaagaggataagaagcacgagc
ggcaccccatcttcggcaacatcgtggacgaggtggcctaccacgagaagtaccccaccatctaccacctgagaaagaaa
ctggtggacagcaccgacaaggccgacctgcggctgatctatctggccctggcccacatgatcaagttccggggccacttc
ctgatcgagggcgacctgaaccccgacaacagcgacgtggacaagctgttcatccagctggtgcagacctacaaccagct
gttcgaggaaaaccccatcaacgccagcggcgtggacgccaaggccatcctgtctgccagactgagcaagagcagaaag
ctggaaaatctgatcgcccagctgcccggcgagaagaagaatggcctgttcggaaacctgattgccctgagcctgggcctg
acccccaacttcaagagcaacttcgacctggccgaggatgccaaactgcagctgagcaaggacacctacgacgacgacct
ggacaacctgctggcccagatcggcgaccagtacgccgacctgtttctggccgccaagaacctgtccgacgccatcctgct
gagcgacatcctgagagtgaacaccgagatcaccaaggcccccctgagcgcctctatgatcaagagatacgacgagcacc
accaggacctgaccctgctgaaagctctcgtgcggcagcagctgcctgagaagtacaaagagattttcttcgaccagagca
agaacggctacgccggctacattgacggcggagccagccaggaagagttctacaagttcatcaagcccatcctggaaaag
atggacggcaccgaggaactgctcgtgaagctgaagagagaggacctgctgcggaagcagcggaccttcgacaacggc
agcatccccccaccagatccacctgggagagctgcacgccattctgcggcggcaggaagatttttacccattcctgaaggaca
accgggaaaagatcgagaagatcctgaccttccgcatcccctactacgtgggccctctggccaggggaaacagcagattcg
cctggatgaccagaaagagcgaggaaaccatcaccccctggaacttcgaggaagtggtggacaagggcgcttccgccca
gagcttcatcgagcggatgaccaacttcgataagaacctgcccaacgagaaggtgctgcccaagcacagcctgctgtacga
gtacttcaccgtgtataacgagctgaccaaagtgaaatacgtgaccgagggaatgagaaagcccgccttcctgagcggcga
gcagaaaaaggccatcgtggacctgctgttcaagaccaaccggaaagtgaccgtgaagcagctgaaagaggactacttca
agaaaatcgagtgcttcgactccgtggaaatctccggcgtggaagatcggttcaacgcctccctgggcacataccacgatct
gctgaaaattatcaaggacaaggacttcctggacaatgaggaaaacgaggacattctggaagatatcgtgctgaccctgaca
ctgtttgaggacagagagatgatcgaggaacggctgaaaacctatgcccacctgttcgacgacaaagtgatgaagcagctg
aagcggcggagatacaccggctggggcaggctgagccggaagctgatcaacggcatccgggacaagcagtccggcaa
gacaatcctggatttcctgaagtccgacggcttcgccaacagaaacttcatgcagctgatccacgacgacagcctgaccttta
aagaggacatccagaaagcccaggtgtccggccagggcgatagcctgcacgagcacattgccaatctggccggcagccc
cgccattaagaagggcatcctgcagacagtgaaggtggtggacgagctcgtgaaagtgatgggccggcacaagcccgag
aacatcgtgatcgaaatggccagagagaaccagaccacccagaagggacagaagaacagccgcgagagaatgaagcg

gatcgaagagggcatcaaagagctgggcagccagatcctgaaagaacaccccgtggaaaacacccagctgcagaacga
gaagctgtacctgtactacctgcagaatgggcgggatatgtacgtggaccaggaactggacatcaaccggctgtccgacta
cgatgtggacgctatcgtgcctcagagctttctgaaggacgactccatcgacaacaaggtgctgaccagaagcgacaagaa
ccggggcaagagcgacaacgtgccctccgaagaggtcgtgaagaagatgaagaactactggcggcagctgctgaacgc
caagctgattacccagagaaagttcgacaatctgaccaaggccgagagaggcggcctgagcgaactggataaggccggc
ttcatcaagagacagctggtggaaacccggcagatcacaaagcacgtggcacagatcctggactcccggatgaacactaa
gtacgacgagaatgacaagctgatccgggaagtgaaagtgatcaccctgaagtccaagctggtgtccgatttccggaagga
tttccagttttacaaagtgcgcgagatcaacaactaccaccacgcccacgacgcctacctgaacgccgtcgtgggaaccgcc
ctgatcaaaaagtaccctaagctggaaagcgagttcgtgtacggcgactacaaggtgtacgacgtgcggaagatgatcgcc
aagagcgagcaggaaatcggcaaggctaccgccaagtacttcttctacagcaacatcatgaactttttcaagaccgagattac
cctggccaacggcgagatccggaagcggcctctgatcgagacaaacggcgaaaccggggagatcgtgtgggataaggg
ccgggattttgccaccgtgcggaaagtgctgagcatgccccaagtgaatatcgtgaaaaagaccgaggtgcagacaggcg
gcttcagcaaagagtctatcctgcccaagaggaacagcgataagctgatcgccagaaagaaggactgggaccctaagaag
tacggcggcttcgacagccccaccgtggcctattctgtgctggtggtggccaaagtggaaaagggcaagtccaagaaactg
aagagtgtgaaagagctgctggggatcaccatcatggaaagaagcagcttcgagaagaatcccatcgactttctggaagcc
aagggctacaaagaagtgaaaaaggacctgatcatcaagctgcctaagtactccctgttcgagctggaaaacggccggaag
agaatgctggcctctgccggcgaactgcagaagggaaacgaactggccctgccctccaaatatgtgaacttcctgtacctgg
ccagccactatgagaagctgaagggctcccccgaggataatgagcagaaacagctgtttgtggaacagcacaagcactac
ctggacgagatcatcgagcagatcagcgagttctccaagagagtgatcctggccgacgctaatctggacaaagtgctgtcc
gcctacaacaagcaccgggataagcccatcagagagcaggccgagaatatcatccacctgtttaccctgaccaatctggga
gcccctgccgccttcaagtactttgacaccaccatcgaccggaagaggtacaccagcaccaaagaggtgctggacgccac
cctgatccaccagagcatcaccggcctgtacgagacacggatcgacctgtctcagctgggaggtgactccggcggaagct
ctggtggcagcaagcggaccgccgacggctctgaattcgagagccctaagaagaaaagaaaggtgagcggaggctctag
cggcggaagcaccctgaacattgaagacgagtatagactgcatgaaacaagcaaggaacccgacgtgtccctgggctcca
cctggctgtccgactttccccaggcctgggccgagacaggaggaatgggcctggccgtgcggcaggcacccctgatcatc
cctctgaaggccacctctacacccgtgagcatcaagcagtaccctatgtctcaggaggccagactgggcatcaagcctcac
atccagaggctgctggaccagggcatcctggtgccatgccagagcccctggaacacaccactgctgcccgtgaagaagcc
aggcaccaatgactatagacccgtgcaggatctgagagaggtgaacaagagggtggaggatatccaccccaccgtgccca
acccttacaatctgctgtccggcctgcccccttctcaccagtggtatacagtgctggacctgaaggatgccttcttttgtctgag
actgcaccctaccagccagccactgttcgcctttgagtggagggaccctgagatgggcatctctggccagctgacctggaca

| |
|---|
| cgcctgcctcagggcttcaagaatagcccaacactgtttaacgaggccctgcaccgcgacctggcagatttccggatccagc<br>acccagatctgatcctgctgcagtacgtggacgatctgctgctggccgccaccagcgagctggattgccagcagggaacac<br>gcgccctgctgcagaccctgggaaacctgggatatagggcatccgccaagaaggcccagatctgtcagaagcaggtgaa<br>gtacctgggctatctgctgaaggagggccagagatggctgacagaggccaggaaggagacagtgatgggccagccaac<br>acccaagaccccaagacagctgagggagttcctgggcaaagcaggattttgcaggctgttcatcccaggattcgcagagat<br>ggcagcacctctgtacccactgaccaagccgggcaccctgtttaattggggccctgaccagcagaaggcctatcaggagat<br>caagcaggccctgctgacagcaccagccctgggcctgccagacctgaccaagcctttcgagctgtttgtggatgagaagca<br>gggctacgccaagggcgtgctgacccagaagctgggaccatggagacggcccgtggcctatctgtccaagaagctggac<br>ccagtggcagcaggatggccaccatgcctgaggatggtggcagcaatcgccgtgctgacaaaggatgccggcaagctga<br>ccatgggacagccactggtcatcctggcaccacacgcagtggaggccctggtgaagcagcctccagatcgctggctgtcta<br>acgcccggatgacacactaccaggccctgctgctggacaccgatcgcgtgcagtttggccctgtggtggccctgaatccag<br>ccaccctgctgcctctgccagaggagggcctgcagcacaactgtctggacatcctggcagaggcacacggaacaaggcc<br>agacctgaccgatcagcccctgcctgacgccgatcacacatggtataccgatggaagctccctgctgcaggagggccaga<br>ggaaggcaggagcagcagtgaccacagagacagaagtgatctgggccaaggccctgccagcaggcacatccgcccag<br>cgggccgagctgatcgccctgacccaggccctgaagatggccgagggcaagaagctgaacgtgtacacagactccagat<br>atgccttcgccaccgcacacatccacggagagatctacaggcgccggggctggctgacctctgagggcaaggagatcaa<br>gaacaaggatgagatcctggccctgctgaaggccctgtttctgcccaagcggctgagcatcatccactgtcctggacaccag<br>aagggacactccgccgaggcaaggggcaatcggatggccgaccaggccgccagaaaggctgctattactgaaactcccg<br>acacttccactctgctgattgaaaactcctccccttctggcggctcaaaaagaaccgccgacggcagcgaattcgagtctccc<br>aagaagaagaggaaagtcggctctggccctgccgctaagagagtgaagctggacGGAAGCGGATAA |
| >Fig. 3b hMLH1dn |
| ATGAGCTTCGTTGCTGGAGTCATCCGGAGACTGGACGAGACAGTGGTGAAC<br>AGAATTGCCGCCGGCGAGGTGATCCAGAGACCTGCCAATGCAATAAAGGAG<br>ATGATCGAGAACTGTCTGGACGCCAAGTCCACAAGCATTCAGGTGATCGTG<br>AAGGAGGGCGGACTGAAGCTGATCCAGATCCAAGACAACGGCACAGGCAT<br>CAGAAAGGAAGATCTGGACATCGTGTGTGAACGGTTCACCACATCTAAGCT<br>GCAGTCTTTTGAGGATCTGGCCTCTATCAGTACCTACGGCTTCAGAGGCGAG<br>GCCCTGGCCAGCATCAGCCACGTGGCCCATGTGACCATCACCACCAAAACC<br>GCCGACGGCAAATGCGCTTATCGCGCTAGCTACAGCGACGGCAAGCTGAAA |

GCCCCGCCAAAGCCTTGCGCCGGCAACCAGGGTACACAGATAACAGTGGA
GGATCTGTTCTACAACATCGCCACCCGGAGAAAGGCCCTGAAAAATCCCAG
CGAGGAGTACGGCAAGATCCTGGAAGTCGTCGGCAGATACTCCGTGCACAA
CGCCGGAATCAGCTTTAGCGTAAAGAAGCAGGGAGAAACCGTGGCCGATGT
GCGCACCCTGCCAAATGCCAGCACCGTGGATAACATCAGAAGCATTTTCGG
AAATGCCGTGTCCAGAGAACTGATCGAGATCGGCTGCGAAGATAAGACCCT
GGCTTTTAAGATGAACGGCTACATCAGCAACGCCAATTACTCTGTGAAGAAG
TGCATCTTTCTTCTGTTCATCAACCACAGACTGGTGGAAAGCACCAGCCTGC
GGAAAGCCATCGAGACAGTGTACGCCGCCTACCTGCCTAAGAACACCCACC
CCTTCCTGTACCTGAGCCTCGAGATCAGCCCTCAGAACGTGGACGTCAATGT
GCATCCTACAAAGCACGAGGTGCACTTCCTGCACGAGGAAAGCATCCTGGA
AAGAGTGCAGCAGCACATTGAGAGCAAGCTGCTGGGCTCTAACAGCAGCA
GAATGTACTTCACACAGACCCTGCTGCCTGGCCTGGCCGGCCCCTCAGGCG
AAATGGTTAAGTCCACAACCTCTCTGACCTCATCCAGCACCAGCGGTTCTTC
CGATAAGGTGTACGCCCACCAGATGGTGCGGACCGACTCTCGGGAGCAGAA
GCTGGACGCCTTTCTGCAACCTCTGAGCAAACCTCTGAGCTCTCAGCCTCA
GGCCATCGTGACCGAGGACAAGACAGATATCTCCTCCGGCCGTGCCAGACA
GCAGGACGAAGAAATGCTCGAGCTGCCAGCTCCTGCCGAGGTGGCCGCCA
AGAACCAGAGCCTGGAGGGAGATACCACAAAGGGCACCAGCGAAATGAGC
GAGAAGCGGGGCCCTACCTCCAGCAACCCCAGAAAACGGCACCGGGAGGA
CAGCGACGTGGAAATGGTGGAGGACGACAGCAGAAAGGAAATGACAGCCG
CTTGTACCCCTAGAAGAAGAATCATCAACCTGACCTCCGTGCTGAGCCTGCA
GGAGGAGATCAACGAGCAGGGCCACGAGGTGCTGAGAGAGATGCTGCACA
ATCACAGCTTCGTGGGCTGCGTGAACCCTCAATGGGCCCTGGCTCAGCATCA
AACAAAGCTGTACCTGCTGAACACCACCAAGCTGAGCGAAGAGCTGTTCTA
CCAGATCCTCATCTACGACTTCGCCAACTTCGGCGTGCTACGCCTGAGCGAG
CCCGCCCCTCTGTTTGACCTGGCCATGCTGGCTCTGGATAGCCCAGAAAGCG
GCTGGACAGAAGAGGACGGACCTAAAGAGGGGCTGGCTGAATACATCGTG
GAGTTCCTGAAGAAAAAGGCCGAGATGCTGGCCGACTACTTTTCTCTGGAA
ATCGACGAGGAAGGCAACCTGATCGGCCTGCCTCTGCTGATCGATAACTACG

| |
|---|
| TGCCTCCCCTGGAAGGCCTGCCCATCTTCATCCTGAGACTGGCTACAGAGGT<br>GAACTGGGACGAGGAAAAGGAATGCTTCGAGTCTCTGAGCAAGGAGTGCG<br>CCATGTTCTATAGCATCAGAAAACAGTACATCTCTGAAGAGAGCACTCTGTC<br>TGGCCAGCAGAGTGAAGTGCCCGGAAGCATCCCCAACAGCTGGAAGTGGA<br>CCGTGGAACACATCGTGTACAAGGCCCTGCGGAGCCACATTCTCCCTCCTAA<br>GCACTTCACCGAGGACGGCAACATCCTGCAGCTGGCCAACCTGCCCGACCT<br>TTATAAGGTTTTCTACCCATACGACGTCCCAGACTACGCTTAA |
| >Fig. 3f PEAR |
| atggtgagcaagggcgaggagctgttcaccggggtggtgcccatcctggtcgaactcgatggagatgtgaacggccacaa<br>gttcagcgtgtccggcgagggcgagggcgatgccacctacggcaagctgaccctgaagttcatctgcaccaccggcaagc<br>tgcccgtgccctggcccaccctcgtgaccaccctgacctacggcgtgcagtgcttcagccgctaccccgaccacatgaagc<br>agcacgacttcttcaagtccgccatgcccgaaggctacgtgcagacaagtccgtttacgtcgccgtccagctcattttttagtta<br>aaatatgggaaagtttaaaagaaggaatgatgaataaactatacaaccactttaaatatgtgttggtcccctttaagtcagcaag<br>cagatatatcagatatggcagctaacttaaggagtccccgtggatactggatgaatagcccccaattccaccaatgcaagaac<br>caacagtctatttggtcttgattaggacagtaaacttcctgcacaccatagtctgtgcactaaacaggagtaaaacaccccttaag<br>gcatctcttacattggctgcattaaagtgtgtcacttaatgacatccgtgatatcagggatagtttgacattggacatacagttctc<br>tctcagtgattgtcacctgcttatatagtacgaggacccatatgtggtcaatcactttcttgttgccactccatgccagggaagac<br>atagttacaatgagttataattataaaagctcagccttgcatgaggatataaaatacatttgccatacctctacatatgccttagttg<br>cttttaagctttgtttggaattgcaaaggccaacttcctaaacacagtttcaaagtattcttttgtcacacctttccactggatacttgt<br>attgtctgcaggaagaggccaccctgcctatgcttctccttagggcctcgttctgcagagtcagcataaactcccagtctgctc<br>aaccaattctgtctgattagggccatatctccttaccatgagttgatttttgtttctttttttgagcaggagcgcaccatcttcttcaag<br>gacgacggcaactacaagacccgcgccgaggtgaagttcgagggcgacaccctggtgaaccgcatcgagctgaagggc<br>atcgacttcaaggaggacggcaacatcctggggcacaagctggagtacaactacaacagccacaacgtctatatcatggcc<br>gacaagcagaagaacggcatcaaggtgaacttcaagatccgccacaacatcgaggacggcagcgtgcagctcgccgacc<br>actaccagcagaacacccccatcggcgacggccccgtgctgctgcccgacaaccactacctgagcacccagtccgccctg<br>agcaaagaccccaacgagaagcgcgatcacatggtcctgctggagttcgtgaccgccgccgggatcactctcggcatgga<br>cgagctgtacaagTAAgagggcctatttcccatgattccttcatatttgcatatacgatacaaggctgttagagagataattgg<br>aattaatttgactgtaaacacaaagatattagtacaaaatacgtgacgtagaaagtaataatttcttgggtagtttgcagttttaaa<br>attatgttttaaaatggactatcatatgcttaccgtaacttgaaagtatttcgatttcttggctttatatatcttgtggaaaggacgaaa |

| |
|---|
| caccgagctggacggcgacgtaaagttttagagctagaaatagcaagttaaaataaggctagtccgttatcaacttgaaaaagtggcaccgagtcggtgcggctacgtgcaggtaagtccgtttacgtcgccgtcctttttttccgttacataacttacggtaaatggcccaattcgagtggctccggtgcccgtcagtgggcagagcgcacatcgcccacagtc |

**Supplementary Table 5.** Sequences of epegRNAs and sgRNAs used in HEK293T cells.

| **Fig. 3k** | |
|---|---|
| RYR2$^{S404R}$-pegRNA-1 | GCATGAAGGCCACATGGATGAGTTTTAGAGCTAGAAATAGCAAGTTAAAATAAGGCTAGTCCGTTATCAACTTGAAAAAGTGGCACCGAGTCGGTGCGATCTCGACAAACgTATGCCATCATCCATGTGCGCGGTTCTATCTAGTTACGCGTTAAACCAACTAGAATTTTTT |
| RYR2$^{S404R}$-pegRNA-2 | GCATGAAGGCCACATGGATGAGTTTTAGAGCTAGAAATAGCAAGTTAAAATAAGGCTAGTCCGTTATCAACTTGAAAAAGTGGCACCGAGTCGGTGCGACAAACgTATGCCATCATCCATGTGCGCGGTTCTATCTAGTTACGCGTTAAACCAACTAGAATTTTTT |
| RYR2$^{S404R}$-pegRNA-3 | GCATGAAGGCCACATGGATGAGTTTTAGAGCTAGAAATAGCAAGTTAAAATAAGGCTAGTCCGTTATCAACTTGAAAAAGTGGCACCGAGTCGGTGCGGGATCTCGACAAACgTATGCCATCATCCATGTGGCCGCGGTTCTATCTAGTTACGCGTTAAACCAACTAGAATTTTTT |
| RYR2$^{S404R}$-pegRNA-4 | GCATGAAGGCCACATGGATGAGTTTTAGAGCTAGAAATAGCAAGTTAAAATAAGGCTAGTCCGTTATCAACTTGAAAAAGTGGCACCGAGTCGGTGCGACAAACgTATGCCATCATCCATGTGGCCGCGGTTCTATCTAGTTACGCGTTAAACCAACTAGAATTTTTT |
| | GCATGAAGGCCACATGGATGAGTTTTAGAGCTAGAAATAGCAAGTTAAAATAAGGCTAGTCCGTTATCAACTTG |

| | |
|---|---|
| RYR2$^{S404R}$-pegRNA-5 | AAAAAGTGGCACCGAGTCGGTGCGGGATCTCGACAAACgTATGCCATCATCCATGTGGCCTTCGCGGTTCTATCTAGTTACGCGTTAAACCAACTAGAATTTTTT |
| RYR2$^{S404R}$-pegRNA-6 | GCATGAAGGCCACATGGATGAGTTTTAGAGCTAGAAATAGCAAGTTAAAATAAGGCTAGTCCGTTATCAACTTGAAAAAGTGGCACCGAGTCGGTGCGACAAACgTATGCCATCATCCATGTGGCCTTCGCGGTTCTATCTAGTTACGCGTTAAACCAACTAGAATTTTTT |
| RYR2$^{S404R}$-pegRNA-7 | GCATGAAGGCCACATGGATGAGTTTTAGAGCTAGAAATAGCAAGTTAAAATAAGGCTAGTCCGTTATCAACTTGAAAAAGTGGCACCGAGTCGGTGCGGGATCTCGACAAACgTATGCCATCATCCATGTGGCCTTCACGCGGTTCTATCTAGTTACGCGTTAAACCAACTAGAATTTTTT |
| RYR2$^{S404R}$-pegRNA-8 | GCATGAAGGCCACATGGATGAGTTTTAGAGCTAGAAATAGCAAGTTAAAATAAGGCTAGTCCGTTATCAACTTGAAAAAGTGGCACCGAGTCGGTGCGACAAACgTATGCCATCATCCATGTGGCCTTCACGCGGTTCTATCTAGTTACGCGTTAAACCAACTAGAATTTTTT |
| RYR2$^{S404R}$-sgRNA | GAACgTATGCCATCATCCATGGTTTAGAGCTAGAAATAGCAAGTTAAAATAAGGCTAGTCCGTTATCAACTTG |
| **Fig. 3j** | |
| RYR2$^{E4950K}$-pegRNA-1 | GATGTATCAAGAAAGGTGTTGTTTTAGAGCTAGAAATAGCAAGTTAAAATAAGGCTAGTCCGTTATCAACTTGAAAAAGTGGCACCGAGTCGGTGCAAAAATTtCCAACACCTTTCTTGCGCGGTTCTATCTAGTTACGCGTTAAACCAACTAGAATTTTTT |

| | |
|---|---|
| RYR2$^{E4950K}$-pegRNA-2 | GATGTATCAAGAAAGGTGTTGTTTTAGAGCTAGAAATAGCAAGTTAAAATAAGGCTAGTCCGTTATCAACTTGAAAAAGTGGCACCGAGTCGGTGCGGAAAAATTtCCAACACCTTTCTTGCGCGGTTCTATCTAGTTACGCGTTAAACCAACTAGAATTTTTT |
| RYR2$^{E4950K}$-pegRNA-3 | GATGTATCAAGAAAGGTGTTGTTTTAGAGCTAGAAATAGCAAGTTAAAATAAGGCTAGTCCGTTATCAACTTGAAAAAGTGGCACCGAGTCGGTGCGCTGGGAAAAATTtCCAACACCTTTCTTGCGCGGTTCTATCTAGTTACGCGTTAAACCAACTAGAATTTTTT |
| RYR2$^{E4950K}$-pegRNA-4 | GATGTATCAAGAAAGGTGTTGTTTTAGAGCTAGAAATAGCAAGTTAAAATAAGGCTAGTCCGTTATCAACTTGAAAAAGTGGCACCGAGTCGGTGCAAAAATTtCCAACACCTTTCTTGATACGCGGTTCTATCTAGTTACGCGTTAAACCAACTAGAATTTTTT |
| RYR2$^{E4950K}$-pegRNA-5 | GATGTATCAAGAAAGGTGTTGTTTTAGAGCTAGAAATAGCAAGTTAAAATAAGGCTAGTCCGTTATCAACTTGAAAAAGTGGCACCGAGTCGGTGCGGAAAAATTtCCAACACCTTTCTTGATACGCGGTTCTATCTAGTTACGCGTTAAACCAACTAGAATTTTTT |
| RYR2$^{E4950K}$-pegRNA-6 | GATGTATCAAGAAAGGTGTTGTTTTAGAGCTAGAAATAGCAAGTTAAAATAAGGCTAGTCCGTTATCAACTTGAAAAAGTGGCACCGAGTCGGTGCGCTGGGAAAAATTtCCAACACCTTTCTTGATACGCGGTTCTATCTAGTTACGCGTTAAACCAACTAGAATTTTTT |
| | GATGTATCAAGAAAGGTGTTGTTTTAGAGCTAGAAATAGCAAGTTAAAATAAGGCTAGTCCGTTATCAACTTGAAAAAGTGGCACCGAGTCGGTGCAAAAATTtCCAACAC |

| | |
|---|---|
| RYR2$^{E4950K}$-pegRNA-7 | CTTTCTTGATACACGCGGTTCTATCTAGTTACGCGTTAAACCAACTAGAATTTTTT |
| RYR2$^{E4950K}$-pegRNA-8 | GATGTATCAAGAAAGGTGTTGTTTTAGAGCTAGAAATAGCAAGTTAAAATAAGGCTAGTCCGTTATCAACTTGAAAAAGTGGCACCGAGTCGGTGCGGAAAAATTtCCAACACCTTTCTTGATACACGCGGTTCTATCTAGTTACGCGTTAAACCAACTAGAATTTTTT |
| RYR2$^{E4950K}$-pegRNA-9 | GATGTATCAAGAAAGGTGTTGTTTTAGAGCTAGAAATAGCAAGTTAAAATAAGGCTAGTCCGTTATCAACTTGAAAAAGTGGCACCGAGTCGGTGCGCTGGGAAAAATTtCCAACACCTTTCTTGATACACGCGGTTCTATCTAGTTACGCGTTAAACCAACTAGAATTTTTT |
| RYR2$^{E4950K}$-sgRNA | GCTGGTCTTCATACTGTTTCGTTTTAGAGCTAGAAATAGCAAGTTAAAATAAGGCTAGTCCGTTATCAACTTGAAAAAGTGGCACCGAGTCGGTGCTTTTTT |
| **Fig. 3m** | |
| SCN5A$^{N406K}$-pegRNA-1 | GTTGCGACCACGGCCAGGATCGTTTTAGAGCTAGAAATAGCAAGTTAAAATAAGGCTAGTCCGTTATCAACTTGAAAAAGTGGCACCGAGTCGGTGCTGGTGAAaCTGATCCTGGCCGTGCGCGGTTCTATCTAGTTACGCGTTAAAC |
| SCN5A$^{N406K}$-pegRNA-2 | GTTGCGACCACGGCCAGGATCGTTTTAGAGCTAGAAATAGCAAGTTAAAATAAGGCTAGTCCGTTATCAACTTGAAAAAGTGGCACCGAGTCGGTGCACCTGGTGAAaCTGATCCTGGCCGTGCGCGGTTCTATCTAGTTACGCGTTAAACCAACTAGAATTTTTT |
| | GTTGCGACCACGGCCAGGATCGTTTTAGAGCTAGAAA |

| | |
|---|---|
| SCN5A$^{N406K}$-pegRNA-3 | TAGCAAGTTAAAATAAGGCTAGTCCGTTATCAACTTG AAAAAGTGGCACCGAGTCGGTGCTCTACCTGGTGAAa CTGATCCTGGCCGTGCGCGGTTCTATCTAGTTACGCGT TAAACCAACTAGAATTTTTT |
| SCN5A$^{N406K}$-pegRNA-4 | GTTGCGACCACGGCCAGGATCGTTTTAGAGCTAGAAA TAGCAAGTTAAAATAAGGCTAGTCCGTTATCAACTTG AAAAAGTGGCACCGAGTCGGTGCTGGTGAAaCTGATC CTGGCCGTGGTCCGCGGTTCTATCTAGTTACGCGTTAA ACCAACTAGAATTTTTT |
| SCN5A$^{N406K}$-pegRNA-5 | GTTGCGACCACGGCCAGGATCGTTTTAGAGCTAGAAA TAGCAAGTTAAAATAAGGCTAGTCCGTTATCAACTTG AAAAAGTGGCACCGAGTCGGTGCACCTGGTGAAaCTG ATCCTGGCCGTGGTCCGCGGTTCTATCTAGTTACGCGT TAAACCAACTAGAATTTTTT |
| SCN5A$^{N406K}$-pegRNA-6 | GTTGCGACCACGGCCAGGATCGTTTTAGAGCTAGAAA TAGCAAGTTAAAATAAGGCTAGTCCGTTATCAACTTG AAAAAGTGGCACCGAGTCGGTGCTCTACCTGGTGAAa CTGATCCTGGCCGTGGTCCGCGGTTCTATCTAGTTACG CGTTAAACCAACTAGAATTTTTT |
| SCN5A$^{N406K}$-pegRNA-7 | GTTGCGACCACGGCCAGGATCGTTTTAGAGCTAGAAA TAGCAAGTTAAAATAAGGCTAGTCCGTTATCAACTTG AAAAAGTGGCACCGAGTCGGTGCTGGTGAAaCTGATC CTGGCCGTGGTCGCCGCGGTTCTATCTAGTTACGCGTT AAACCAACTAGAATTTTTT |
| SCN5A$^{N406K}$-pegRNA-8 | GTTGCGACCACGGCCAGGATCGTTTTAGAGCTAGAAA TAGCAAGTTAAAATAAGGCTAGTCCGTTATCAACTTG AAAAAGTGGCACCGAGTCGGTGCACCTGGTGAAaCTG ATCCTGGCCGTGGTCGCCGCGGTTCTATCTAGTTACGC |

| | |
|---|---|
| | GTTAAACCAACTAGAATTTTTT |
| SCN5A$^{N406K}$-pegRNA-9 | GTTGCGACCACGGCCAGGATCGTTTTAGAGCTAGAAATAGCAAGTTAAAATAAGGCTAGTCCGTTATCAACTTGAAAAAGTGGCACCGAGTCGGTGCTCTACCTGGTGAAaCTGATCCTGGCCGTGGTCGCCGCGGTTCTATCTAGTTACGCGTTAAACCAACTAGAATTTTTT |
| SCN5A$^{N406K}$-sgRNA | GCTACCTGGTGAAaCTGATCCGTTTTAGAGCTAGAAATAGCAAGTTAAAATAAGGCTAGTCCGTTATCAACTTGAAAAAGTGGCACCGAGTCGGTGCTTTTTT |
| **Fig. 3n** | |
| KCNQ1$^{A341V}$-pegRNA-1 | GCCCACGGGGCACCTACCGCTGTTTTAGAGCTAGAAATAGCAAGTTAAAATAAGGCTAGTCCGTTATCAACTTGAAAAAGTGGCACCGAGTCGGTGCTTCTTTGtGctgCCAGCGGTAGGTGCCCGCGGTTCTATCTAGTTACGCGTTAAACCAACTAGAATTTTTT |
| KCNQ1$^{A341V}$-pegRNA-2 | GCCCACGGGGCACCTACCGCTGTTTTAGAGCTAGAAATAGCAAGTTAAAATAAGGCTAGTCCGTTATCAACTTGAAAAAGTGGCACCGAGTCGGTGCTCCTTCTTTGtGctgCCAGCGGTAGGTGCCCGCGGTTCTATCTAGTTACGCGTTAAACCAACTAGAATTTTTT |
| KCNQ1$^{A341V}$-pegRNA-3 | GCCCACGGGGCACCTACCGCTGTTTTAGAGCTAGAAATAGCAAGTTAAAATAAGGCTAGTCCGTTATCAACTTGAAAAAGTGGCACCGAGTCGGTGCATCTCCTTCTTTGtGctgCCAGCGGTAGGTGCCCGCGGTTCTATCTAGTTACGCGTTAAACCAACTAGAATTTTTT |
| | GCCCACGGGGCACCTACCGCTGTTTTAGAGCTAGAAA |

| | |
|---|---|
| KCNQ1$^{A341V}$-pegRNA-4 | TAGCAAGTTAAAATAAGGCTAGTCCGTTATCAACTTGAAAAAGTGGCACCGAGTCGGTGCGCCATCTCCTTCTTTGtGctgCCAGCGGTAGGTGCCCGCGGTTCTATCTAGTTACGCGTTAAACCAACTAGAATTTTTT |
| KCNQ1$^{A341V}$-pegRNA-5 | GCCCACGGGGCACCTACCGCTGTTTTAGAGCTAGAAATAGCAAGTTAAAATAAGGCTAGTCCGTTATCAACTTGAAAAAGTGGCACCGAGTCGGTGCTTCTTTGtGctgCCAGCGGTAGGTGCCCCGCGCGGTTCTATCTAGTTACGCGTTAAACCAACTAGAATTTTTT |
| KCNQ1$^{A341V}$-pegRNA-6 | GCCCACGGGGCACCTACCGCTGTTTTAGAGCTAGAAATAGCAAGTTAAAATAAGGCTAGTCCGTTATCAACTTGAAAAAGTGGCACCGAGTCGGTGCTCCTTCTTTGtGctgCCAGCGGTAGGTGCCCCGCGCGGTTCTATCTAGTTACGCGTTAAACCAACTAGAATTTTTT |
| KCNQ1$^{A341V}$-pegRNA-7 | GCCCACGGGGCACCTACCGCTGTTTTAGAGCTAGAAATAGCAAGTTAAAATAAGGCTAGTCCGTTATCAACTTGAAAAAGTGGCACCGAGTCGGTGCATCTCCTTCTTTGtGctgCCAGCGGTAGGTGCCCCGCGCGGTTCTATCTAGTTACGCGTTAAACCAACTAGAATTTTTT |
| KCNQ1$^{A341V}$-pegRNA-8 | GCCCACGGGGCACCTACCGCTGTTTTAGAGCTAGAAATAGCAAGTTAAAATAAGGCTAGTCCGTTATCAACTTGAAAAAGTGGCACCGAGTCGGTGCGCCATCTCCTTCTTTGtGctgCCAGCGGTAGGTGCCCCGCGCGGTTCTATCTAGTTACGCGTTAAACCAACTAGAATTTTTT |
| KCNQ1$^{A341V}$-pegRNA-9 | GCCCACGGGGCACCTACCGCTGTTTTAGAGCTAGAAATAGCAAGTTAAAATAAGGCTAGTCCGTTATCAACTTGAAAAAGTGGCACCGAGTCGGTGCTTCTTTGtGctgCCAGCGGTAGGTGCCCCGTGCGCGGTTCTATCTAGTTACGC |

| | |
|---|---|
| | GTTAAACCAACTAGAATTTTTT |
| KCNQ1$^{A341V}$-pegRNA-10 | GCCCACGGGGCACCTACCGCTGTTTTAGAGCTAGAAATAGCAAGTTAAAATAAGGCTAGTCCGTTATCAACTTGAAAAAGTGGCACCGAGTCGGTGCTCCTTCTTTGtGctgCCAGCGGTAGGTGCCCCGTGCGCGGTTCTATCTAGTTACGCGTTAAACCAACTAGAATTTTTT |
| KCNQ1$^{A341V}$-pegRNA-11 | GCCCACGGGGCACCTACCGCTGTTTTAGAGCTAGAAATAGCAAGTTAAAATAAGGCTAGTCCGTTATCAACTTGAAAAAGTGGCACCGAGTCGGTGCATCTCCTTCTTTGtGctgCCAGCGGTAGGTGCCCCGTGCGCGGTTCTATCTAGTTACGCGTTAAACCAACTAGAATTTTTT |
| KCNQ1$^{A341V}$-pegRNA-12 | GCCCACGGGGCACCTACCGCTGTTTTAGAGCTAGAAATAGCAAGTTAAAATAAGGCTAGTCCGTTATCAACTTGAAAAAGTGGCACCGAGTCGGTGCGCCATCTCCTTCTTTGtGctgCCAGCGGTAGGTGCCCCGTGCGCGGTTCTATCTAGTTACGCGTTAAACCAACTAGAATTTTTT |
| KCNQ1$^{A341V}$-sgRNA | GCTCCTTCTTTGtGctgCCAGGTTTTAGAGCTAGAAATAGCAAGTTAAAATAAGGCTAGTCCGTTATCAACTTGAAAAAGTGGCACCGAGTCGGTGCTTTTTT |
| **Fig. 6c** | |
| TRPM4$^{A320V}$-pegRNA-1 | GCCTGGCTCCCCCACTCCCTGGTTTTAGAGCTAGAAATAGCAAGTTAAAATAAGGCTAGTCCGTTATCAACTTGAAAAAGTGGCACCGAGTCGGTGCACTCTGGTCCCAGGGAGTGGGGGCGCGGTTCTATCTAGTTACGCGTTAAACCAACTAGAATTTTTT |
| | GCCTGGCTCCCCCACTCCCTGGTTTTAGAGCTAGAAA |

| | |
|---|---|
| TRPM4$^{A320V}$-pegRNA-2 | TAGCAAGTTAAAATAAGGCTAGTCCGTTATCAACTTGAAAAAGTGGCACCGAGTCGGTGCGACACTCTGGTCCCAGGGAGTGGGGGCGCGGTTCTATCTAGTTACGCGTTAAACCAACTAGAATTTTTT |
| TRPM4$^{A320V}$-pegRNA-3 | GCCTGGCTCCCCCACTCCCTGGTTTTAGAGCTAGAAATAGCAAGTTAAAATAAGGCTAGTCCGTTATCAACTTGAAAAAGTGGCACCGAGTCGGTGCGAAGACACTCTGGTCCCAGGGAGTGGGGGCGCGGTTCTATCTAGTTACGCGTTAAACCAACTAGAATTTTTT |
| TRPM4$^{A320V}$-pegRNA-4 | GCCTGGCTCCCCCACTCCCTGGTTTTAGAGCTAGAAATAGCAAGTTAAAATAAGGCTAGTCCGTTATCAACTTGAAAAAGTGGCACCGAGTCGGTGCACTCTGGTCCCAGGGAGTGGGGGAGCCGCGGTTCTATCTAGTTACGCGTTAAACCAACTAGAATTTTTT |
| TRPM4$^{A320V}$-pegRNA-5 | GCCTGGCTCCCCCACTCCCTGGTTTTAGAGCTAGAAATAGCAAGTTAAAATAAGGCTAGTCCGTTATCAACTTGAAAAAGTGGCACCGAGTCGGTGCGACACTCTGGTCCCAGGGAGTGGGGGAGCCGCGGTTCTATCTAGTTACGCGTTAAACCAACTAGAATTTTTT |
| TRPM4$^{A320V}$-pegRNA-6 | GCCTGGCTCCCCCACTCCCTGGTTTTAGAGCTAGAAATAGCAAGTTAAAATAAGGCTAGTCCGTTATCAACTTGAAAAAGTGGCACCGAGTCGGTGCGAAGACACTCTGGTCCCAGGGAGTGGGGGAGCCGCGGTTCTATCTAGTTACGCGTTAAACCAACTAGAATTTTTT |
| TRPM4$^{A320V}$-pegRNA-7 | GCCTGGCTCCCCCACTCCCTGGTTTTAGAGCTAGAAATAGCAAGTTAAAATAAGGCTAGTCCGTTATCAACTTGAAAAAGTGGCACCGAGTCGGTGCACTCTGGTCCCAGGGAGTGGGGGAGCCACGCGGTTCTATCTAGTTACGCGT |

| | |
|---|---|
| | TAAACCAACTAGAATTTTTT |
| TRPM4$^{A320V}$-pegRNA-8 | GCCTGGCTCCCCCACTCCCTGGTTTTAGAGCTAGAAATAGCAAGTTAAAATAAGGCTAGTCCGTTATCAACTTGAAAAAGTGGCACCGAGTCGGTGCGACACTCTGGTCCCAGGGAGTGGGGGAGCCACGCGGTTCTATCTAGTTACGCGTTAAACCAACTAGAATTTTTT |
| TRPM4$^{A320V}$-pegRNA-9 | GCCTGGCTCCCCCACTCCCTGGTTTTAGAGCTAGAAATAGCAAGTTAAAATAAGGCTAGTCCGTTATCAACTTGAAAAAGTGGCACCGAGTCGGTGCGAAGACACTCTGGTCCCAGGGAGTGGGGGAGCCACGCGGTTCTATCTAGTTACGCGTTAAACCAACTAGAATTTTTT |
| TRPM4$^{A320V}$-sgRNA | GCTGGAAGACACTCTGGTCCCGTTTTAGAGCTAGAAATAGCAAGTTAAAATAAGGCTAGTCCGTTATCAACTTGAAAAAGTGGCACCGAGTCGGTGCTTTTTT |

**Supplementary Table 6.** Primer sequences used for amplicon amplification.

| | **Sequence 5'~3'** |
|---|---|
| **Fig. 3k,l and 4d** | |
| RYR2$^{S404R}$-F | GAGACGTTGGGAGTAATGGCC |
| RYR2$^{S404R}$-R | TGAAGTATCAAACAACTACG |
| **Fig. 3j and l** | |
| RYR2$^{E4950K}$-F | CCGTGCGATATAGGTAACAG |
| RYR2$^{E4950K}$-R | TTAGAGGTGATTGGGTCTGAG |
| **Fig. 3m and o** | |

| | |
|---|---|
| $SCN5A^{N406K}$-F | CCTCCTGGAAGCGCTTTTCC |
| $SCN5A^{N406K}$-R | TCAGGTCCGCAGGGAAGATC |
| **Fig. 3n and o** | |
| $KCNQ1^{A341V}$-F | AGTCACCACCATCGGCTATG |
| $KCNQ1^{A341V}$-R | CGTAAGTGGGTCTGCTCACAG |
| **Fig. 4d** | |
| $DNMT1^{\Delta 1\_15}$-F | CGTGTTCCCCAGAGTGAC |
| $DNMT1^{\Delta 1\_15}$-R | CATGTACCACACATGTGAACG |
| **Fig. 6c and d** | |
| $TRPM4^{A320V}$-F | AGGCTCAGCTCCCATGTCTC |
| $TRPM4^{A320V}$-R | GCCTGCAGGACCTCAAGGTC |

**Supplementary Table 7.** Genomic loci analyzed by amplicon sequencing.

| | |
|---|---|
| $RYR2^{S404R}$ | GAGACGTTGGGAGTAATGGCCTTATTTTGCTTTCTTACAGGCTATTATGCATCATGAAGGCCACATGGATGATGGCATAAGTTTGTCGAGATCCCAGCATGAAGAATCACGCACAGCCCGAGTTATCCGGAGCACAGTCTTCCTTTTCAATAGATTTATAAGGTACTTTTTCTTTTGTAGGCGTAGTTGTTTGATACTTCA |
| $RYR2^{E4950K}$ | CCGTGCGATATAGGTAACAGAATTGAGTTTCTACCTTATGTTTTGTTAGCACACACTTTGGGGAAAATGTTAATACATTTCCTTGACTTTTGCAGGAATCTTATGTCTGGAAGATGTATCAAGAAAGGTGTTGGGAATTTTTCCCAGCAGGGGATTGCTTCCGGAAACAGTATGAAGACCAGCTAAATTAAACTCAGACCCAATCACCTCTAA |

| | |
|---|---|
| SCN5A$^{N406K}$ | CCTCCTGGAAGCGCTTTTCCTTCTCCTCGGTCTCAGCGATGGTGGCTTGGTTTTGCTCCTCATAGGCCATTGCGACCACGGCCAGGATCAGGTTCACCAGGTAGAAGGACCCCAGGAAGATGACAAGCATGAAGAAGATCATGTAGATCTTCCCTGCGGACCTGA |
| KCNQ1$^{A341V}$ | AGTCACCACCATCGGCTATGGGGACAAGGTGCCCCAGACGTGGGTCGGGAAGACCATCGCCTCCTGCTTCTCTGTCTTTGCCATCTCCTTCTTTGCGCTCCCAGCGGTAGGTGCCCCGTGGGTGCGTTTTCCCTGGCTCCTTGGACAGCTGGGGTCCTGGGGTGGCTGCACGCCCCTCCCTGTGAGCAGACCCACTTACG |
| TRPM4$^{A320V}$ | AGGCTCAGCTCCCATGTCTCCTCGTGGCTGGCTCAGGGGGAGCTGCGGACTGCCTGGCGGAGACCCTGGAAGACACTCTGGCCCCAGGGAGTGGGGGAGCCAGGCAAGGCGAAGCCCGAGATCGAATCAGGCGTTTCTTTCCCAAAGGGGACCTTGAGGTCCTGCAGGC |
| DNMT1$^{\Delta 1_15}$ | CGTGTTCCCCAGAGTGACTTTTCCTTTTATTTCCCTTCAGCTAAAATAAAGGAGGAGGAAGCTGCTAAGGACTAGTTCTGCCCTCCCGTCACCCCTGTTTCTGGCACCAGGAATCCCCAACATGCACTGATGTTGTGTTTTTAACATGTCAATCTGTCCGTTCACATGTGTGGTACATG |

**Supplementary Table 8.** Templates used to synthesize sgRNAs for SSB-ddPCR.

| | **Sequence 5'~3'** |
|---|---|
| SCN5A$^{N406K}$-ddPCR | TTCTAATACGACTCACTATAGCTACCTGGTGAACCTGATCCGTTTTAGAGCTAGA |
| RYR2$^{E4950K}$-ddPCR | TTCTAATACGACTCACTATAGAGCAGGGGATTGCTTCCGGAGTTTTAGAGCTAGA |
| TRPM4$^{A320V}$-ddPCR | TTCTAATACGACTCACTATAGGAGTGTCTTCCAGGGTCTCCGTTTTAGAGCTAGA |

**Supplementary Table 9.** Primer and probe sequences used for SSB-ddPCR.

| **Fig. 5d and e** | **Sequence 5'~3'** |
|---|---|
| SCN5A$^{N406K}$-FOR1 | CGCAGGGAAGATCTACATGA |
| SCN5A$^{N406K}$-REV1 | GCTTGGTTTTGCTCCTCATA |
| SCN5A$^{N406K}$-FOR2 | TTCTGGTGGGGTAGTGAAGA |
| SCN5A$^{N406K}$-REV2 | GCTCACTCTATAGCCAGCCT |
| SCN5A$^{N406K}$-Probe-FAM | TCTACCTGGTGAACCTGATCCTGGCC |
| SCN5A$^{N406K}$-Probe-HEX | TAGCCATCCCTAGTCCCCCAC |
| SCN5A$^{N406K}$-Probe-FAM | /56-FAM/TCTACCTGG/ZEN/TGAACCTGATCCTGGCC/3IABkFQ/ |
| SCN5A$^{N406K}$-Probe-HEX | /5HEX/TAGCCATCC/ZEN/CTAGTCCCCCAC/3IABkFQ/ |
| **Extended Data Fig. 4a** | |
| RyR2$^{E4950K}$-FOR1 | AGAGGTGATTGGGTCTGAGTT |
| RyR2$^{E4950K}$-REV1 | ACATTTCCTTGACTTTTGCAGG |
| RyR2$^{E4950K}$-FOR2 | AGAAAACGCCACATGCAAAT |
| RyR2$^{E4950K}$-REV2 | AGCCTCTAGGGTCAAATAGCA |
| RyR2$^{E4950K}$-Probe-FAM | /56-FAM/ATCCCCTGC/ZEN/TGGGAAAAATTCCCA/3IABkFQ/ |
| RyR2$^{E4950K}$-Probe-HEX | /5HEX/ACCAGAAAG/ZEN/AGGAAAGGCTGTGGAC/3IABk |

| | |
|---|---|
| | FQ/ |
| **Extended Data Fig. 4b** | |
| TRPM4$^{A320V}$-FOR1 | CGAATAGAGAACGCCACCC |
| TRPM4$^{A320V}$-REV1 | CTGATTCGATCTCGGGCTTC |
| TRPM4$^{A320V}$-FOR2 | TCATCCACCCTCTCTAATCCA |
| TRPM4$^{A320V}$-REV2 | GGAAAGCAGATGCTGGAACATA |
| TRPM4$^{A320V}$-Probe-FAM | /56-FAM/AGACCCTGG/ZEN/AAGACACTCTGGCC/3IABkFQ/ |
| TRPM4$^{A320V}$-Probe-HEX | /5HEX/ATGGTGCTA/ZEN/TTTGGGCAGTTTTCCC/3IABkFQ/ |
